# Programmable large DNA deletion, replacement, integration, and inversion with twin prime editing and site-specific recombinases

**DOI:** 10.1101/2021.11.01.466790

**Authors:** Andrew V. Anzalone, Xin D. Gao, Christopher J. Podracky, Andrew T. Nelson, Luke W. Koblan, Aditya Raguram, Jonathan M. Levy, Jaron A. M. Mercer, David R. Liu

## Abstract

The targeted deletion, replacement, integration, or inversion of DNA sequences at specified locations in the genome could be used to study or treat many human genetic diseases. Here, we describe twin prime editing (twinPE), a method for the programmable replacement or excision of DNA sequence at endogenous human genomic sites without requiring double-strand DNA breaks. TwinPE uses a prime editor (PE) protein and two prime editing guide RNAs (pegRNAs) that template the synthesis of complementary DNA flaps on opposing strands of genomic DNA, resulting in the replacement of endogenous DNA sequence between the PE-induced nick sites with pegRNA-encoded sequences. We show that twinPE in human cells can perform precise deletions of at least 780 bp and precise replacements of genomic DNA sequence with new sequences of at least 108 bp. By combining single or multiplexed twinPE with site-specific serine recombinases either in separate steps or in a single step, we demonstrate targeted integration of gene-sized DNA plasmids (>5,000 bp) into safe-harbor loci including *AAVS1*, *CCR5,* and *ALB* in human cells. To our knowledge, these results represent the first RNA-programmable insertion of gene-sized DNA sequences into targeted genomic sites of unmodified human cells without requiring double-strand breaks or homology-directed repair. Twin PE combined with recombinases also mediated a 40,167-bp inversion at *IDS* that corrects a common Hunter syndrome allele. TwinPE expands the capabilities of precision gene editing without requiring double-strand DNA breaks and synergizes with other tools to enable the correction or complementation of large or complex pathogenic alleles in human cells.

---

Disease-associated human genetic variants arise through a variety of sequence changes, ranging from single-base pair substitutions to mega base-scale duplications, deletions and rearrangements^1–3^. Gene editing approaches that can install, correct, or complement these pathogenic variants in human cells have the potential to advance our understanding of human genetic disease and could also lead to the development of new therapeutics^4, 5^. Several mammalian cell gene editing approaches based on CRISPR-Cas systems have been developed over the past decade^6^, including nucleases^7–9^, base editors^10, 11^, and prime editors^12^, each of which has the potential to address a subset of known pathogenic sequence changes.

CRISPR-Cas nucleases such as Cas9 can be used to disrupt genes through the generation of double-strand DNA breaks (DSBs) that lead to stochastic indels. In addition, paired Cas9 nuclease strategies have been developed for the targeted deletion of genomic DNA sequences ranging from ∼50 to >100,000 base pairs in length^13^. By providing a linear donor DNA sequence, targeted insertion of new DNA sequences can be performed at single cut sites or between paired cut sites through end-joining or homology-directed repair (HDR) processes^14, 15^.

While versatile in some ways, single-nuclease and paired-nuclease editing approaches have substantial drawbacks. HDR is inefficient compared to the generation of uncontrolled mixtures of indels in the vast majority of cell types^16, 17^. Likewise, the use of paired nucleases for targeted deletion also generates multiple byproducts, including the desired deletion accompanied by undesired indels, indels at individual DSB sites without the desired deletion, undesired inversion of the intervening sequence between DSB sites, and unintended integration of exogenous DNA sequence at DSB sites^13, 18^. In addition, the precise location of the deletions is restricted by PAM availability, which dictates the location of nuclease-mediated DNA cutting. Similar restrictions and byproducts exist for DNA donor knock-in, which typically occurs without control of sequence orientation when homology-independent approaches are used^14^ and is accompanied by efficient indel byproduct generation^19^. Finally, the simultaneous generation of multiple DSBs at on-target or off-target sites can promote large deletions^20–22^, chromosomal rearrangements such as translocations^23^, and chromothripsis^24^. The tendency of DSBs to generate large mixtures of undesired byproducts and chromosomal changes^25–27^ poses considerable challenges when applying nuclease-based editing strategies for the excision, replacement, or insertion of larger DNA sequences, especially in therapeutic settings.

Base editors can be used to precisely install C•G-to-T•A^10^ or A•T-to-G•C^11^ transition mutations, or C•G-to-G•C transversion mutations^28–30^ without requiring DSBs^6^. Prime editors can precisely install any of the twelve possible base pair substitutions as well as small insertions and deletions without requiring DSBs^12^. In their original form, however, these DSB-free editing technologies cannot be directly used to efficiently install edits that alter hundreds or thousands of base pairs, such as large deletions, gene-sized insertions, large replacements, or large inversions.

Prime editing has been shown to be capable of making precise insertions of up to ∼40 bp and deletions of up to ∼80 bp with PE2 and PE3 systems in human cells with high ratios of desired edits to byproducts^12^. While prime editing offers the flexibility to replace one DNA segment with a new sequence, PE2 and PE3 systems have not yet been able to mediate larger insertions and deletions of the size of typical exons or gene coding sequences, and the presumed mechanism of simple prime editing reactions makes these larger DNA changes difficult by requiring long pegRNA reverse transcription templates and long-range DNA polymerization. In contrast, site-specific DNA recombinases have the ability to perform DNA excision, inversion, integration, or exchange of large DNA sequences in mammalian cells^31, 32^. The longstanding challenge of reprogramming site-specific recombinases^33–36^, however, has limited their use for precision gene editing applications.

Here, we report the development of twin prime editing (twinPE), which enables the deletion, substitution, or insertion of larger DNA sequences at targeted endogenous genomic sites with high efficiencies in human cells. In addition to these capabilities, twinPE can also be used to integrate one or more recombinase recognition sites with high efficiency at targeted sites in the human genome. These recognition sites can then be used as substrates for site-specific serine recombinases to enable the targeted integration of gene-sized DNA segments, or the targeted inversion of gene-sized genomic DNA sequences. The twinPE and recombinase reagents can be delivered either stepwise, or simultaneously to enable one-step targeted integration or inversion of large DNA segments at endogenous genomic loci.

## Results

### Twin prime editing strategy

Prime editing uses a prime editor protein comprising a fusion of a catalytically impaired Cas9 nickase and a wild-type (PE1) or engineered (PE2) MMLV reverse transcriptase (RT) enzyme, and a prime editing guide RNA (pegRNA) that both specifies the target genomic site and encodes the desired edit^12^. Upon target site recognition, PE•pegRNA complexes nick the PAM-containing DNA strand and directly reverse transcribe the pegRNA’s RT template into genomic DNA using the nicked strand as a primer. Following reverse transcription, the newly synthesized 3’ flap containing the edited sequence invades the adjacent DNA to replace the redundant 5’ flap sequence. The opposing nonedited strand is then repaired using the edited DNA strand as a template. This proposed prime editing pathway presents at least two opportunities for cellular DNA repair to reject the desired edit and revert the DNA sequence to its original form: during 3’ flap annealing and ligation, and during heteroduplex resolution.

We hypothesized that bypassing potentially disfavored steps in DNA repair might allow prime editing to occur with increased efficiency and enable new classes of targeted gene edits in mammalian cells. We envisioned a twin prime editing (twinPE) strategy that uses a pair of pegRNAs, each of which targets a different DNA strand and templates the synthesis of a 3’ flap that is complementary to the 3’ flap templated by the other pegRNA (Fig. 1a). We hypothesized that if the newly synthesized DNA strands were highly dissimilar to the endogenous target site, the complementary 3’ flaps would preferentially hybridize with each other to create an intermediate containing annealed 3’ overhangs of new DNA sequence and annealed 5’ overhangs of original DNA sequence (Fig. 1a). As both edited strands are synthesized by prime editor complexes, there is no requirement for invasion of the target site by edit-containing flap strands, or for the edit to be copied to the complementary DNA strand. Excision of the original DNA sequence (annealed 5’ overhangs), filling in of gaps by polymerases, and ligation of the pair of nicks would result in the replacement of the endogenous sequence between the nick sites with the paired 3’ flap sequences (Fig. 1a). Due to the flexibility in template design, the edit could in principle insert a new DNA sequence, delete a portion of original DNA sequence, or replace a portion of original DNA.

**Figure 1.**
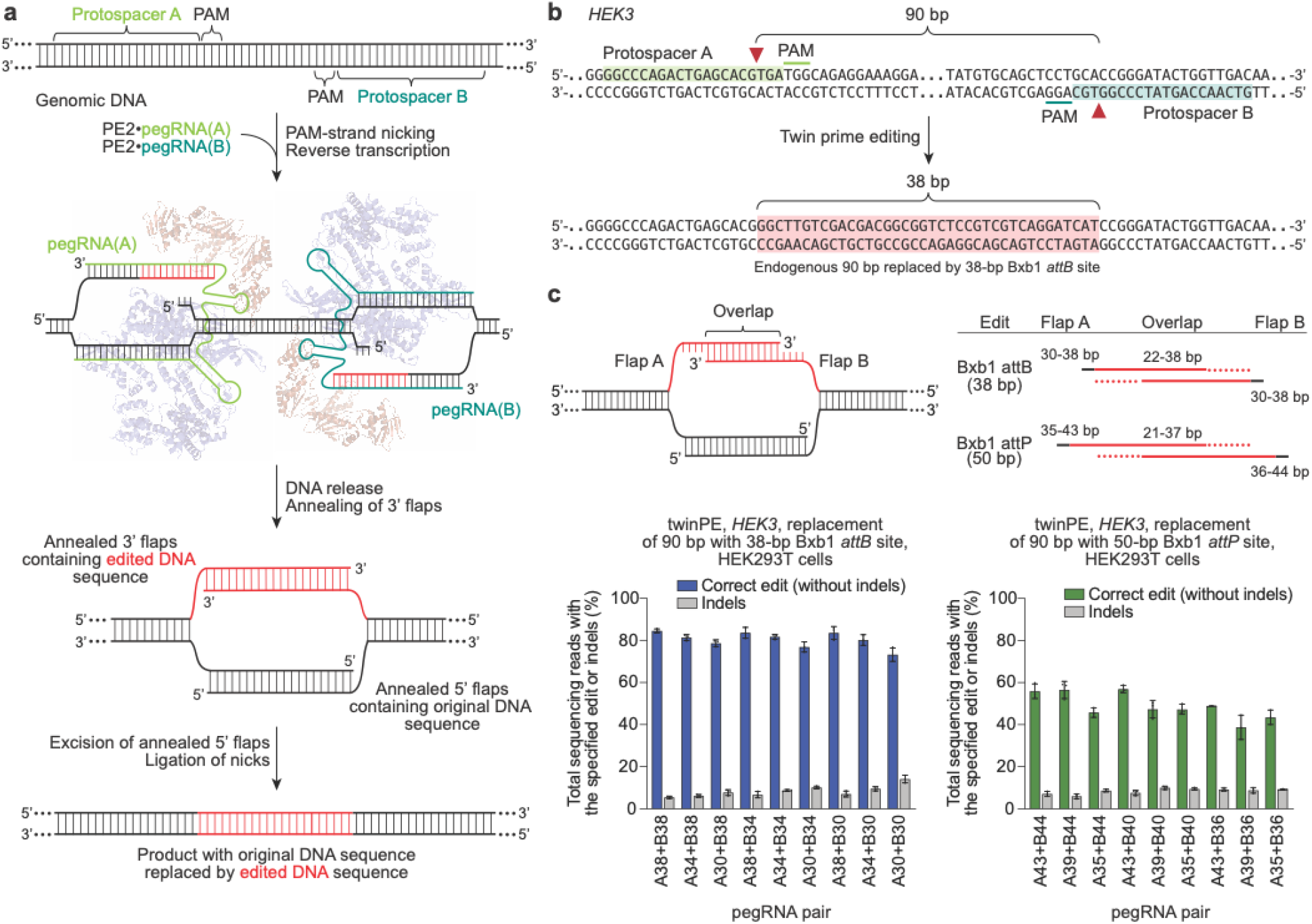
Overview of twinPE and twinPE-mediated sequence replacement. (**a**) TwinPE systems target genomic DNA sequences that contain two protospacer sequences on opposite strands of DNA. PE2•pegRNA complexes target each protospacer, generate a single-stranded nick, and reverse transcribe the pegRNA-encoded template containing the desired insertion sequence. After synthesis and release of the 3’ DNA flaps, a hypothetical intermediate exists possessing annealed 3’ flaps containing the edited DNA sequence and annealed 5’ flaps containing the original DNA sequence. Excision of the original DNA sequence contained in the 5’ flaps, followed by ligation of the 3’ flaps to the corresponding excision sites, generates the desired edited product. (**b**) Example of twinPE-mediated replacement of a 90-bp sequence in *HEK3* with a 38-bp Bxb1 *attB* sequence. (**c**) Evaluation of twinPE in HEK293T cells for the installation of the 38-bp Bxb1 *attB* site as shown in (b) or the 50-bp Bxb1 *attP* site at *HEK3* using pegRNAs that template varying lengths of the insertion sequence. pegRNA names indicate spacer (A or B) and length of RT template. Values and error bars reflect the mean and s.d. of three independent biological replicates.

### TwinPE-mediated large sequence replacement and deletion

To test the twinPE strategy, we initially targeted the HEK293T site 3 locus (hereafter referred to as *HEK3*) in HEK293T cells to replace 90 bp of endogenous sequence with a 38-bp Bxb1 recombinase *attB* substrate sequence^37^ (Fig. 1b). For each protospacer, three pegRNAs were designed with RT templates that contained 30, 34, or 38 nt of the 38-bp *attB* sequence (Fig. 1c). Pairwise combinations of these pegRNAs are predicted to generate 3’ flaps with overlapping complementarity ranging from 22 to 38 bp (Fig. 1c). When both pegRNAs were transfected into HEK293T cells along with PE2, we observed highly efficient *attB* site insertion, with some combinations of pegRNAs yielding >80% conversion to the desired product (Fig. 1c). A similar strategy for the replacement of the 90-bp endogenous sequence with the 50-bp Bxb1 *attP* attachment sequence also achieved high editing efficiencies up to 58% (Fig. 1c). Notably, it was not necessary for either pegRNA to encode the full insertion sequence, since partially overlapping complementary flaps enabled full-length *attB* or *attP* sequence incorporation. However, 3’ flaps with greater overlap led to slightly higher editing efficiencies and fewer indels for insertion of the Bxb1 *attB* and *attP* sequences at *HEK3* (Fig. 1c).

Independent of our efforts, Shendure and co-workers^38^ and Gao and co-workers^39^ reported elegant dual-pegRNA prime editing systems that can be used to perform precise large deletions (up to 710 bp) in human cells or improve base substitution and small insertion or deletion edits in plant cells, respectively. For each of these strategies, the templated 3’ flap sequence is homologous to the target site DNA sequence to facilitate DNA repair for incorporation of the edit (Fig. 2a). In contrast, twinPE was designed to bypass the need for any homologous DNA sequence in the pegRNA RT template, thus offering more template sequence flexibility and the freedom to mediate larger insertions.

**Figure 2.**
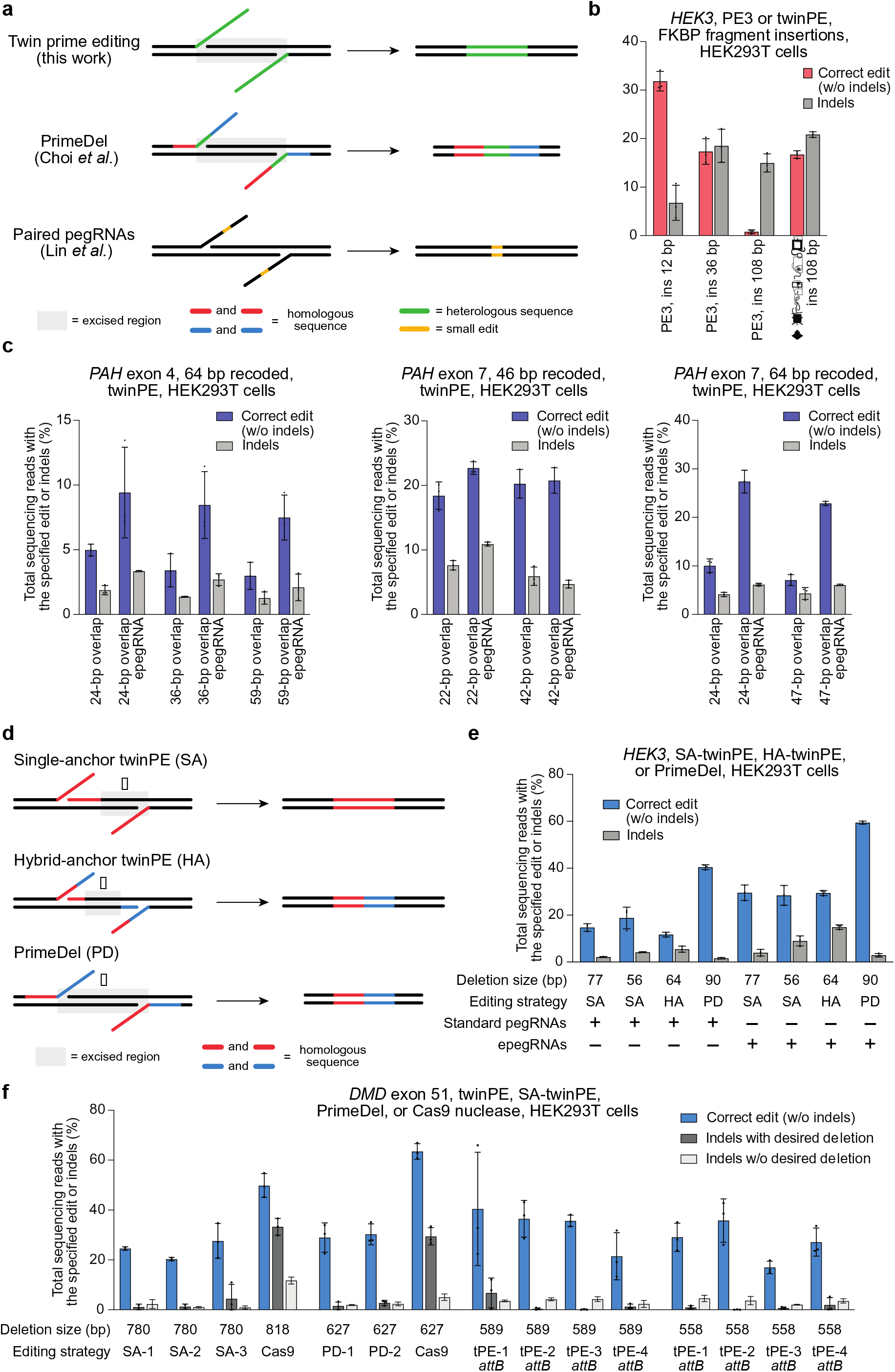
Targeted sequence insertion, deletion, and recoding with twinPE in human cells. (**a**) Schematic diagram illustrating designs and edits generated for twinPE, PrimeDel^36^ and paired pegRNAs^37^. Shaded gray boxes indicate regions where DNA is excised, green lines indicate the incorporation of heterologous DNA sequence, red and blue lines indicate regions of sequence homology (red to red, blue to blue), and yellow lines indicate regions with small edits. PrimeDel can introduce but does not require introduction of heterologous sequence (green). (**b**) Insertion of FKBP coding sequence fragments with PE3 (12 bp, 36 bp, 108 bp) or twinPE (108 bp) at *HEK3* in HEK293T cells. (**c**) Recoding of sequence within exons 4 and 7 in *PAH* in HEK293T cells using twinPE. A 64-bp target sequence in exon 4 was edited using 24, 36, or 59 bp of overlapping flaps, a 46-bp target sequence in exon 7 was edited using 22 or 42 bp of overlapping flaps, or a 64-bp sequence in exon 7 was edited using 24 or 47 bp of overlapping flaps. Editing activity was compared using standard pegRNAs or epegRNAs containing 3’ evoPreQ1 motifs. (**d**) Schematic diagram showing three distinct dual-flap deletion strategies that were investigated for carrying out targeted deletions. The “Single-anchor (SA)” twinPE strategy allows for flexible deletion starting at an arbitrary position 3’ of one nick site and ending at the other nick site. The “Hybrid-anchor (HA)” twinPE strategy allows for flexible deletion of sequence at arbitrarily chosen positions between the two nick sites. The “PrimeDel (PD)” strategy of Shendure and co-workers tested here allows for deletion of the sequence starting at one nick site and ending at another nick site. Shaded gray boxes indicate regions where DNA is excised, red and blue lines indicate regions of sequence homology (red to red, blue to blue). (**e**) Deletion of sequences at *HEK3* in HEK293T cells using the SA-twinPE, HA-twinPE, or PD strategies targeting the same protospacer pair. Editing activity was compared using standard pegRNAs or epegRNAs containing 3’ evoPreQ1 motifs. (**f**) Deletion of exon 51 sequence at the *DMD* locus in HEK293T cells using SA-twinPE, PD, paired Cas9 nuclease, or twinPE-mediated *attB* sequence replacement. A unique molecular identifier (UMI) protocol was applied to remove PCR bias (see **Supplementary Note 1**). Values and error bars in (b-f) reflect the mean and s.d. of three independent biological replicates.

Next, we tested if twinPE could support the insertion of DNA sequences larger than the ≤44-bp insertions previously demonstrated^12^ using PE2 or PE3 prime editing systems. First, we examined PE3-mediated insertions of *FKBP12* coding fragments ranging from 12 to 108 bp. Targeting the *HEK3* locus with PE3 in HEK293T cells, we achieved moderate editing efficiencies for the shorter 12-bp and 36-bp insertions (32% and 17%, respectively), but very little desired product for the 108-bp insertion (0.80%) (Fig. 2b). By contrast, twinPE enabled 16% insertion efficiency for the 108-bp fragment with concomitant deletion of 90 bp of *HEK3* sequence, a 20-fold improvement over a the 108-bp insertion with PE3. Similarly, twinPE was 2- to 4-fold more efficient than PE3 for inserting a 108-bp *FKBP12* cDNA fragment at *CCR5* (Extended Data Fig. 1). Finally, 113-bp and 103-bp insertions containing pairs of Bxb1 recombinase sites (*attB*–27 bp spacer–*attP* or *attB*–27 bp spacer–*attB*) were installed with similar efficiencies (11% and 9.7%, respectively) at *CCR5* using twinPE (Extended Data Fig. 1). These results demonstrate the ability of twinPE to mediate larger insertions than have been previously demonstrated with PE2 and PE3 systems.

One potential application of the twinPE strategy is the replacement of exonic coding sequence with recoded DNA sequence that maintains the protein sequence and has the potential to correct any combination of mutations between target twinPE-induced nick sites. Such a capability raises the possibility of using a single pegRNA set and prime editor to correct any of a variety of pathogenic mutations within a stretch of DNA, potentially addressing multiple cohorts of animal models or patients. To test this approach, we targeted *PAH*, encoding phenylalanine hydroxylase. Mutations within *PAH* cause the genetic metabolic disorder phenylketonuria (PKU)^40^. We tested the ability of twin PE to recode portions of *PAH* exon 4 and exon 7, which commonly harbor mutations in PKU patients, in HEK293T cells. By testing different flap overlap lengths using engineered pegRNAs (epegRNAs) containing a 3’ evoPreQ1 motif^41^, we achieved the desired sequence recoding with average efficiencies of up to 23% for a 46-bp recoding and up to 27% for a longer overlapping 64-bp recoding in exon 7, and up to 9.4% average efficiency for a 64-bp recoding of exon 4 (Fig. 2c and Extended Data Fig. 2). Additional exons in *PAH* could also be recoded, albeit with lower efficiency (Extended Data Fig. 2). These results demonstrate that twinPE can replace stretches of dozens of nucleotides in human cells with a single pair of pegRNAs.

In addition to insertion and replacement of DNA sequences, twinPE may also mediate precise deletions more effectively than previously described methods. Paired-nuclease deletion strategies generate deletions of sequence between the two DSBs and are thus restricted by PAM availability. In addition, desired deletions are accompanied by undesired indel and inversion byproducts. In contrast, twinPE has the potential to generate deletions with greater flexibility and precision due to the lack of required DSBs and the ability to write additional DNA sequence at the twinPE-induced nick sites. To assess twinPE for precise deletions, we compared three strategies using paired pegRNAs: a “single-anchor” (SA) twinPE strategy, which fixes the deletion junction at one of the two nick sites; a “hybrid-anchor” (HA) twinPE strategy, which allows flexible deletion junctions between the nick sites; and, the PrimeDel (PD) strategy recently reported by Shendure and coworkers^38^, which generates deletions between the nick sites with the option of inserting additional DNA sequence (Fig. 2d). Each strategy differs in the relationship of the sequences of the two flaps and thus in the positioning of the deletion with respect to the nick sites (Fig. 2d).

Targeting the *HEK3* site in HEK293T cells and using the single-anchor strategy, 13-nt complementary flaps were used to delete 77 bp of sequence adjacent to one of the pegRNA-induced nick sites with 15% efficiency and 2.1% indels (Fig. 2e, SA-Δ77), and 34-nt complementary flaps were used to precisely excise 56 bp of sequence with 19% efficiency and 4.2% indels (Fig. 2e, SA-Δ56). Using the hybrid-anchor strategy, we deleted 64 bp between the pegRNA-induced nick sites such that the product retains 13 bp of sequence 3’ of each nick with 12% efficiency and 5.5% indels (Fig. 2e, HA-Δ64). Finally, we tested the PrimeDel strategy for the deletion of 90 bp between the pegRNA-induced nick sites, which occurred with 40% efficiency and 1.6% indels (Fig. 2e, PD-Δ90). The higher efficiency of the PrimeDel strategy may arise from its ability to disrupt the PAM sequences on both strands, which likely increases deletion efficiency by limiting target re-engagement that can lead to indels or re-nicking of the prime-edited strand. Editing efficiencies could be improved for all three approaches by 1.5-fold to 2.5-fold upon the addition of the evoPreQ1 motif the 3’ end of the resulting epegRNAs^41^ (Fig. 2e). Together, these data show that twinPE offers a strategy for performing targeted deletions with high flexibility and high product purity that does not rely on the availability of nuclease cut sites.

In a final application of twinPE to mediate targeted deletions, we targeted a therapeutically relevant locus, *DMD*. Pathogenic *DMD* alleles, which are responsible for Duchenne muscular dystrophy, commonly contain large deletions in regions containing exons that result in frameshifted mRNA transcripts^42^. Because production of nearly full-length dystrophin protein without replacement of deleted exons can lead to partial rescue of protein function^43^, deletion of a second exon to restore the reading frame has been proposed as a potential therapeutic strategy^44^. We examined three twinPE deletion strategies along with a previously reported Cas9 nuclease deletion strategy for excising exon 51 in *DMD*^45^. Using single-anchor twinPE deletion approaches, we observed 28% average efficiency for the deletion of a 780-bp sequence containing exon 51 in *DMD* (Fig. 2f). While the paired sgRNA Cas9 nuclease strategy for deleting 818 bp achieved higher deletion efficiency (averaging 50% precise deletion) compared to twinPE strategies, paired Cas9 nuclease-mediated deletion was also accompanied by much higher indel levels of 45% (33% desired deletion with additional indels, plus 12% indels without the desired deletion) compared to twinPE, which averaged 4% total indels (Fig. 2f).

Exon 51 of *DMD* could also be excised using alternative spacer pairs that generate a 627-bp deletion with the PrimeDel strategy (30% average efficiency) or by using twinPE to replace a 589-bp sequence with a 38-bp Bxb1 *attB* sequence (up to 40% average efficiency) (Fig. 2f). These experiments show that twinPE and PrimeDel are capable of generating large deletions at therapeutically relevant loci in human cells with far fewer indel byproducts compared to paired Cas9 nuclease strategies.

Both standard prime editing and twinPE require complementarity between the pegRNA spacer and target DNA sequence for Cas9 targeting and nicking, complementarity between the pegRNA PBS and the nicked genomic DNA primer for initiation of reverse transcription, and complementarity between the newly synthesized 3’ DNA flap and a local DNA sequence (adjacent DNA for standard PE, the second 3’ flap for twinPE) for productive prime editing. These three DNA hybridization requirements may explain the observation of reduced off-target prime editing compared to that of Cas9 nuclease reported by our group and others^12, 46–52^. To examine potential off-target editing by twinPE, we amplified and sequenced four previously characterized off-target loci for one of the *HEK3* spacer sequences following treatment by eight different sets of twinPE reagents targeting *HEK3* (SA-Δ77, SA-Δ56, HA-Δ64, and PD-Δ90 with standard pegRNAs and epegRNAs). For each of the eight twinPE treatments, no detectable off-target edits or indels were detected at any of the four off-target sites beyond background levels in untreated control samples (Supplementary Table 5).

### Targeted DNA integration at safe harbor loci with twinPE and Bxb1 integrase

Although twinPE can perform larger insertion edits than PE3, the upper limit of sequence insertion with twinPE is insufficient to integrate gene-sized DNA fragments of thousands of base pairs. Having successfully inserted Bxb1 recombinase substrate sequences into endogenous human genomic sites with high efficiency using twinPE, we sought to combine twinPE with serine recombinases for the site-specific integration of DNA cargo (Fig. 3a). Researchers have previously used Bxb1 recombinase to integrate exogenous DNA ranging in size from single genes to entire genetic circuits into genomically-integrated *attB* and *attP* DNA sequences in a variety of cultured mammalian cells^53, 54^ and in *Drosophila*^55^. To identify sites for DNA cargo integration, we first tested twinPE-mediated insertion of Bxb1 *attB* and *attP* attachment sequences at established human genome safe harbor loci in HEK293T cells. We screened 19 spacer pairs targeting the *CCR5* locus for insertion of the 38-bp Bxb1 *attB* sequence (Extended Data Fig. 3). Optimal pegRNAs for six spacer pairs achieved >50% editing efficiency of perfectly edited alleles with 3.9–5.4% indel byproducts (Fig. 3b). Likewise, 32 pegRNA pairs targeting the *AAVS1* locus were screened for insertion of the 50-bp Bxb1 *attP* sequence (Extended Data Fig. 4), of which 17 optimal spacer combinations achieved >50% efficiency of perfectly edited alleles with a median of 6.6% indels (Fig. 3c). Notably, twinPE outperformed PE3 for the replacement of endogenous sequence at *CCR5* in HEK293T cells with Bxb1 recombinase *attB* sites. TwinPE yielded 62% perfectly edited alleles with 3.3% indels at one *CCR5* site, and 67% perfectly edited alleles with 4.3% indels at a second *CCR5* site, compared to 3.3% perfectly edited alleles with 0.1% indels at the first site and 25% perfectly edited alleles with 1.8% indels at the second site by PE3 (Extended Data Fig. 5). These results demonstrate that twinPE can be used to insert recombinase substrate sequences at safe harbor loci in human cells with high efficiency.

**Figure 3.**
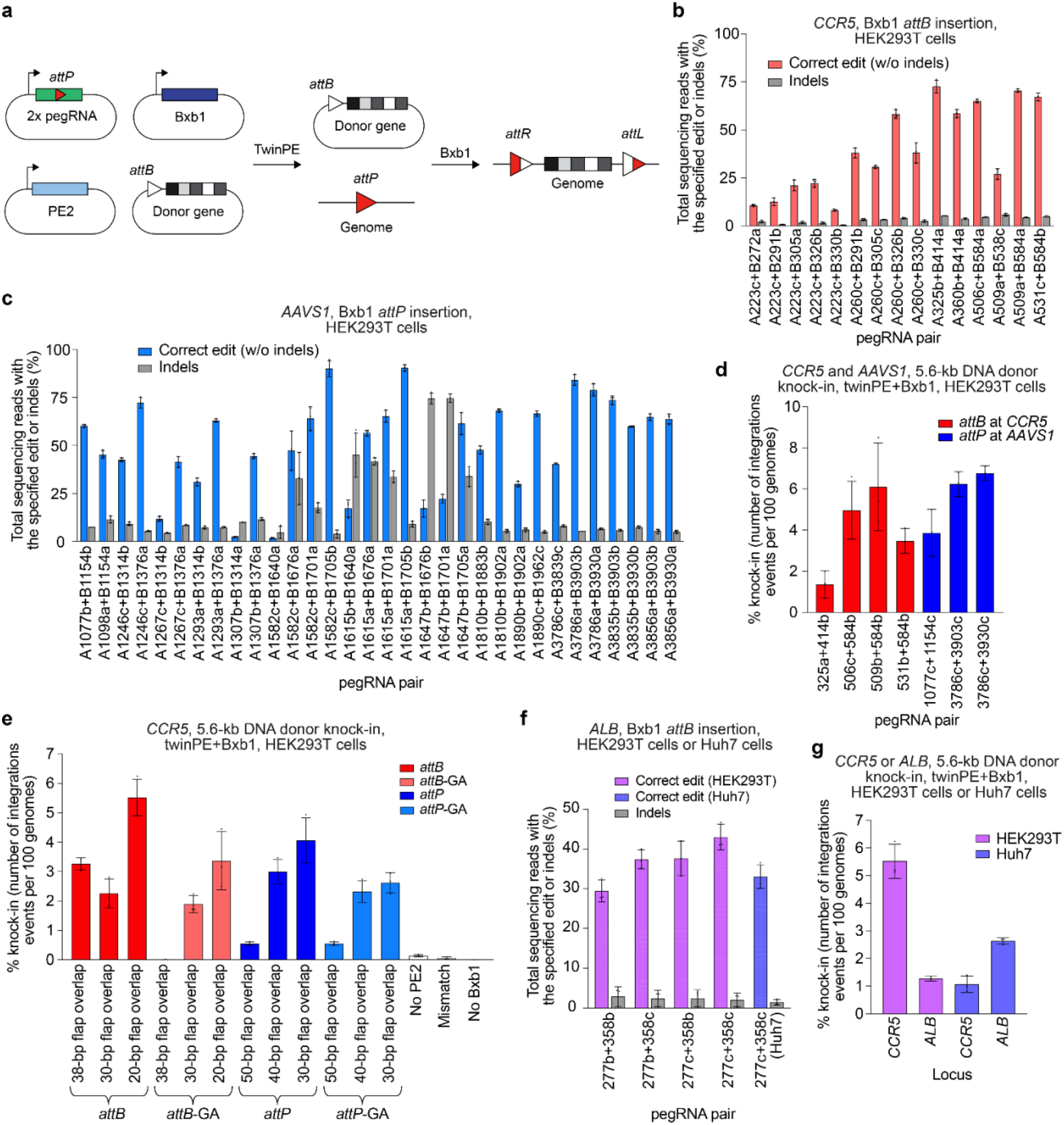
Site-specific genomic integration of DNA cargo with twinPE and Bxb1 recombinase in human cells. (**a**) Schematic diagram of twinPE and Bxb1 recombinase-mediated site-specific genomic integration of DNA cargo. (**b**) Screening of twinPE pegRNA pairs for insertion of the Bxb1 *attB* sequence at the *CCR5* locus in HEK293T cells. (**c**) Screening of twinPE pegRNA pairs for installation of the Bxb1 *attP* sequence at the *AAVS1* locus in HEK293T cells. (**d**) Single transfection knock-in of 5.6-kb DNA donors using twinPE pegRNA pairs targeting *CCR5* (red) or *AAVS1* (blue). The twinPE pegRNAs install *attB* at *CCR5* or *attP* at *AAVS1*. Bxb1 integrates a donor bearing the corresponding attachment site into the genomic attachment site. The number of integration events per 100 genomes is defined as the ratio of the target amplicon spanning the donor-genome junction to a reference amplicon in *ACTB*, as determined by ddPCR. (**e**) Optimization of single-transfection integration at *CCR5* using the A531+B584 spacers for the twinPE pegRNA pair. Identity of the templated edit (*attB* or *attP*), identity of the central dinucleotide (wild-type GT or orthogonal mutant GA), and length of the overlap between flaps were varied to identify combinations that supported the highest integration efficiency. % knock-in quantified as in (d). (**f**) Pairs of pegRNAs were assessed for their ability to insert Bxb1 *attB* into the first intron of *ALB*. Protospacer sequences (277 and 358) are constant across the pegRNA pairs. The pegRNAs vary in their PBS lengths (variant b or c). The 277c/358c pair that performs best in HEK293T cells can also introduce the desired edit in Huh7 cells. (**g**) Comparison of single transfection knock-in efficiencies at *CCR5* and *ALB* in HEK293T and Huh7 cell lines. % knock-in quantified as in (d). Values and error bars reflect the mean and s.d. of three independent biological replicates.

Next, we examined if twinPE-incorporated *attB* or *attP* sequences could serve as target substrates for the BxB1-mediated integration of DNA plasmids containing partner *attP* or *attB* sequences. First, we used twinPE to generate single-cell HEK293T clones bearing homozygous *attB* site insertions at the *CCR5* locus. Transfection of this clonal HEK293T cell line with a plasmid expressing Bxb1 recombinase and a 5.6-kB *attP*-containing donor DNA plasmid yielded an average of 12-17% integration events per genome of the 5.6-kB plasmid at the target *CCR5* site as measured by ddPCR and comparison with an *ACTB* reference (Extended Data Fig. 6). This efficiency is consistent with previously reported Bxb1-mediated plasmid integration efficiencies in mammalian cells^56^.

Encouraged by these results, we explored whether twinPE-mediated recombinase site insertion and Bxb1 recombinase-mediated DNA donor integration could be achieved by delivering all necessary components in a single transfection step. Encouragingly, our initial efforts to transfect HEK293T cells with plasmids encoding PE2, both pegRNAs, BxB1, and a 5.6-kB donor plasmid resulted in 1.4–6.8% knock-in efficiency as measured by ddPCR (Fig. 3d). The anticipated junction sequences containing the expected *attL* and *attR* recombination products were confirmed by amplicon sequencing, with very high product purities ranging from 91.6-99.8% average *attL* or *attR* junctions without indels (median = 99.0%) (Extended Data Fig. 6).

In an effort to improve one-step plasmid integration efficiency with twinPE and Bxb1 recombinase, we tested the twinPE-mediated incorporation of either *attB* or *attP* attachment sites, Bxb1 attachment sites containing the canonical GT central dinucleotide core sequence or an alternative GA dinucleotide core sequence^56^, and pegRNAs encoding varying flap overlap lengths. We found that one-step twinPE and Bxb1-mediated plasmid integration performed better when integrating the *attB* site compared to the *attP* site, especially when pegRNAs encoded the full-length *attB* or *attP* sequence (3.3% vs. 0.5% desired integration, Fig. 3e). In addition, we observed that integrations with *attB* and *attP* attachment sites containing the canonical GT core sequence generally led to higher knock-in efficiencies compared to those using the alternative GA core sequences (Fig. 3e). Notably, attachment sites containing canonical and alternative core sequences are orthogonal to one another, allowing simultaneous knock-in of an *attB-*GT donor at an *attP-*GT site in *AAVS1 a*nd GA dinucleotide donors at GA attachment sites in *CCR5* (Extended Data Fig. 6).

Reducing the extent of flap overlap between pegRNA-encoded recombinase sequences also improved single-transfection plasmid integration efficiency from 3.3±0.2% with 38 base pairs of overlap to 5.5±0.6% with 20 base pairs of overlap when inserting *attB*, or from 0.55±0.1% with 50 base pairs of overlap to 4.1±0.8% with 30 base pairs of overlap when inserting *attP* (Fig. 3e). Similar twinPE editing efficiencies were observed for inserting *attB* sequences with long (38-bp) or short (20-bp) flap overlaps. However, products of recombination between *attP*-containing donor DNA and pegRNA-encoding plasmid were observed when using the 38-bp overlapping pegRNA pair but were not detected when using the 20-bp flap overlap pegRNA pair (Extended Data Fig. 6). Therefore, enhanced integration efficiency with partially overlapping flaps may arise from reducing or eliminating usage of the pegRNA-expressing plasmid DNA as a substrate for Bxb1 recombinase.

We next evaluated plasmid integration at the *ALB* locus, which is being used clinically for therapeutic transgene expression in hepatocytes^57–59^. As a result of high albumin production by the liver, therapeutic transgene integration at the *ALB* locus in a relatively small percentage of hepatocytes in principle could achieve therapeutic levels of protein expression for many loss-of-function diseases^57^. In the case of Factor IX transgene expression, increasing circulating protein levels to just 1% of normal levels has shown a therapeutic effect for the treatment of hemophilia^60^. We devised a twinPE-enabled strategy targeting intron 1 of *ALB* in which Bxb1 recombinase mediates the integration of circular DNA containing a splice acceptor sequence followed by the cDNA for a protein of interest. Following integration, splicing of the albumin secretion signal encoded in exon 1 to the therapeutic transgene coding sequence would enable its expression and secretion.

We screened pegRNA pairs for the twinPE-mediated insertion of *attB* within intron 1 of *ALB* (Fig. 3f) and identified a spacer combination that achieved 43% correct insertion of the *attB* sequence. Single-transfection integration of a mock donor plasmid (a promoter-less copy of EGFP and PuroR-T2A-BFP under control of the EF1α promoter; 5.6-kB total) at this locus was achieved with 1.3% efficiency in HEK293T cells (Fig. 3g). In Huh7 cells, we observed 34% correct insertion of the *attB* sequence (Fig. 3f) and achieved single transfection knock-in efficiency of 2.6%. Interestingly, knock-in at *ALB* in Huh7 cells was more efficient than knock-in at *CCR5* (2.6% vs. 1.1%), despite *CCR5* knock-in being more efficient than *ALB* knock-in in HEK293T cells (Fig. 3g). The transfection of Huh7 cells with plasmids encoding Bxb1, PE2, an *attP*-containing donor harboring a splice acceptor followed by the cDNA for human factor IX (hFIX) exons 2-8, and pegRNAs programming the insertion of Bxb1 *attB* at intron 1 of *ALB* led to detectable levels of hFIX in conditioned media, whereas no hFIX was detected when the pegRNAs targeted *CCR5* instead of *ALB* (Extended Data Fig. 7). Collectively, these results establish a new method for the insertion of gene-sized DNA sequences into targeted genomic loci in previously unmodified human cells without double strand breaks or HDR.

To assess the possibility of unintended Bxb1-mediated donor integration or sequence modifications at endogenous off-target sites in the human genome, we studied five Bxb1 pseudo-sites for further characterization. Pseudo-sites were selected based on the presence of a minimal Bxb1 recognition motif identified by high-resolution recombinase specificity profiling (ACNACNGNNNNNNCNGTNGT) that is common to both *attP* and *attB* substrates^61^. Additionally, candidate pseudo-sites required a central GT dinucleotide to match that of the donor DNA plasmid used in our experiments, as corresponding central dinucleotides are necessary for recombination between *attB* and *attP*^62^. To search for undesired sequence modifications, we performed targeted amplicon sequencing for each of the five nominated pseudo-sites from seven different samples that had been treated with a 5.6-kb donor DNA plasmid, twinPE reagents targeting *CCR5* or *AAVS1*, and Bxb1 recombinase. For all pseudo-sites amplified from treated samples, we observed undetectable levels of indels (<0.1%) or near-background levels of indels as compared to untreated controls (Extended Data Fig. 7). To capture donor DNA plasmid integration events at the nominated pseudo-sites, we attempted to PCR amplify predicted integration junctions from treated genomic DNA samples. For each sample and pseudo-site target, we observed no PCR product corresponding to off-target donor integration, while amplicons reporting on the presence of on-target donor integration were readily observed in all cases (Extended Data Fig. 7). These results demonstrate that twinPE+Bxb1 can mediate precise donor integration at the intended target site without generating detected levels of undesired donor integration or sequence alteration at predicted Bxb1 pseudo-sites in the human genome.

### TwinPE and recombinase-mediated large inversions

Large structural variants are found in many human pathogenic alleles, such as the large 600-kb inversion at the *F8* locus that causes ∼50% of severe hemophilia A cases^63^. Inspired by the high efficiency of recombinase attachments site insertions using twinPE, we reasoned that multiplexing the insertion of both *attB* and *attP* Bxb1 attachment sites could be used to correct more complex genetic variants by unidirectional deletion or inversion of the intervening DNA sequence. We first tested whether twinPE and Bxb1 can revert an inverted H2B-EGFP coding sequence that is stably integrated into the HEK293T genome via lentivirus transduction (Extended Data Fig. 8). After transfection of the reporter cells with twinPE and Bxb1, we observe up to 19% GFP positive cells by flow cytometry, indicating successful inversion (Extended Data Fig. 8).

To test the combined twinPE-recombinase inversion approach on a therapeutically relevant locus, we performed a 40-kb inversion between *IDS* and its pseudogene *IDS2*. Inversions between these sites have been observed in ∼13% of Hunter syndrome patients^64^, and characterization of the breakpoints in pathogenic alleles revealed that the inversion often occurs within a recombination hotspot that is present in both *IDS* and *IDS2*^64^. We targeted regions flanking these recombination hotspots to insert *attB* and *attP* sequences, which when oriented in opposing directions can be used as substrates for unidirectional inversion by Bxb1 recombinase to correct the pathogenic allele (Fig. 4a). After screening pegRNA spacer pairs and optimizing pegRNA sequence features, we found pegRNA pairs that are capable of inserting *attB* or *attP* attachment sites at the left and right targeted sites with 70% and 74% efficiency, respectively in HEK293T cells (Fig. 4b).

**Figure 4.**
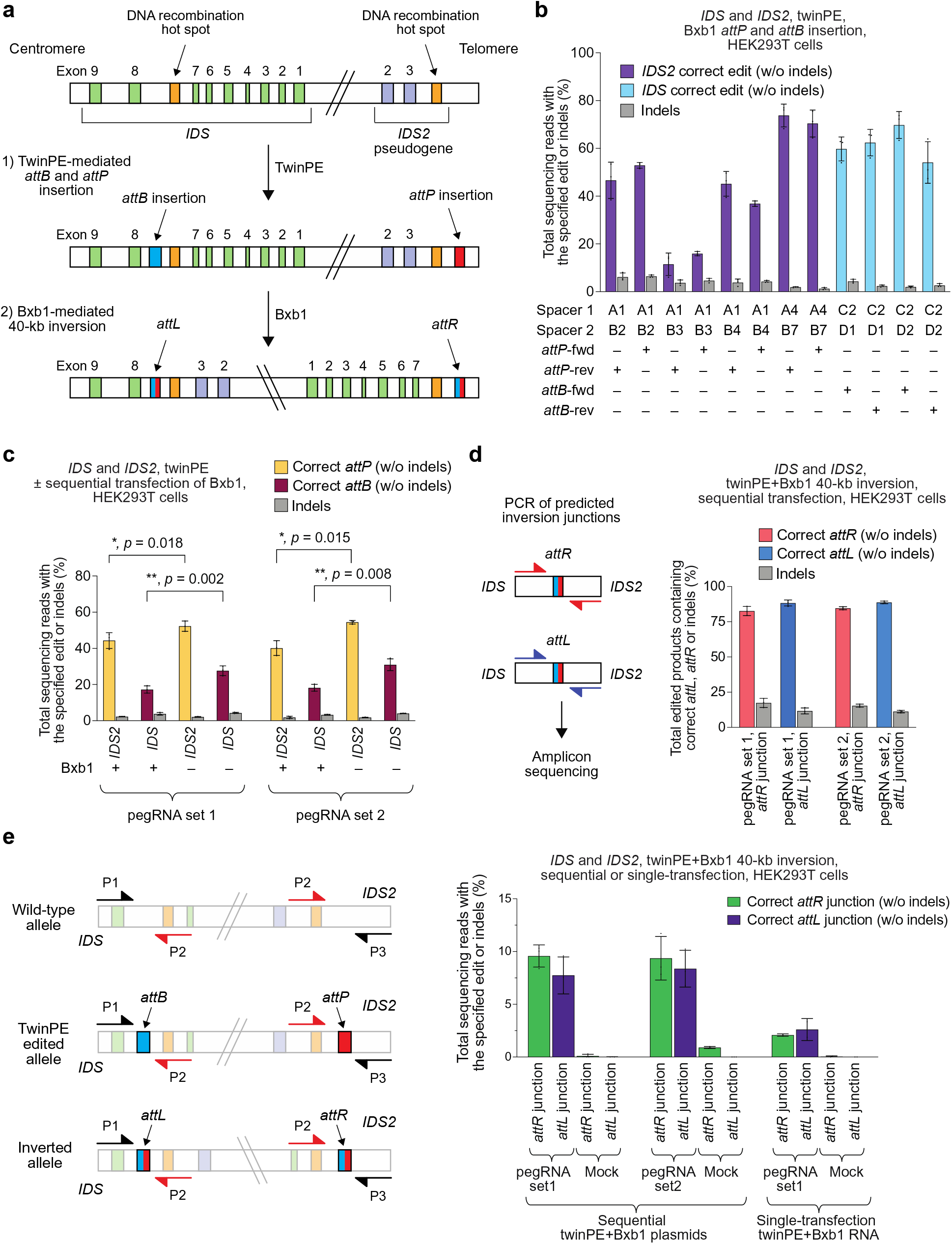
Site-specific large genomic sequence inversion with twinPE and Bxb1 recombinase in human cells. (**a**) Schematic diagram of DNA recombination hot spots in *IDS* and *IDS2* that lead to pathogenic 39-kb inversions, and the combined twinPE-Bxb1 strategy for installing or correcting the *IDS* inversion. (**b**) Screen of pegRNA pairs at *IDS* and *IDS2* for insertion of *attP* or *attB* recombination sites. (**c**) DNA sequencing analysis of the *IDS* and *IDS2* loci after twinPE-mediated insertion of *attP* or *attB* sequences, with or without subsequent transfection with Bxb1 recombinase. P-values were derived from a Student’s two-tailed *t*-test. (**d**) 40,167-bp *IDS* inversion product purities at the anticipated inversion junctions after twinPE-mediated attachment site installation and sequential transfection with Bxb1 recombinase. (**e**) Analysis of inversion efficiency by amplicon sequencing at *IDS* and *IDS2* loci after sequential transfection or single-step transfection of twinPE editing components and Bxb1 recombinase. Values and error bars reflect the mean and s.d. of three independent biological replicates for (b, c and d) and two or three independent biological replicates for (e).

Next, we explored multiplexing twinPE-mediated insertion of both *attB* and *attP* attachment sites with Bxb1 recombinase-mediated inversion of the 40-kb sequence in *IDS* and *IDS2*. Initially, we explored sequential DNA transfections with twinPE components followed by Bxb1 recombinase. A first set of pegRNAs (set 1) was tested that installs a reverse-oriented *attP* sequence in intron 3 of *IDS2* and a forward-oriented *attB* sequence in intron 7 of *IDS*. In addition, a second set of pegRNAs (set 2) was used to install a forward-oriented *attP* sequence in *IDS2* and a reverse-oriented *attB* sequence in *IDS*. We observed 52% or 55% *attP* sequence insertion and 28% or 31% *attB* sequence insertion for multiplexed editing with pegRNA set 1 and set 2, respectively (Fig. 4c). When edited cells were subsequently transfected with Bxb1 recombinase, we observed a significantly decreased percentage of amplified alleles containing *attP* and *attB* sequence compared to mock transfection controls (p-value < 0.05), suggesting that some edited alleles were used as substrates for Bxb1-mediated recombination (Fig. 4c). Amplification of the anticipated inversion junctions followed by amplicon sequencing revealed the presence of both *attL* and *attR* Bxb1 recombination products with product purity at or above 80% at both junctions (Fig. 4d).

To carry out one-step twinPE and Bxb1-mediated inversion and circumvent unwanted recombination between pegRNA plasmid DNA, we nucleofected all-RNA components comprising PE2 mRNA, synthetic pegRNAs from set 1, and Bxb1 mRNA. Using amplicon sequencing, we captured the expected inverted allele junctions containing *attR* and *attL* sequences (Extended Data Fig. 9). To quantify inversion efficiency, we designed a reverse primer that can bind to an identical sequence in both the non-inverted and inverted alleles and therefore amplify both edits using the same primer pair (Extended Data Fig. 9). We observed 7.7–9.6% and 2.1–2.6% inversion efficiency for sequential and one-step methods, respectively (Fig. 4e). Collectively, these data suggest that combining twinPE with site-specific serine recombinases can be used to correct a common 39-kb inversion found in Hunter syndrome alleles and may eventually serve as a therapeutic strategy for correcting other large or complex pathogenic gene variants.

## Discussion

We developed a twin prime editing approach that can be used to replace, insert, or delete DNA sequences at targeted locations in the genomes of human cells. First, we showed that twinPE can efficiently replace endogenous genomic DNA sequences with exogenous sequences containing Bxb1 *attB* or *attP* recombination sites with observed editing efficiencies of >80% at *HEK3* and >40% in four different genomic loci in HeLa, U2OS, and K562 cells (Extended Data Fig. 10). An active RT is required to achieve these edits, as Cas9 nickase (H840A) and Cas9 nickase fused to an inactivated RT (K103L/R110S) both failed to generate the desired edits at all sites tested, in contrast to efficient editing with twinPE (Extended Data Fig. 10c). We further used twinPE to recode portions of exon sequences within *PAH*, which in principle could be used to correct multiple pathogenic variants associated with different phenylketonuria patient cohorts using a single prime editor and pegRNA pair. Moreover, since twinPE does not require pegRNAs with RT templates that possess homology to the target site, a feature that is typically required for standard prime editing, and because each RT template does not need to possess the entire replacement sequence, twinPE pegRNAs can encode larger insertions for a given RT template length, including the insertion of a 108-bp sequence with 16% efficiency using twinPE, representing a 20-fold improvement over PE3.

We also established precise and flexible twinPE deletion strategies, which make use of the programmability of 3’ flap sequences that can be used to fully specify deletion junctions. Previously, this was a limitation of nuclease-based deletions strategies, which are largely limited to deletions between the cut sites and therefore nuclease accessibility. Using twinPE, we tested three deletion strategies, including the previously reported PrimeDel strategy^38^. TwinPE achieved precise deletions of up to 780-nt in *DMD* sequences including the deletion of exon 51 with up to 28% efficiency and 4% total indels. Although Cas9 nuclease-mediated deletion at this site can achieve 50% deletion efficiency, Cas9 nuclease generates 45% indels in the process. TwinPE thus may be especially useful for applications in which uncontrolled indel byproducts are undesired.

Although efficient targeting of both DNA protospacers is likely necessary for achieving efficient twinPE, we observed only weak correlations between *in silico* predictions of Cas9 spacer activity and observed twinPE editing efficiencies (Supplementary Note 3). This poor correlation could arise from other determinants of twinPE efficiency beyond the protospacer, including RT template and PBS choice. While PBS optimization was important for high editing efficiency at many sites, we did not observe an optimal PBS melting temperature across the pegRNA designs in this study. A correlation between prime editing efficiency and the distance between pegRNA-induced nicks was observed, which may suggest an optimal spacing of 50 to 100 bp, although many exceptions exist (Supplementary Note 3). Finally, the use of epegRNAs instead of canonical pegRNAs improved twinPE editing efficiency in many cases (Fig. 2c, 2e, and Extended Data Fig. 9c). Therefore, empirical testing of epegRNA pairs will continue to be important to optimize the efficiency of twinPE.

When combined with site-specific serine recombinase enzymes, twinPE can also support the integration of large DNA cargo, or the inversion of DNA sequence. To enable targeted integration of gene-sized DNA plasmids, we combined twinPE and Bxb1 recombinase in a single transfection and successfully inserted a 5.6-kb DNA donor plasmid into the genome of human cells at safe harbor loci *AAVS1, CCR5*, and *ALB* with up to 6.8%, 6.1%, and 2.6% efficiency, respectively. Notably, Bxb1-mediated knock-in of an *attP* donor into a HEK293T clonal cell line bearing homozygous *attB* integration peaked at 17% in our hands, consistent with observations from others^56^. Given that twinPE-mediated insertions of *attB* and *attP* sequences already can exceed 80%, improving the activity of Bxb1 integrase is likely the best opportunity to enhance overall donor knock-in efficiencies. As Bxb1 has been used to insert up to 27 kB in mammalian cells^54^, even larger insertions may be possible. Notably, the orthogonality of dinucleotide core sequences may enable gene-sized insertions to be multiplexed to allow simultaneous targeting of two loci with two different donors.

The methods developed here should be applicable for genome engineering and could be applied in the future for the targeted integration of therapeutic genes. TwinPE and Bxb1-mediated recombination offer advantages over other approaches. The serine integrase phiC31^65^ and fusions of zinc fingers or dCas9 to the catalytic domain of Gin recombinase^35, 66, 67^ have been used to integrate or excise DNA at endogenous pseudo-sites in the human genome, but the extensive sequence preferences inherent to these recombinases greatly limits the number of targetable genomic loci to a minute fraction of genomic loci. The programmable integration of attachment sites by twinPE overcomes many of these challenges by enabling insertion of cognate recombinase recognition sequences at any PE-targetable locus. By comparison to nuclease-based methods, the twinPE and recombinase approach avoids the generation of DSBs, which typically lead to uncontrolled mixtures of by products^19^, and can also lead to chromosomal rearrangements^23^, chromothripsis^24^, large deletions^20–22^, and p53 activation^25–27^. Integration orientation using twinPE and Bxb1 recombinase is strictly controlled by the directionality of the *attB* and *attP* sequences^62^, in contrast to uncontrolled integration orientation using homology-independent repair^14^. Methods that leverage HDR also enable control of sequence orientation, and can achieve efficiencies on the order of 5-10% without drug selection or suppression of NHEJ^68–70^, but HDR is less efficient than NHEJ in most cell types and typically requires DSBs^16, 17^. Methods have also been developed for making targeted gene-sized insertions through paired nicking of the genome and a donor cassette^71, 72^. These approaches do not require double strand breaks, but remain reliant on HDR and supportive cell types^72^.

We used twinPE multiplex insertion of *attP* and *attB* with Bxb1 to correct a large sequence inversion at *IDS* and *IDS2* associated with Hunter syndrome, resulting in precise inversion of 40 kB in *IDS* with 9.6% editing efficiency via sequential transfection and 2.6% editing efficiency via one-step RNA nucleofection (Fig. 4e). Although zinc finger nuclease, TALEN, and Cas9 nucleases-based approaches have achieved targeted DNA sequence inversions previously^73–75^, DSBs-induced repair pathways also generate undesired products and can lead to *de-novo* structural variants, including deletion of the targeted DNA sequence. TwinPE and Bxb1 circumvented the issues of nuclease-induced DSBs and enable comparable (19% by TwinPE+Bxb1 vs. 23% by Cas9^75^ for a 1.2-kb inversion) or higher inversion efficiencies (9.6% by TwinPE+Bxb1 for 40-kb vs. 1.4% by TALEN^74^ for a 140-kb inversion) through a precise series of reactions that circumvent the uncontrolled nature of DSB repair.

TwinPE thus expands the capabilities of prime editing to include targeted deletion, replacement, integration, or inversion of larger DNA sequences at specified locations in the genome. By combining twinPE and Bxb1, we achieved gene-size DNA integration and inversion without double-strand breaks, which may provide a strategy to potentially treat genetic diseases arising from loss-of-function or complex structural mutations. Except for spacer sequences and the 8-13 base pairs needed to prime reverse transcription, pegRNAs and donor plasmids in our method do not require homology to the genome. Additional research is needed to fully understand the cellular determinants of twinPE outcomes. Performing large cellular screens to unveil DNA repair components that suppress or enhance twinPE editing efficiency could illuminate mechanistic details that further improve this approach. Future studies on orchestrating twinPE and Bxb1 activity through engineering either or both components could augment the scope and efficiency of twinPE and recombinase editing in mammalian cells.

## Supporting information

Complete SI

## Acknowledgements

We thank Erik Sontheimer’s group for sharing Huh7 cells and Liu Lab members for helpful discussions. This work was supported by the Merkin Institute of Transformative Technologies in Healthcare, US NIH grants U01 AI142756, RM1 HG009490, and R35 GM118062, and the HHMI. A.V.A. acknowledges a Jane Coffin Childs postdoctoral fellowship through the HHMI.

## Author contributions

A.V.A., X.D.G., C.J.P., A.T.N., and J.M.L. designed experiments. A.V.A., X.D.G., C.J.P., A.T.N., L.W.K., A.R., and J.A.M.M. performed experiments and analyzed data. A.V.A., X.D.G., C.J.P. and D.R.L. wrote the manuscript. D.R.L. supervised the research.

## Methods

### General methods

DNA amplification was conducted by PCR using Phusion U Green Multiplex PCR Master Mix (ThermoFisher Scientific) or Q5 Hot Start High-Fidelity 2x Master Mix (New England BioLabs) unless otherwise noted. DNA oligonucleotides were obtained from Integrated DNA Technologies. Plasmids expressing sgRNAs were constructed by ligation of annealed oligonucleotides into *Bsm*BI-digested acceptor vector as previously described^12^.Plasmids expressing pegRNAs were constructed by Gibson assembly or Golden Gate assembly as previously described^12^. Sequences of sgRNA and pegRNA constructs used in this work are listed in Supplementary Table 1. All vectors for mammalian cell experiments were purified using Plasmid Plus Midiprep kits (Qiagen), PureYield plasmid miniprep kits (Promega), or QIAprep Spin Miniprep kits. Synthetic pegRNAs were ordered from IDT without HPLC purification.

### General mammalian cell culture conditions

HEK293T (ATCC CRL-3216), U2OS (ATTC HTB-96), K562 (CCL-243), and HeLa (CCL-2) cells were purchased from ATCC and cultured and passaged in Dulbecco’s Modified Eagle’s Medium (DMEM) plus GlutaMAX (ThermoFisher Scientific), McCoy’s 5A Medium (Gibco), RPMI Medium 1640 plus GlutaMAX (Gibco), or Eagle’s Minimal Essential Medium (EMEM, ATCC), respectively, each supplemented with 10% (v/v) fetal bovine serum (Gibco, qualified) and 1x Penicillin+Streptomycin (Corning). All cell types were incubated, maintained, and cultured at 37 °C with 5% CO_2_. Cell lines were authenticated by their respective suppliers and tested negative for mycoplasma.

### HEK293T, HeLa, and Huh7 tissue culture transfection protocol and genomic DNA preparation

HEK293T cells were seeded on 48-well poly-D-lysine coated plates (Corning). 16-24 h post-seeding, cells were transfected at approximately 60% confluency with 1 µL of Lipofectamine 2000 (Thermo Fisher Scientific) according to the manufacturer’s protocols and either: 750 ng of PE2 plasmid DNA, 125 ng of pegRNA 1 plasmid DNA, and 125 ng of pegRNA 2 plasmid DNA (for twinPE transfections); 750 ng of PE2 plasmid DNA, 250 ng of pegRNA plasmid DNA, and 83 ng of sgRNA plasmid DNA (for PE3 transfections); or, 750 ng of Cas9 plasmid DNA and 125 ng of sgRNA 1 plasmid DNA, and 125 ng of sgRNA 2 plasmid DNA (for paired Cas9 nuclease transfections). Unless otherwise stated, cells were cultured 3 days following transfection, after which the media was removed, the cells were washed with 1x PBS solution (Thermo Fisher Scientific), and genomic DNA was extracted by the addition of 150 µL of freshly prepared lysis buffer (10 mM Tris-HCl, pH 7.5; 0.05% SDS; 25 µg/mL Proteinase K (ThermoFisher Scientific)) directly into each well of the tissue culture plate. The genomic DNA mixture was incubated at 37 °C for 1-2 h, followed by an 80 °C enzyme inactivation step for 30 min. Primers used for mammalian cell genomic DNA amplification are listed in Supplementary Table 2. For HeLa cell transfections, cells were grown and seeded in 96-well plates (Falcon Catalog #353075). 16-24 h post-seeding, cells were transfected 0.75 µL of TransIT-HeLaMONSTER transfection reagent (Mirus) using 190 ng of PE2-P2A-Blast plasmid DNA, 31.5 ng of pegRNA 1 plasmid DNA, and 31.5 ng of pegRNA 2 plasmid DNA (for twinPE transfections). 24 h after transfection, cells were treated with blasticidin to a final concentration of 10 µg/mL. Genomic DNA isolation was performed as described above for HEK293T cells using 50 µL of lysis buffer. Huh7 cells were seeded at 150,000 cells per well in poly-D-lysine coated 24-well plates (Corning). 16-24 h post-seeding, cells were transfected with 2 µL of Lipofectamine 2000 (Thermo Fisher Scientific) according to the manufacturer’s protocols and up to 800 ng of plasmid DNA (same ratios as in HEK293T transections described above, scaled proportionally). Genomic DNA isolation was performed as described above for HEK293T transfections.

### High-throughput DNA sequencing of genomic DNA samples

Genomic sites of interest were amplified from genomic DNA samples and sequenced on an Illumina MiSeq as previously described with the following modifications^10^. Briefly, amplification primers containing Illumina forward and reverse adapters (Supplementary Table 2) were used for a first round of PCR (PCR 1) amplifying the genomic region of interest. 25-µL PCR 1 reactions were performed with 0.5 µM of each forward and reverse primer, 1 µL of genomic DNA extract and 12.5 µL of Phusion U Green Multiplex PCR Master Mix. PCR reactions were carried out as follows: 98 °C for 2 min, then 30 cycles of [98 °C for 10 s, 61 °C for 20 s, and 72 °C for 30 s], followed by a final 72 °C extension for 2 min. Unique Illumina barcoding primer pairs were added to each sample in a secondary PCR reaction (PCR 2). Specifically, 25 µL of a given PCR 2 reaction contained 0.5 µM of each unique forward and reverse Illumina barcoding primer pair, 1 µL of unpurified PCR 1 reaction mixture, and 12.5 µL of Phusion U Green Multiplex PCR 2x Master Mix. The barcoding PCR 2 reactions were carried out as follows: 98 °C for 2 min, then 12 cycles of [98 °C for 10 s, 61 °C for 20 s, and 72 °C for 30 s], followed by a final 72 °C extension for 2 min. PCR products were evaluated analytically by electrophoresis in a 1.5% agarose gel. PCR 2 products (pooled by common amplicons) were purified by electrophoresis with a 1.5% agarose gel using a QIAquick Gel Extraction Kit (Qiagen), eluting with 40 µL of water. DNA concentration was measured by fluorometric quantification (Qubit, ThermoFisher Scientific) or qPCR (KAPA Library Quantification Kit-Illumina, KAPA Biosystems) and sequenced on an Illumina MiSeq instrument according to the manufacturer’s protocols.

Sequencing reads were demultiplexed using MiSeq Reporter (Illumina). Alignment of amplicon sequences to a reference sequence was performed using CRISPResso2^76^. For all prime editing yield quantification, prime editing efficiency was calculated as: % of [# of reads with the desired edit that do not contain indels] ÷ [# of total reads]. For quantification of editing, CRISPResso2 was run in HDR mode using the desired allele as the expected allele (e flag), and with “discard_indel_reads” on. Any sequence containing an indels with respect to the allele to which it aligns was counted separately and did not contribute to the correctly edited allele percentage. The percent editing was quantified as the number of non-discarded reads aligning to the anticipated edited allele (not containing indels) divided by the total number of sequencing reads (which includes those aligned to reference with or without indels and those aligned to the desired edit with or without indels). Indels were quantified as the total number of discarded reads (from either the original or edited allele alignments) divided by the total number of sequencing reads. Editing yield was then calculated as: [# of non-discarded HDR aligned reads] ÷ [total reads]. Indel yields were calculated as: [# of indel-containing discarded reads] ÷ [total reads].

Unique molecular identifiers (UMIs) were applied to quantify the deletion efficiency and assess PCR bias (see Supplementary Note 1 for discussion) in a three-step PCR protocol. Briefly, in the first step of linear amplification, 1 µL of genomic DNA extract was linearly amplified by 0.1 µM of only the forward primer containing a 15-nt or 16-nt UMI with Phusion U Green Multiplex PCR Master Mix in a 25-µL reaction (10 cycles of 98 °C for 1 min, 61 °C for 25 s, and 72 °C for 1 min). The PCR products were then purified by 1.6X AmPure beads (Beckman Coulter) and eluted in 20 µL of QIAGEN elution buffer. In the second step, 1 or 2 µL of purified linearly amplified PCR products were then amplified for 30 cycles with 0.5 µM of each forward and reverse primer with Phusion U Green Multiplex PCR mix in a 25-µL reaction as described above. In this case, the forward primer anneals to the P5 Illumina adaptor sequence located at the 5’ of the UMI primer and upstream of the UMI sequence. For the *DMD* locus library preparation, the PCR products were purified by 1X AmPure beads and eluted in 25 µL of elution buffer. In the third step, the purified PCR products (1 µL) were amplified for 12 cycles as described above for adding unique Illumina barcodes and adaptors. To assess large deletions at the *DMD* locus, the top band (unedited large amplicon) and bottom band (edited amplicons with deletions) were excised separately from a 1.5% agarose gel and loaded on two separate MiSeq runs to avoid biased clustering of amplicons. For library preparation at other loci, 1 µL of PCR product was used directly for the barcoding PCR step without 1X AmPure beads purification. For UMI-based PCR bias assessment described in Supplementary Note 1, the editing efficiency was calculated from the libraries prepared following the UMI protocols analyzed with UMI-deduplication or without UMI-deduplication.

Raw sequencing reads were UMI deduplicated using AmpUMI^77^. For paired-end reads, SeqKit^78^ was used to concatenate (merge without overlap) R1s with the reverse complement of R2s. The concatenated R1+R2s were UMI deduplicated using the UMI at the 5’ end of R1. UMI-deduplicated R1s or concatenated R1+R2s were analyzed using CRISPResso2. For analyzing concatenated R1+R2s, an appropriate concatenated reference amplicon sequence was provided to minimize sequencing alignment artifacts due to the concatenation.

### Nucleofection of U2OS and K562

Nucleofection was performed in all experiments that used K562 and U2OS cells. 200,000 cells were used per nucleofection. Counted cells were pelleted and washed with PBS, then resuspended in nucleofection solution following the recommendation of Lonza SE Cell Line 4D-Nucleofector Kit. After nucleofection of the cells, the cells were allowed to incubate in the cuvette at room temperature for 10 minutes. After this time, the contents of the cuvette were transferred to a 48 well plate containing pre-warmed (37 °C) media. Genomic DNA was extracted and prepared for Illumina MiSeq preparation as described above.

### Single-step twinPE and Bxb1-mediated DNA donor knock-in and inversions

For single-step knock-in, HEK293T cells were transfected with 500 ng of PE2 plasmid DNA, 50 ng of each pegRNA plasmid DNA (two in total), 200 ng of codon-optimized Bxb1 plasmid DNA, and 200 ng of DNA donor using Lipofectamine 2000 as described above. For multiplex knock-in, HEK293T cells were transfected with 200 ng of PE2 plasmid, 50 ng of each pegRNA plasmid DNA (four in total), 300 ng of Bxb1 plasmid DNA, and 150 ng of each DNA donor (two in total). Donor plasmid sequences can be found in Supplementary Sequences 1. The codon-optimized Bxb1 gene was purchased from Genscript and can be found in Supplementary Sequence 2. It was assembled into a pCMV expression vector using standard molecular cloning techniques.

For inversion experiments, sequential plasmid transfection and single-step mRNA nucleofection were performed. In the sequential plasmid transfection experiment, HEK293T cells were transfected with 750 ng of PE2 plasmid DNA, 62.5 ng of each pegRNA plasmid DNA (four in total) using Lipofectamine 2000 as described above. After three days, cells were detached and plated in 24-well plates, then serially passaged for about seven days. 48-well plates were seeded with 20,000 cells per well and transfected 16-24 h later with Lipofectamine 2000 and 500 ng of BxB1 plasmid DNA. Genomic DNA was extracted and prepared for Illumina MiSeq as described above. For single-step mRNA nucleofection, 200,000 HEK293T cells were nucleofected with 1,000 ng of PE2 mRNA, 30 pmol each of synthetic pegRNAs (4 pegRNAs in total), and 750 ng of BxB1 mRNA using 20 µL of Lonza buffer and cells with program CM-130. Cells were recovered with 80 µL of pre-warmed (37 °C) media for five minutes. 25-µL samples from the nucleofection cuvette were then added to each well of the 48-well plate and incubated at 37 °C for 72 hours prior to isolation of genomic DNA as described above.

### Droplet digital PCR analysis of knock-in efficiency

Genomic DNA from crude cell lysates was column purified (Zymo) and DNA concentrations were determined by Nanodrop (Thermo Scientific). Droplet digital PCR was used to determine the abundance of an amplicon containing the genome-donor junction in comparison to a reference gene (*ACTB*). 100-200 ng of DNA was added to a reaction mixture containing ddPCR Supermix for Probes (Bio-Rad, 1863026), HindIII-HF (0.25 units/μL, New England BioLabs, R3104L), *ACTB* primers and probes (Supplementary Table 3; 900 nM each primer, 250 nM probe) and genome-donor junction primers and probes (Supplementary Table 3; 900 nM each primer, 250 nM probe) according to the manufacturer’s protocol. Droplets were generated using a QX200 Manual Droplet Generator (Bio-Rad, 186-4002). Digital droplet PCR was performed as follows: 95 °C for 10 min, then 50 cycles of 94 °C for 30 seconds, 58 °C for 2 min. Following PCR cycles, a final incubation was conducted at 98 °C for 10 min. Droplets were read by a QX200 Droplet Reader (Bio-Rad, 1864001) and data were analyzed using QuantaSoft (Bio-Rad).

### FIX expression from ALB locus in Huh7 cells

Huh7 cells were transfected with 160 ng of Bxb1 plasmid DNA, 160 ng of donor plasmid DNA (*attP*-splice acceptor-cDNA of *F9* exons 2-8), 400 ng of prime editor plasmid DNA, and pegRNAs plasmid DNA for installation of *attB* in the first intron of *ALB* or at *CCR5* (40 ng each). Three days post-transfection, cells were passaged and allowed to grow to confluence. Their media was changed, and they were left to condition the fresh media, with aliquots taken at days 4, 7, and 10. Factor IX concentration in conditioned media was measured by ELISA (Innovative Research Human Total Factor IX ELISA Kit, IHUFIXKTT).

### Lentivirus production, GFP reporter cell line construction, and flow cytometry analysis

The lentiviral transfer vector described in Extended Data Figure 9 and Supplementary Sequence 3 was assembled using standard molecular cloning techniques. The lentivirus was produced as previously described^79, 80^. Briefly, a T-75 flask of rapidly dividing HEK293T cells (ATCC; Manassas, VA, USA) were transfected at 80-90% confluence with lentivirus helper plasmids pVSV-G and psPAX2 in along with the constructed lentiviral transfer vector using FuGENE HD (Promega) according to the manufacturer’s protocol. After 48 hours, supernatant was collected, centrifuged at 3,000 x *g* for 15 minutes to remove cellular debris, and filtered using a 0.45-µm PVDF filter. Filtered supernatant was concentrated using the PEG-it Virus Precipitation Solution (System Biosciences) according to the manufacturer’s instructions. The resulting pallet was resuspended in Opti-MEM (Thermo Fisher Scientific) using 1% of the original medium volume. Resuspended pellet was used to transduce the HEK293T cells with multiplicity of infection. Successfully transduced cells that had stable GFP reporter integration were selected with 2.5 µg/mL puromycin for more than 3 passages and then seeded into 48-well plates for transfection with PE2 plasmid DNA (750 ng), four *AAVS1*-targeting pegRNAs (62.5 ng each) for multiplexed *attP* and *attB* insertion, and Bxb1 plasmid DNA (100 ng, 200 ng, 500 ng, or 1000 ng) in a single transfection step. Cells were then collected after three days and analyzed by CytoFLEX S Flow Cytometer (Beckman Coulter) and FlowJo v10.

### Production of PE2 and Bxb1 mRNA

The mRNA was produced using the previously reported protocol^41, 81^. Briefly, the PE2 editor and codon optimized Bxb1 coding sequences were cloned into a plasmid encoding a mutated T7 promoter that cannot initiate with mononucleotides. This plasmid was used as template in a PCR reaction (PhusionU 2X Master Mix) using PAGE-purified primers (IDT) Forward: 5’-TCGAGCTCGGTACCTAATACGACTCACTATAAGG-3’ and Reverse: 5’-TTTTTTTTTTTTTTTTTTTTTTTTTTTTTTTTTTTTTTTTTTTTTTTTTTTTTTTTTTTTT TTTTTTTTTTTTTTTTTTTTTTTTTTTTTTTTTTTTTTTTTTTTTTTTTTTTTTTTTTCTTCCTAC TCAGGCTTTATTCAAAGACCA-3’, and the resulting PCR product was purified and used as template for *in vitro* transcription. The HiScribe T7 High-Yield RNA Synthesis Kit (New England BioLabs) was used according to the company’s recommended instruction but with full substitution of N1-methyl-pseudouridine for uridine and co-transcriptional capping with CleanCap AG (TriLink Biotechnologies). RNA was precipitated with lithium chloride (2.5 M final concentration), the pellet was washed with ice cold 70% ethanol, and then resuspended in water.

### Data availability

High-throughput sequencing data have been deposited to the NCBI Sequence Read Archive database under accession PRJNA770428.

---

**Extended Data Figure 1.**
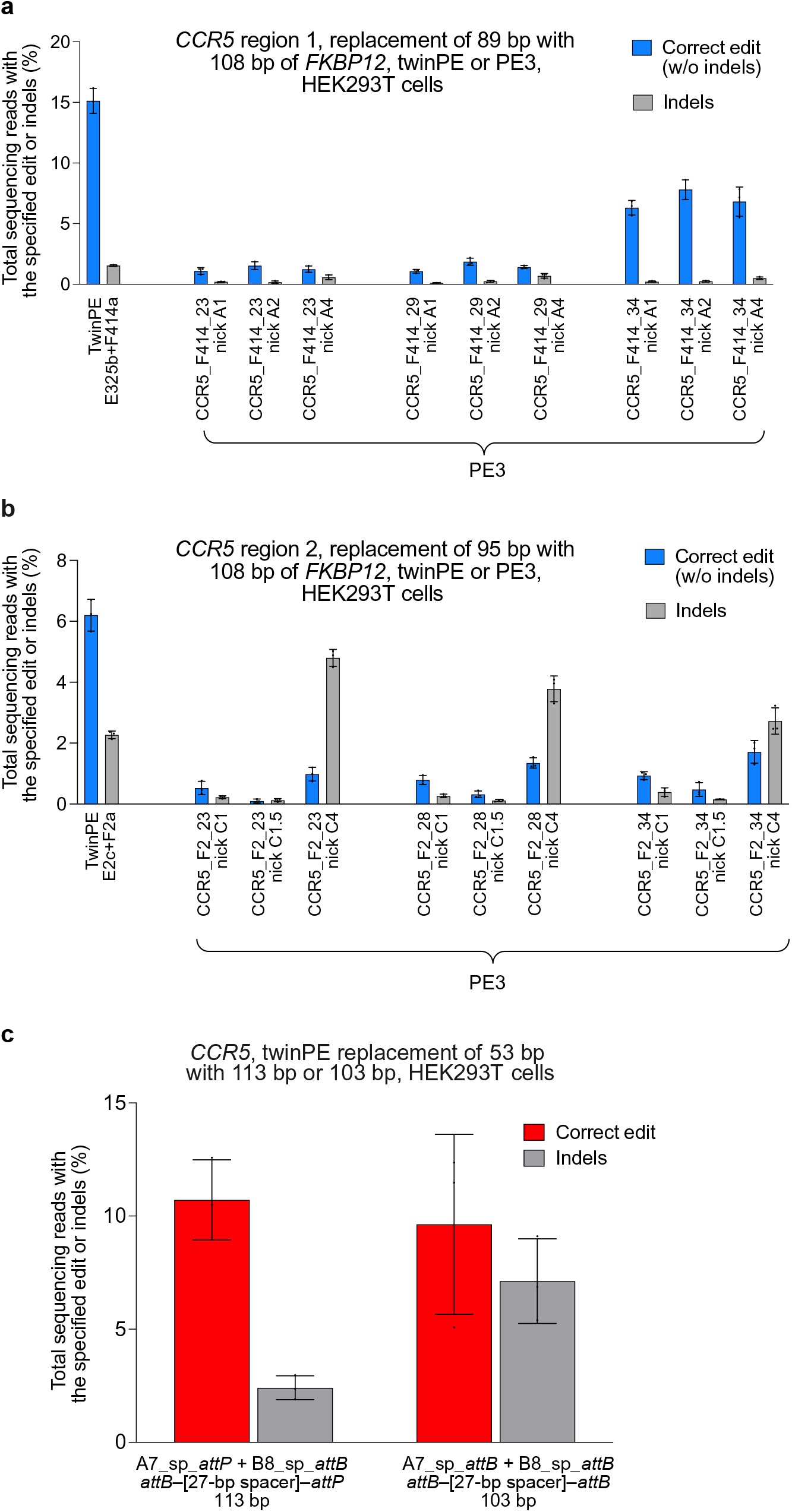
Twin prime editing mediates sequence replacements at *CCR5*. (**a**) Replacement of endogenous sequence within *CCR5* region 1 with a 108-bp fragment of *FKBP12* cDNA using twinPE (*FKBP12* sequence oriented in the forward direction,) or PE3 (*FKBP12* sequence oriented in the reverse direction). For PE3 editing, pegRNA RT templates were designed to encode 108 base pairs of *FKBP12* cDNA sequence and one of three different target-site homology sequence lengths. For PE3 edits, each pegRNA was tested with three nicking sgRNAs. (**b**) Replacement of endogenous sequence within *CCR5* region 2 with a 108-bp fragment of *FKBP12* cDNA sequence using twinPE (*FKBP12* sequence oriented in the forward direction) or PE3 (*FKBP12* sequence oriented in the reverse direction). As in (a), PE3 edits were tested with pegRNAs containing RT templates that were designed to encode 108 base pairs of *FKBP12* cDNA sequence and one of three different target-site homology sequence lengths. For PE3 edits, each pegRNA was tested with three nicking sgRNAs. Values and error bars reflect the mean and s.d. of three independent biological replicates. (**c**) Transfection of HEK293T cells with a pair of pegRNAs targeting *CCR5* leads to replacement of 53 base pairs of endogenous sequence with 113 base pairs (*attB*–[27-bp spacer]–*attP*) or 103 base pairs (*attB*–[27-bp spacer]–*attB*) of exogenous sequence. Values and error bars reflect the mean and s.d. of three independent biological replicates.

**Extended Data Figure 2.**
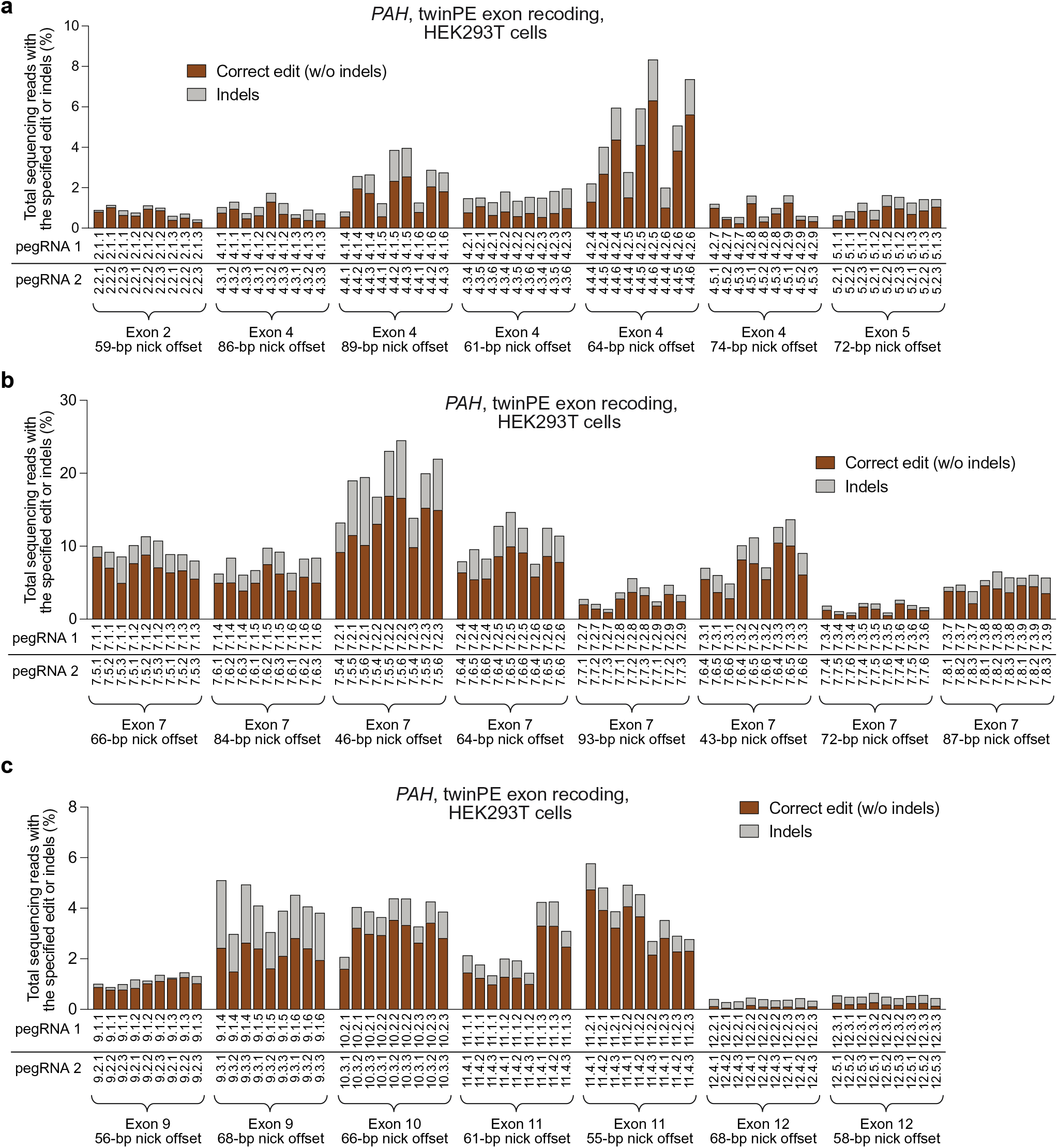
Recoding of *PAH* exon sequences in HEK293T cells via twinPE. Screen of pegRNA pairs targeting *PAH* for recoding of (**a**) exons 2, 4 and 5, (**b**) exon 7, and (**c**) exons 9, 10, 11, and 12. RT templates of pegRNAs encoded partially recoded exonic sequence to optimize orthogonality to the endogenous gene sequence. For each spacer pair, nine pegRNA combinations were tested using three PBS variants for each spacer in a three-by-three matrix, with RT templates encoding the recoded exonic sequence, which was held constant for given spacer pairs. Sequences of pegRNAs are listed in **Supplementary Table 1**. Sequences of recoded exonic sequences are listed in **Supplementary Table 4**. Values in (a), (b) and exon 9 in (c) reflect single biological replicates. Values for exons 10, 11 and 12 in (c) reflect the mean of three independent biological replicates.

**Extended Data Figure 3.**
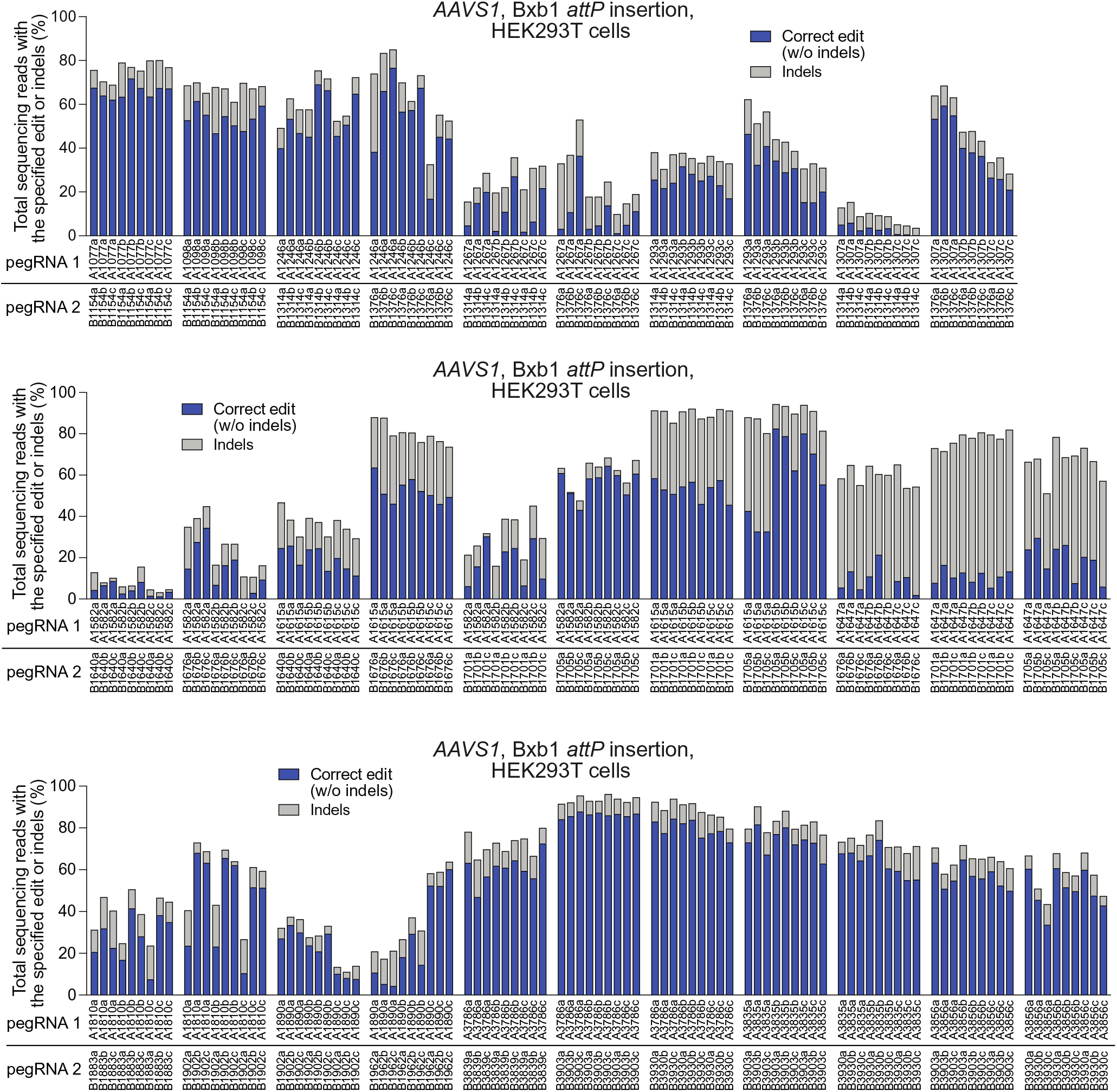
Installation of a 50-bp Bxb1*attP* site at *AAVS1* with twinPE. Spacer pairs targeting the *AAVS1* locus were designed for twinPE-mediated insertion of the Bxb1 *attP* attachment site. For each spacer, three pegRNAs were designed having three different PBS lengths and a fixed RT template that encodes a portion (43-44 bp) of the Bxb1 *attP* sequence. Sequences of pegRNAs are listed in **Supplementary Table 1**. For each spacer pair, a three-by-three matrix of pegRNA combinations was tested by plasmid DNA co-transfection with PE2 in HEK293T cells. Each pegRNA pair is specified below the x-axis. Values reflect single biological replicates.

**Extended Data Figure 4.**
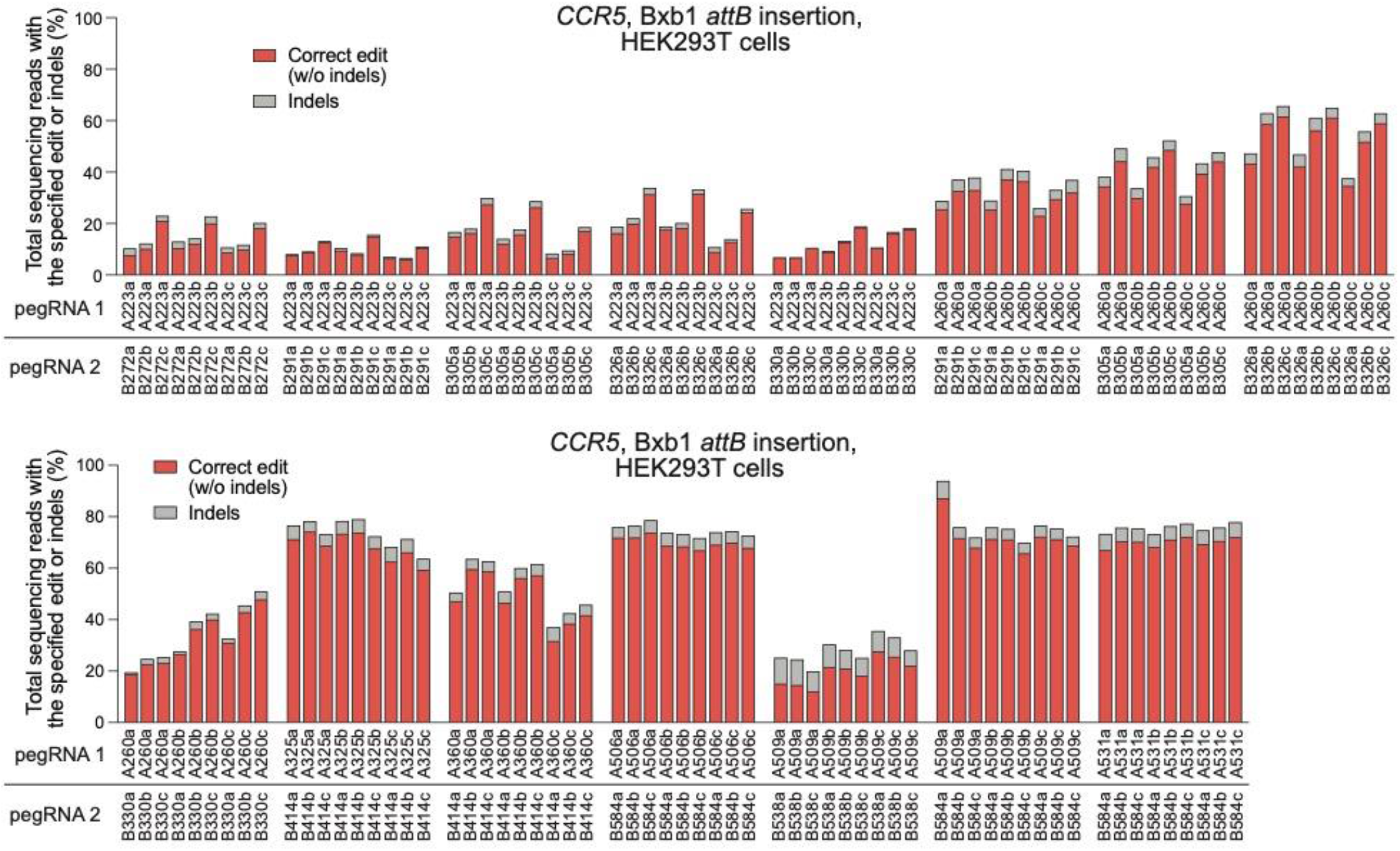
Installation of a 38-bp Bxb1 *attB* site at *CCR5* with twinPE. Spacer pairs targeting the *CCR5* locus were designed for twinPE-mediated insertion of the Bxb1 *attB* attachment site. For each spacer, three pegRNAs were designed having three different PBS lengths and a fixed RT template that encodes the full-length Bxb1 *attB* sequence (38 bp). Sequences of pegRNAs are listed in **Supplementary Table 1**. For each spacer pair, a three-by-three matrix of pegRNA combinations was tested by plasmid DNA co-transfection with PE2 in HEK293T cells. Each pegRNA pair is specified below the x-axis. Values reflect single biological replicates.

**Extended Data Figure 5.**
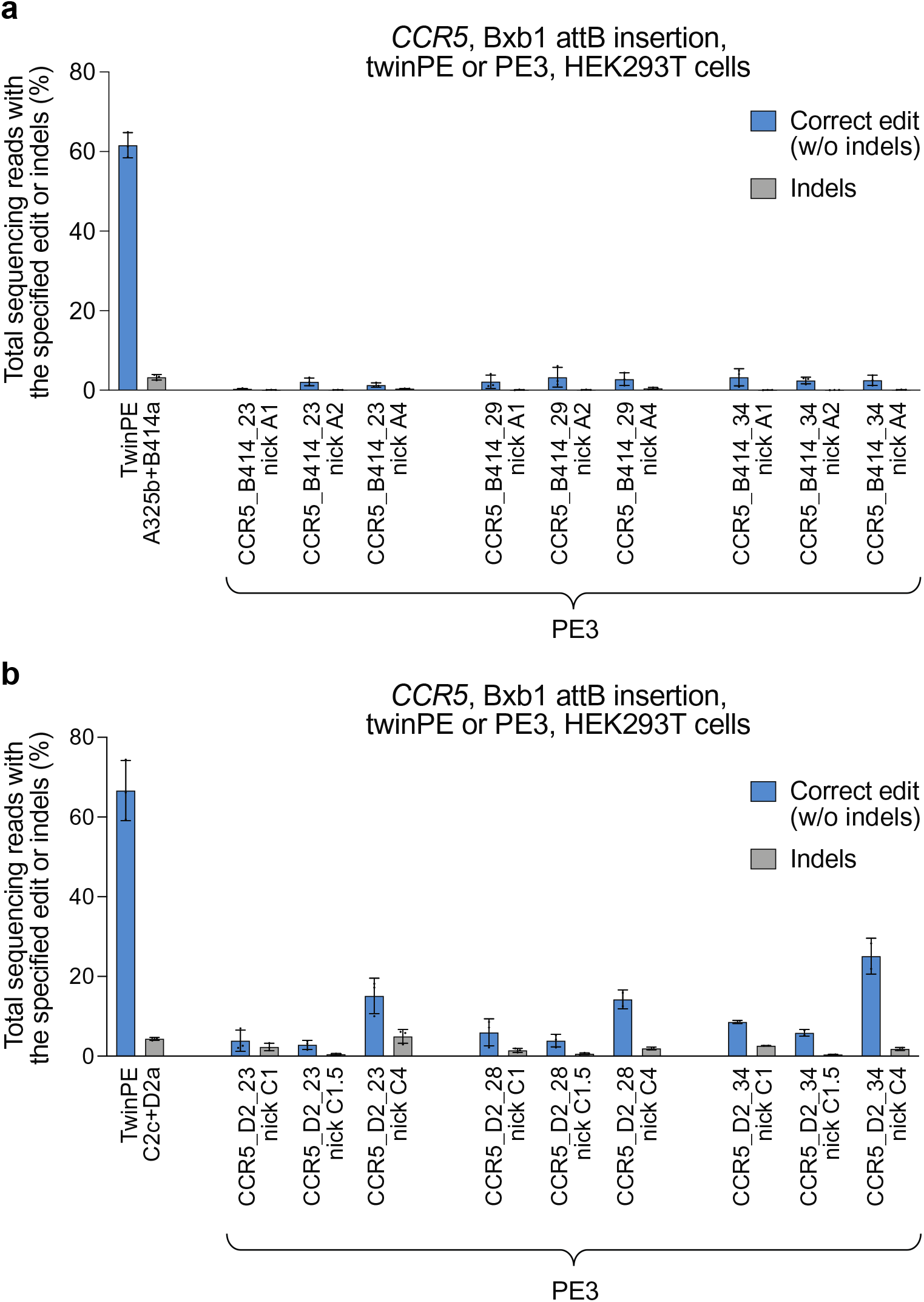
Comparison of twinPE and PE3 for Bxb1 *attB* insertion at *CCR5*. (**a**) Replacement of endogenous sequence within *CCR5* region 1 with the Bxb1 *attB* site using twinPE or PE3. For PE3 editing systems, pegRNA RT templates were designed to encode the Bxb1 *attB* sequence and one of three different target-site homology sequence lengths. For PE3 edits, each pegRNA was tested with three nicking sgRNAs. (**b**) Replacement of endogenous sequence within *CCR5* region 2 with the Bxb1 *attB* sequence using twinPE or PE3. As in (a), PE3 edits were tested with pegRNAs containing RT templates that were designed to encode the Bxb1 *attB* sequence and one of three different target-site homology sequence lengths and tested with three nicking sgRNAs. Values and error bars reflect the mean and s.d. of three independent biological replicates.

**Extended Data Figure 6.**
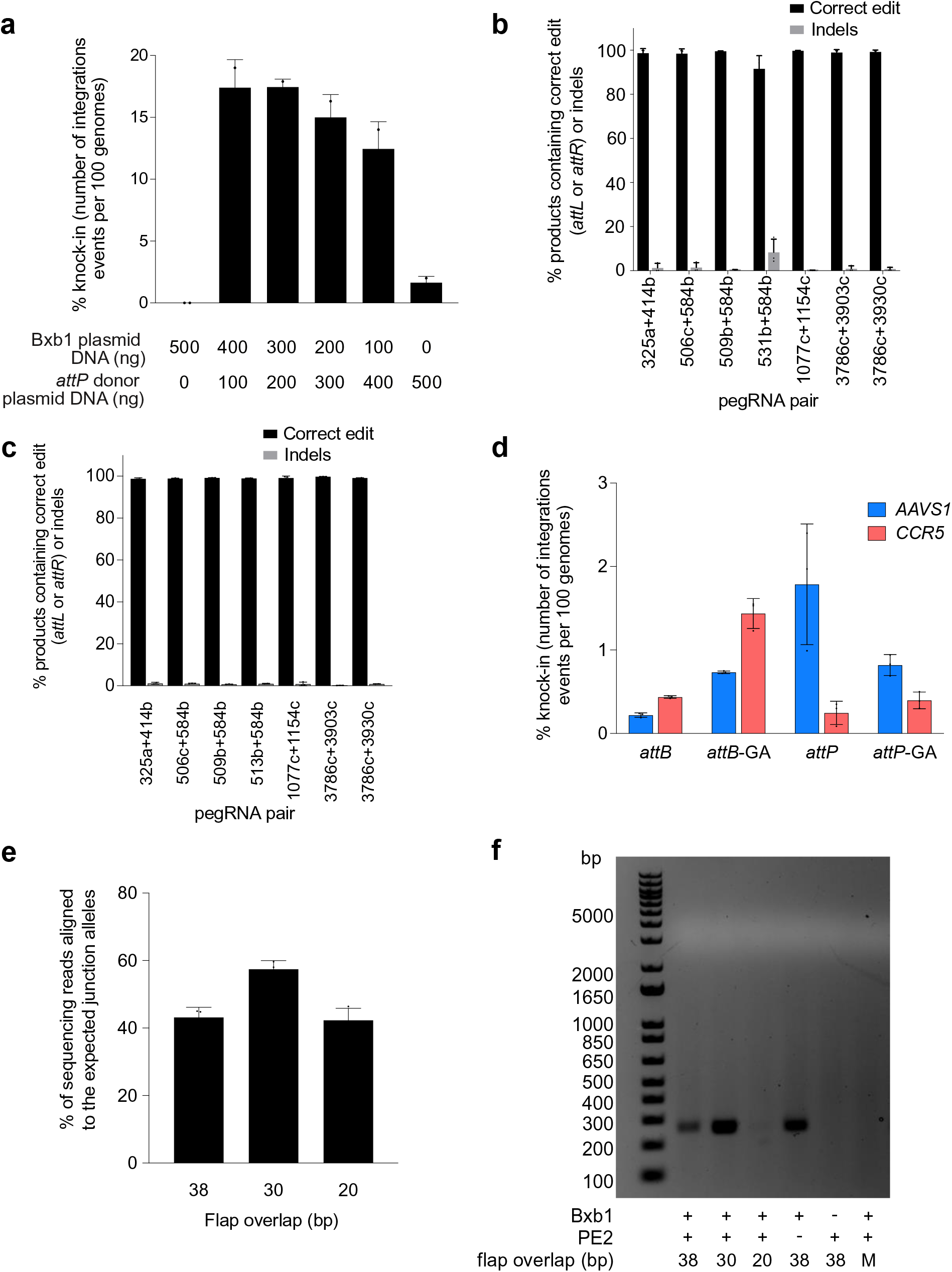
TwinPE combined with Bxb1 recombinase for targeted knock-in of donor DNA plasmids. (**a**) Bxb1-mediated DNA donor knock-in in clonal HEK293T cell lines. Transfection of a HEK293T clonal cell line containing homozygous *attB* site insertion at *CCR5* with varying amounts of Bxb1-expressing plasmid and *attP*-containing donor DNA plasmid. Knock-in efficiency was quantified by ddPCR. Values and error bars reflect the mean and s.d. of two independent biological replicates. (**b**) Assessment of genome-donor junction purity by high-throughput sequencing. Genomic DNA from single-transfection knock-in experiments was amplified with a forward primer that binds the genome and a reverse primer that binds within the donor plasmid (Supplementary Table 2). Values and error bars reflect the mean and s.d. of three independent biological replicates. (**c**) Assessment of genome-donor junction purity at the other junction by high-throughput sequencing as performed in (b). (**d**) Multiplexed single-transfection knock-in at *AAVS1* and *CCR5*. HEK293T cells were transfected with plasmids encoding PE2, Bxb1, a pair of pegRNAs for the insertion of *attP* at *AAVS1*, an *attB*-donor, a pegRNA pair for the insertion of one of four attachment sites (*attB*, *attB-GA*, *attP*, or *attP-GA*) at *CCR5*, and a corresponding donor. Knock-in was observed at both target loci under all four conditions. Insertion of *attP* at *AAVS1* and *attB* at *CCR5* gave the lowest knock-in efficiencies overall (0.2% at *AAVS1*, 0.4% at *CCR5*). Insertion of *attP* at both sites yielded the highest levels of knock-in at *AAVS1* (1.8%) but low levels (0.2%) at *CCR5*. When an orthogonal edit (*attB*-GA or *attP*-GA) was introduced at *CCR5*, *AAVS1* knock-in was 0.7-0.8%. Higher knock-in at *CCR5* was observed with *attB*-GA (1.4%) than with *attP*-GA (0.4%), consistent with our single locus knock-in results. Values and error bars reflect the mean and s.d. of three independent biological replicates. (**e**) and (**f**) Effects of reducing pegRNA overlap on twinPE efficiency and donor/pegRNA recombination. (**e**) The editing efficiencies of pairs of pegRNAs for insertion of Bxb1 *attB* at *CCR5* were measured by high-throughput sequencing. The pairs differed in the amount of overlap shared between their flaps, from 38 bp (full-length *attB* sequence) down to 20 bp. Editing efficiency of the pairs with shorter overlaps was comparable to the pair with full-length overlap. Values and error bars reflect the mean and s.d. of three independent biological replicates. (**f**) Assessment of recombination between *attB*-containing pegRNA plasmids and *attP*-containing donor plasmids. Following transfection of HEK293T cells with the indicated samples, isolated DNA was amplified with a forward primer that binds the pegRNA expression plasmid (TTGAAAAAGTGGCACCGAGT) and a reverse primer that binds the donor plasmid (CTCCCACTCATGATCTA). A positive 256-bp PCR band confirms recombination between the two plasmids. When the pegRNA encodes full-length *attB* (38-bp) or a truncated version of *attB* with 30-bp of overlap between flaps, a band is observed; however, recombination is not observed when the pegRNAs encode a truncated *attB* with only 20-bp of flap overlap. The “No PE2” control uses the 38-bp overlap pegRNA pair. No recombination is observed in the absence of Bxb1 or if the donor and pegRNA plasmids both bear *attB* (Mismatch, “M”).

**Extended Data Figure 7.**
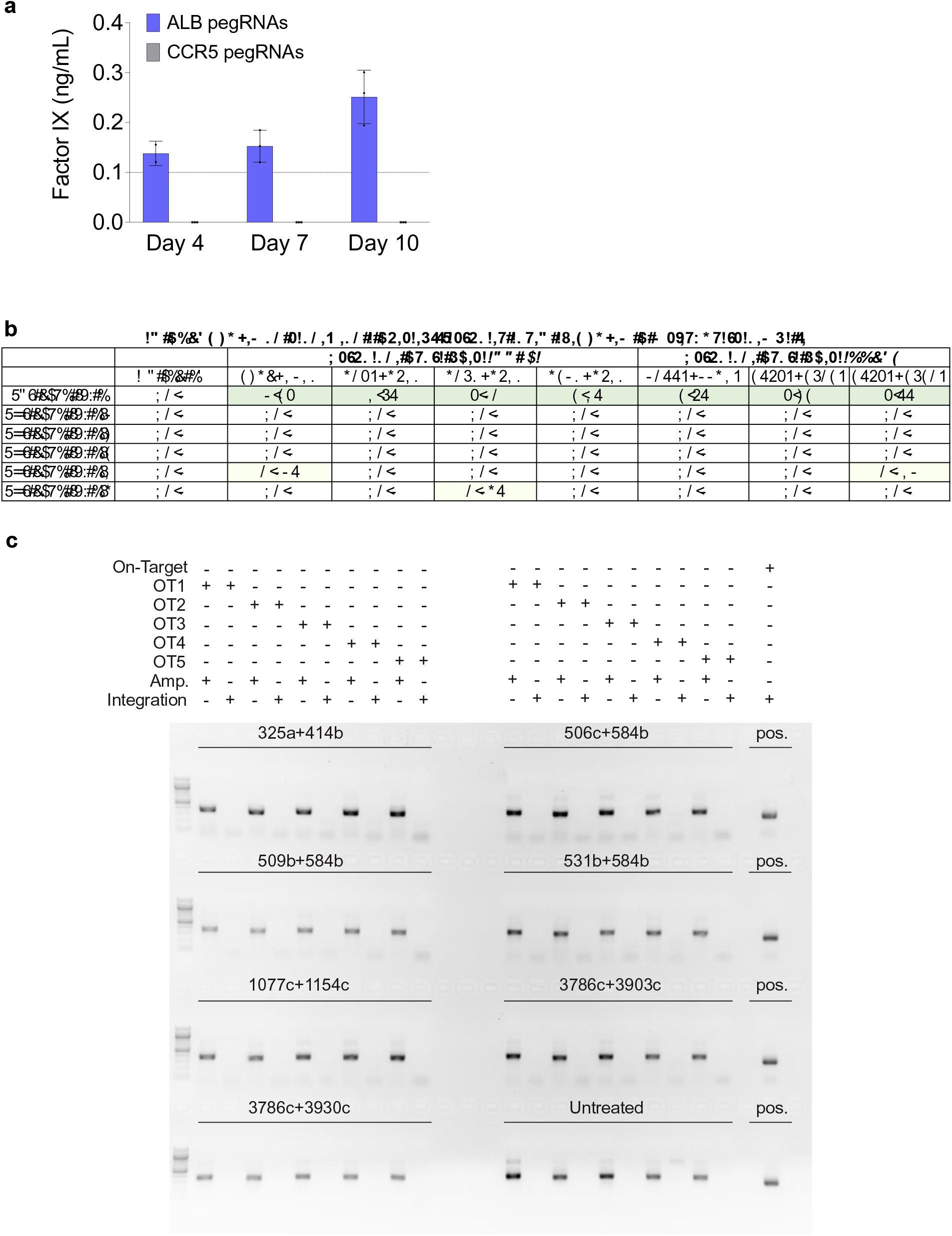
Expression of human Factor IX from the *ALB* promoter following twinPE-recombinase knock-in and characterization of Bxb1 off-target editing. (a) Huh7 cells were transfected with Bxb1, donor (*attP*-splice acceptor-cDNA of *F9* exons 2-8), PE2, and pegRNAs for installation of *attB* in the first intron of *ALB* or at *CCR5*. Three days post-transfection, cells were split and allowed to grow to confluence. Their media was changed, and they were left to condition the fresh media, with aliquots taken at days 4, 7, and 10. Factor IX was present at detectable levels by ELISA (dashed line represents the lower limit of detection) in two of three samples treated with *ALB* pegRNAs at Day 4, and in all samples treated with *ALB* pegRNAs at Day 7 and Day 10. Factor IX was never detected in the conditioned media of any samples treated with *CCR5* pegRNAs. Values and error bars reflect the mean and s.d. of two or three independent biological replicates. (**b**) Targeted amplicon sequencing was performed for each of the five nominated pseudo-sites (OT1-OT5) from seven different samples treated with 5.6-kb donor DNA plasmid, twinPE reagents targeting *CCR5 or AAVS1,* and Bxb1 recombinase. The indels in all five pseudo-sites are either below the limit of detection (<0.1%) or near-background compared to untreated controls. (**c**) To capture potential donor plasmid integration events at nominated pseudo-sites, primers were used to amplify predicted integration junctions. The gel depicts PCR reactions performed for each off-target site as indicated in the above legend. Confirmation of on-target donor integration from the samples is shown in the right-most column of the gel. In (b) and (c), two or three independent biological replicates were performed.

**Extended Data Fig. 8.**
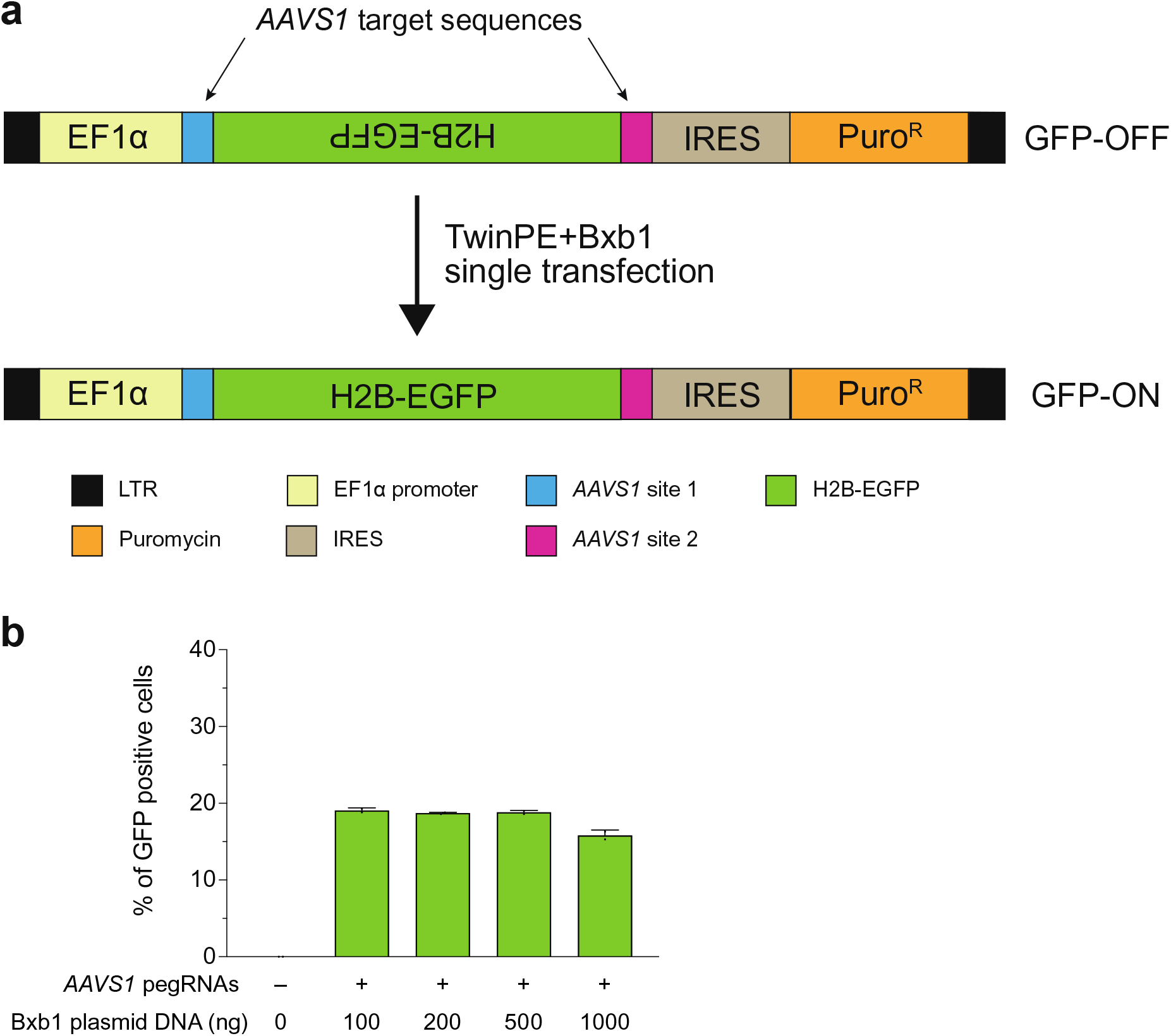
TwinPE and Bxb1-mediated inversion in HEK293T GFP reporter cells. (**a**) The lentiviral fluorescent reporter construct used to assess inversion efficiency with twinPE and Bxb1 recombinase. The reporter contains an EF1α promoter followed by an inverted H2B-EGFP coding sequence that is flanked by partial *AAVS1* DNA sequence, an internal ribosome entry site (IRES), and a puromycin resistance gene. Successful installation of opposite-facing *attB* (left) and *attP* (right) sequences at the *AAVS1* target sequences and subsequent inversion by Bxb1 corrects the orientation of GFP for functional expression. (**b**) The fluorescent reporter construct was stably integrated into HEK293T cells via lentiviral transduction and puromycin selection. The polyclonal GFP reporter cell line was then transfected with twinPE plasmid components (PE2 and four pegRNAs) and varying amounts of Bxb1 plasmid for single-transfection inversion. Cells were analyzed by flow cytometry and gated for live single cells. Quantification of GFP positive cells by flow cytometry. Values and error bars reflect the mean and s.d. of two independent biological replicates.

**Extended Data Fig. 9.**
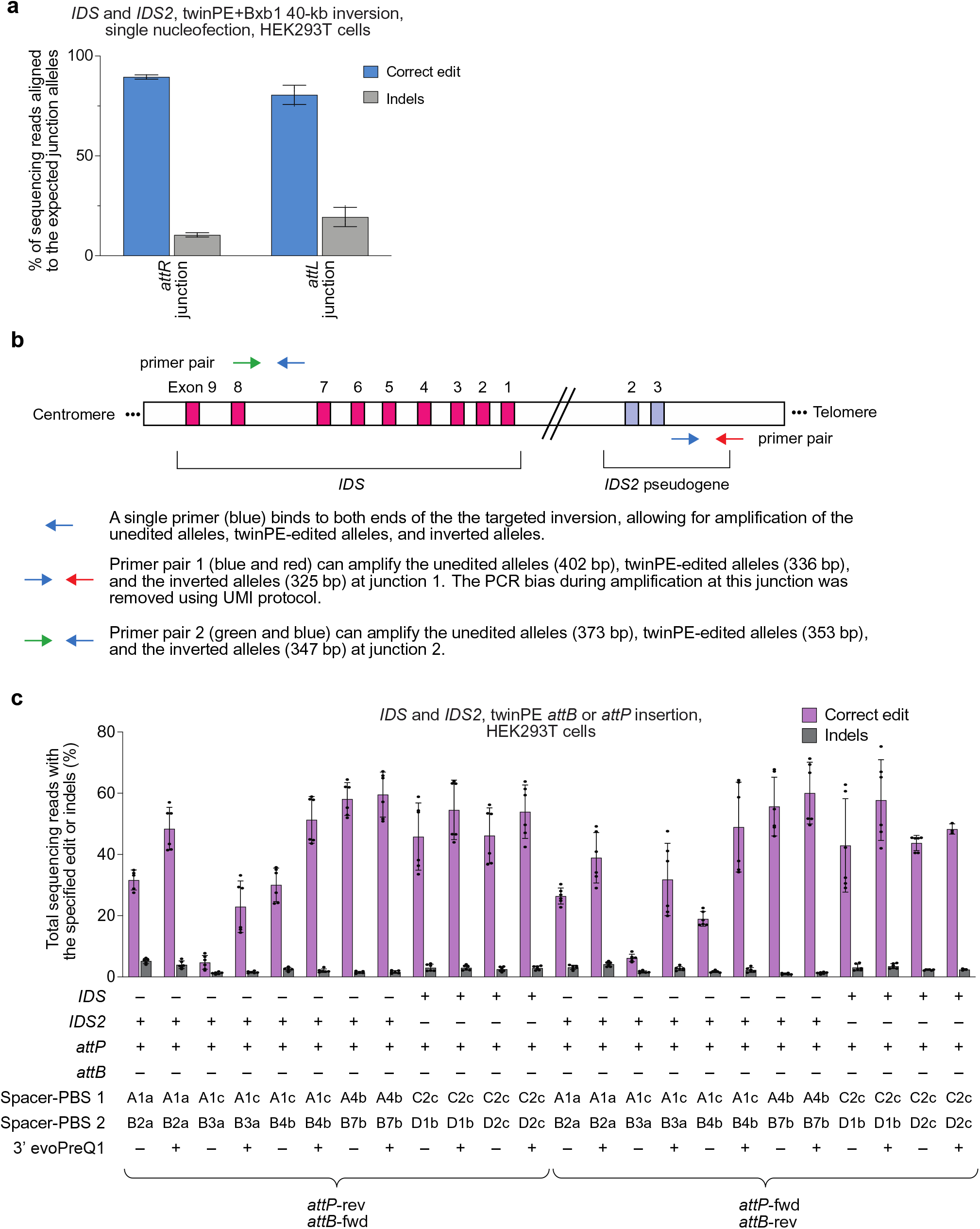
TwinPE and Bxb1 recombinase-mediated inversion between IDS and IDS2. (**a**) Assessment of the inverted *IDS* junction purity by high-throughput sequencing in HEK293T cells. Frequency of expected junction sequences containing *attR* and *attL* recombination products after twinPE and BxB1-mediated single-step inversion. The product purities range from 81-89%. Values and error bars reflect the mean and s.d. of three independent biological replicates. (**b**) Schematic diagram of the designed PCR strategies for quantifying *IDS* inversion efficiency. Primer pair 1 (green forward and blue reverse primer) can amplify the unedited alleles (403 bp), twinPE-edited alleles (337 bp), and the inverted alleles (326 bp) at junction 1 in a single PCR reaction. Due to the size difference, a UMI protocol was applied to eliminate PCR bias during quantification of inversion efficiency. Similarly, using primer pair 2 (red forward and blue reverse primer), the unedited alleles (346 bp), twinPE-edited alleles (326 bp), and inverted alleles (320 bp) at junction 2 can be amplified in a single PCR reaction. Amplicons can then be sequenced by standard high-throughput sequencing protocols for amplicon sequencing. (**c**) Screening of pegRNA pairs for the insertion of Bxb1 *attB* and *attP* sequences at *IDS* and *IDS2*. TwinPE editing was tested with standard pegRNAs and epegRNAs containing a 3’ evoPreQ1 motif. Values and error bars reflect the mean and s.d. of three independent biological replicates.

**Extended Data Fig. 10.**
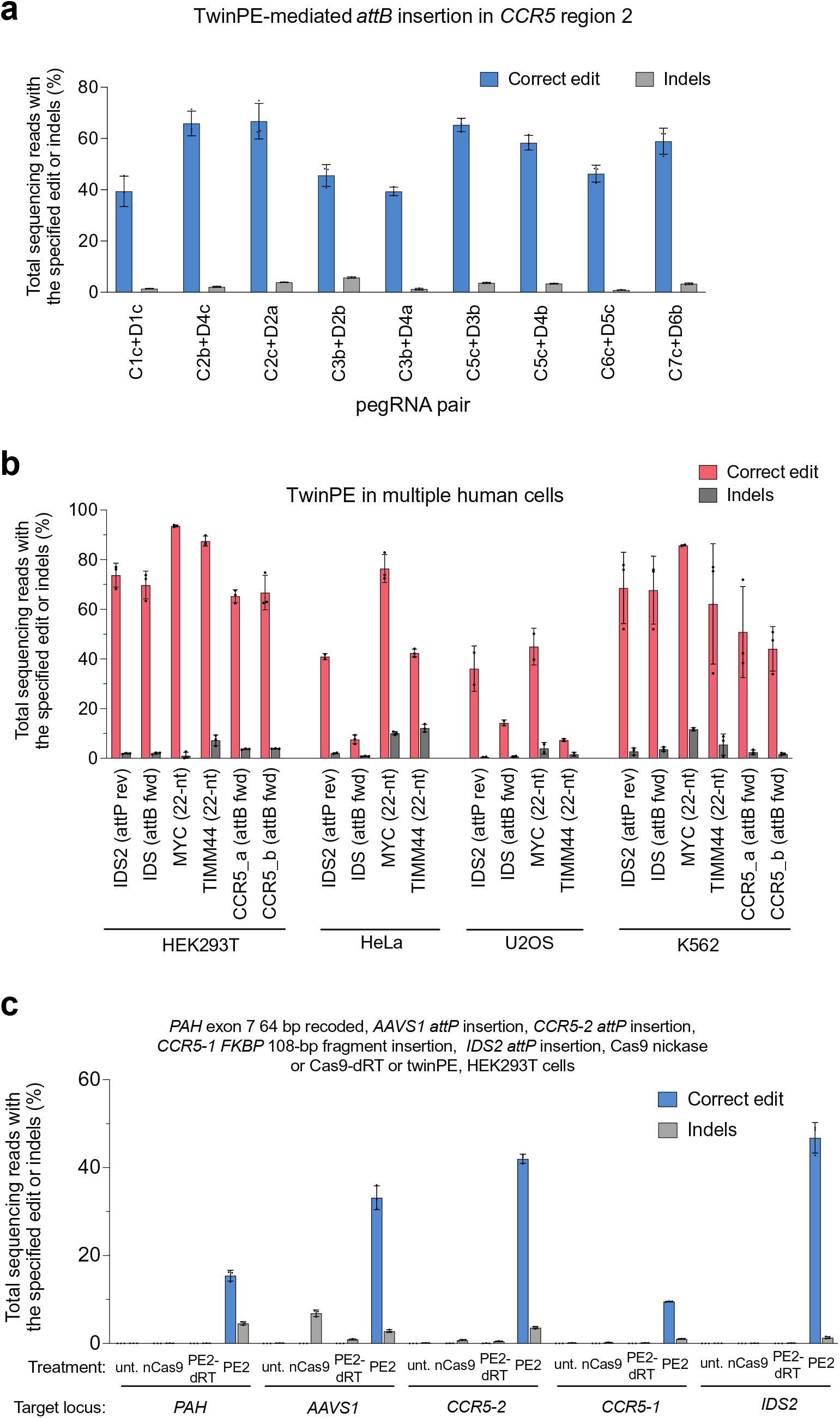
Twin prime editing mediated insertion in *CCR5* region 2 in HEK293T cells, twin prime editing in multiple human cell lines, and editing activity of Cas9 nickase and PE2-dead RT variants. (**a**) TwinPE-mediated endogenous sequence replacement with Bxb1 *attB* attachment site in *CCR5* region 2 in HEK293T cells. (**b**) TwinPE-mediated endogenous sequence replacement with *attP*, *attB,* or 22-nt DNA sequences in multiple human cell lines. Six different pegRNA pairs targeting five loci were tested in HEK293T, HeLa, U2OS and K562 cells. HEK293T and HeLa cell were transfected with PE2 and pegRNA plasmids via Lipofectamine 2000 (Thermo Fisher) and *Trans*IT-HeLaMonster (Mirus), respectively. U2OS and K562 cells were nucleofected using Lonza 4D-Nucleofector and SE kit. DNA loci and the specified insertion edits are shown in the x-axis. (**c**) HEK293T cells were transfected with twinPE pegRNA pairs and either Cas9–H840A nickase (nCas9), PE2-dRT (a PE2 variant that contains K103L and R110S inactivating mutations to the RT domain), or PE2. Treatment with either nCas9 or PE2-dRT did not result in desired edits, while PE2 installed the specified edits as indicated. Values and error bars in (a) and (c) reflect the mean and s.d. of three independent biological replicates. Values and error bars in (b) reflect the mean and s.d. of at least two independent biological replicates.

