## Supplementary material for "Programmable large DNA deletion, replacement, integration, and inversion with twin prime editing and site-specific recombinases": Complete SI

| | Page # |
|---|---|
| **Supplementary Table 1**. Sequences of pegRNAs and sgRNAs used in mammalian cell experiments | 2 |
| **Supplementary Table 2**. Sequences of primers used for mammalian cell genomic DNA amplification and HTS | 16 |
| **Supplementary Table 3**. Sequences of primers and probes used for ddPCR assays | 19 |
| **Supplementary Table 4**. Sequence of recoded exonic *PAH* sequences | 20 |
| **Supplementary Table 5**. TwinPE-mediated off-target genome editing | 21 |
| **Supplementary Note 1**. Analysis of editing quantification bias | 22 |
| **Supplementary Note 2.** Representative plot of FACS gating for GFP reporter assay | 25 |
| **Supplementary Note 3.** Analysis of twinPE design principles | 26 |
| **Supplementary Sequences 1**. Sequences of plasmid donor DNA harboring Bxb1 recombination sites | 27 |
| **Supplementary Sequence 2.** Sequence of codon-optimized Bxb1 plasmid | 35 |
| **Supplementary Sequence 3**. Sequences of the lentiviral GFP reporter vector | 37 |
| **Supplementary References** | 40 |

**Supplementary Table 1:** Sequences of pegRNAs and sgRNAs used in mammalian cell experiments

All sequences are shown in 5' to 3' orientation. pegRNAs are a concatenation of the spacer sequence, the sgRNA scaffold, and the 3' extension (contains PBS and RT template, and a 3' motif in the case of epegRNAs).

sgRNA scaffold sequence (5' to 3')
gttttagagctagaaatagcaagttaaaataaggctagtccgttatcaacttgaaaaagtggcaccgagtcggtgc

**Figure 1c**

| **pegRNA** | **spacer sequence** | **3' extension** | **Edits made by the specified pegRNA** |
|---|---|---|---|
| HEK3_attB_A_38 | ggcccagactgagcacgtga | atgatcctgacgacggagaccgccgtcgtcgacaagcccgtgctcagtctg | indicated on the x-axis |
| HEK3_attB_A_34 | ggcccagactgagcacgtga | tcctgacgacggagaccgccgtcgtcgacaagcccgtgctcagtctg | indicated on the x-axis |
| HEK3_attB_A_30 | ggcccagactgagcacgtga | gacgacggagaccgccgtcgtcgacaagcccgtgctcagtctg | indicated on the x-axis |
| HEK3_attB_B_38 | gtcaaccagtatcccggtgc | ggcttgtcgacgacggcggtctccgtcgtcaggatcatccgggatactgg | indicated on the x-axis |
| HEK3_attB_B_34 | gtcaaccagtatcccggtgc | tgtcgacgacggcggtctccgtcgtcaggatcatccgggatactgg | indicated on the x-axis |
| HEK3_attB_B_30 | gtcaaccagtatcccggtgc | gacgacggcggtctccgtcgtcaggatcatccgggatactgg | indicated on the x-axis |
| HEK3_attP_A_43 | ggcccagactgagcacgtga | taccgtacaccactgagaccgcggtggttgaccagacaaacctcgtgctcagtctg | indicated on the x-axis |
| HEK3_attP_A_39 | ggcccagactgagcacgtga | gtacaccactgagaccgcggtggttgaccagacaaacctcgtgctcagtctg | indicated on the x-axis |
| HEK3_attP_A_35 | ggcccagactgagcacgtga | accactgagaccgcggtggttgaccagacaaacctcgtgctcagtctg | indicated on the x-axis |
| HEK3_attP_B_44 | gtcaaccagtatcccggtgc | gtctggtcaaccaccgcggtctcagtggtgtacggtacaaacctccgggatactgg | indicated on the x-axis |
| HEK3_attP_B_40 | gtcaaccagtatcccggtgc | ggtcaaccaccgcggtctcagtggtgtacggtacaaacctccgggatactgg | indicated on the x-axis |
| HEK3_attP_B_36 | gtcaaccagtatcccggtgc | aaccaccgcggtctcagtggtgtacggtacaaacctccgggatactgg | indicated on the x-axis |

**Figure 2b**

| **pegRNA** | **spacer sequence** | **3' extension** | **Edits made by the specified pegRNA** |
|---|---|---|---|
| PE3_HEK3_FKBP_ins12bp | ggcccagactgagcacgtga | tggaggaagcagggcttcctttcctctgccatcacacctgcactcccgtgctcagtctg | indicated on the x-axis |
| PE3_HEK3_FKBP_ins36bp | ggcccagactgagcacgtga | tggaggaagcagggcttcctttcctctgccatcacccgtctcctggggagatggtttccacctgcactcccgtgctcagtctg | indicated on the x-axis |
| PE3_HEK3_FKBP_ins108bp | ggcccagactgagcacgtga | tggaggaagcagggcttcctttcctctgccatcaaaatttctttccatcttcaagcatcccggtgtagtgcaccacgcaggtctggccgcgcttggggaaggtgcgcccgtctcctggggagatggtttccacctgcactcccgtgctcagtctg | indicated on the x-axis |
| twinPE_HEK3_FKBP_ins108bp_A | ggcccagactgagcacgtga | accacgcaggtctggccgcgcttggggaaggtgcgcccgtctcctggggagatggtttccacctgcactcccgtgctcagtctg | indicated on the x-axis |
| twinPE_HEK3_FKBP_ins108bp_B | gtcaaccagtatcccggtgc | accttccccaagcgcggccagacctgcgtggtgcactacaccgggatgcttgaagatggaaagaaatttccgggatactgg | indicated on the x-axis |
| **sgRNA** | **spacer sequence** | | |
| HEK3_3b_+90_nicking | gtcaaccagtatcccggtgc | | |

**Figure 2c**

| **pegRNA** | **spacer sequence** | **3' extension** | **Edits made by the specified pegRNA** |
|---|---|---|---|
| PAH_E4.2_45 | gcccaagaaccattcaagagc | gttcggctccgtaggacaagatttgattagcgaacctatcaagttcttgaatggttc | exon 4 24-bp overlap (64 bp recoded) |
| PAH_E4.4_43 | gctacgggccatggactcaca | aaatcttgtcctacggagccgaacttgacgctgatcatcctgtgagtccatggcc | exon 4 24-bp overlap (64 bp recoded) |
| PAH_E4.2_45_EvoPreQ1 | gcccaagaaccattcaagagc | gttcggctccgtaggacaagatttgattagcgaacctatcaagttcttgaatggttctctctctcttgacgcggttctatctagttacgcgttaaaccaactagaaa | exon 4 24-bp overlap epegRNA (64 bp recoded) |
| PAH_E4.4_43_EvoPreQ1 | gctacgggccatggactcaca | aaatcttgtcctacggagccgaacttgacgctgatcatcctgtgagtccatggcctctctctcttgacgcggttctatctagttacgcgttaaaccaactagaaa | exon 4 24-bp overlap epegRNA (64 bp recoded) |
| PAH_E4..2_50 | gcccaagaaccattcaagagc | gtcaagttcggctccgtaggacaagatttgattagcgaacctatcaagttcttgaatggttc | exon 4 36-bp overlap (64 bp recoded) |
| PAH_E4..4_50 | gctacgggccatggactcaca | gctaatcaaatcttgtcctacggagccgaacttgacgctgatcatcctgtgagtccatggcc | exon 4 36-bp overlap (64 bp recoded) |
| PAH_E4..2_50_EvoPreQ1 | gcccaagaaccattcaagagc | gtcaagttcggctccgtaggacaagatttgattagcgaacctatcaagttcttgaatggttctctctctcttgacgcggttctatctagttacgcgttaaaccaactagaaa | exon 4 36-bp overlap epegRNA (64 bp recoded) |
| PAH_E4..4_50_EvoPreQ1 | gctacgggccatggactcaca | gctaatcaaatcttgtcctacggagccgaacttgacgctgatcatcctgtgagtccatggcctctctctcttgacgcggttctatctagttacgcgttaaaccaactagaaa | exon 4 36-bp overlap epegRNA (64 bp recoded) |
| PAH_E4.2_62 | gcccaagaaccattcaagagc | aggatgatcagcgtcaagttcggctccgtaggacaagatttgattagcgaacctatcaagttcttgaatggttc | exon 4 59-bp overlap (64 bp recoded) |
| PAH_E4.4_61 | gctacgggccatggactcaca | ttgataggttcgctaatcaaatcttgtcctacggagccgaacttgacgctgatcatcctgtgagtccatggcc | exon 4 59-bp overlap (64 bp recoded) |
| PAH_E4.2_62_EvoPreQ1 | gcccaagaaccattcaagagc | aggatgatcagcgtcaagttcggctccgtaggacaagatttgattagcgaacctatcaagttcttgaatggttctctctctcttgacgcggttctatctagttacgcgttaaaccaactagaaa | exon 4 59-bp overlap epegRNA (64 bp recoded) |
| PAH_E4.4_61_EvoPreQ1 | gctacgggccatggactcaca | ttgataggttcgctaatcaaatcttgtcctacggagccgaacttgacgctgatcatcctgtgagtccatggcctctctctcttgacgcggttctatctagttacgcgttaaaccaactagaaa | exon 4 59-bp overlap epegRNA (64 bp recoded) |

| | | | |
|---|---|---|---|
| PAH_E7.2_34 | gtggtttccgcctccgacctg | agaaagtctctactgctcaagagcccggcaacgggtcggaggcg | exon 7 22-bp overlap (46 bp recoded) |
| PAH_E7.5_34 | gagtggaagactcggaaggcc | tcttgagcagtagagactttctggggggtctcgccttccgagtcttc | exon 7 22-bp overlap (46 bp recoded) |
| PAH_E7.2_34_EvoPreQ1 | gtggtttccgcctccgacctg | agaaagtctctactgctcaagagcccggcaacgggtcggaggcgtctctcttgacgcggttctatctagttacgcgttaaaccaactagaaa | exon 7 22-bp overlap epegRNA (46 bp recoded) |
| PAH_E7.5_34_EvoPreQ1 | gagtggaagactcggaaggcc | tcttgagcagtagagactttctggggggtctcgccttccgagtcttctctctctcttgacgcggttctatctagttacgcgttaaaccaactagaaa | exon 7 22-bp overlap epegRNA (46 bp recoded) |
| PAH_E7.2_44 | gtggtttccgcctccgacctg | gagaccccccagaaagtctctactgctcaagagcccggcaacgggtcggaggcg | exon 7 42-bp overlap (46 bp recoded), exon 7 24-bp overlap (64 bp recoded) |
| PAH_E7.5_44 | gagtggaagactcggaaggcc | gttgccgggctcttgagcagtagagactttctggggggtctcgccttccgagtcttc | exon 7 42-bp overlap (46 bp recoded) |
| PAH_E7.2_44_EvoPreQ1 | gtggtttccgcctccgacctg | gagaccccccagaaagtctctactgctcaagagcccggcaacgggtcggaggcgtctctctcttgacgcggttctatctagttacgcgttaaaccaactagaaa | exon 7 42-bp overlap epegRNA (46 bp recoded), exon 7 24-bp overlap epegRNA (64 bp recoded) |
| PAH_E7.5_44_EvoPreQ1 | gagtggaagactcggaaggcc | gttgccgggctcttgagcagtagagactttctggggggtctcgccttccgagtcttctctctctcttgacgcggttctatctagttacgcgttaaaccaactagaaa | exon 7 42-bp overlap epegRNA (46 bp recoded) |
| PAH_E7.6_44 | gtctgatgtactgtgtgcag | agtagagactttctggggggtctcgcattccgcgtgtttcattgcacacagtaca | exon 7 24-bp overlap (64 bp recoded) |
| PAH_E7.6_44_EvoPreQ1 | gtctgatgtactgtgtgcag | agtagagactttctggggggtctcgcattccgcgtgtttcattgcacacagtacatctctctcttgacgcggttctatctagttacgcgttaaaccaactagaaa | exon 7 24-bp overlap epegRNA (64 bp recoded) |
| PAH_E7.2_55 | gtggtttccgcctccgacctg | acgcggaatgcgagaccccccagaaagtctctactgctcaagagcccggcaacgggtcggaggcg | exon 7 47-bp overlap (64 bp recoded) |
| PAH_E7.6_56 | gtctgatgtactgtgtgcag | gggctcttgagcagtagagactttctggggggtctcgcattccgcgtgtttcattgcacacagtaca | exon 7 47-bp overlap (64 bp recoded) |
| PAH_E7.2_55_EvoPreQ1 | gtggtttccgcctccgacctg | acgcggaatgcgagaccccccagaaagtctctactgctcaagagcccggcaacgggtcggaggcgtctctctcttgacgcggttctatctagttacgcgttaaaccaactagaaa | exon 7 47-bp overlap epegRNA (64 bp recoded) |
| PAH_E7.6_56_EvoPreQ1 | gtctgatgtactgtgtgcag | gggctcttgagcagtagagactttctggggggtctcgcattccgcgtgtttcattgcacacagtacatctctctcttgacgcggttctatctagttacgcgttaaaccaactagaaa | exon 7 47-bp overlap epegRNA (64 bp recoded) |

**Figure 2e**

| pegRNA | spacer sequence | 3' extension | Edits made by the specified pegRNA |
|---|---|---|---|
| HEK3_DF_A_SA_del77nt | ggcccagactgagcacgtga | tcctctgccatcacgtgctcagtctg | SA (Δ 77nt) |
| HEK3_DF_B_SA_del77nt | gtcaaccagtatcccggtgc | tgatggcagaggaccgggatactgg | SA (Δ 77nt) |
| HEK3_DF_A_SA_del56nt | ggcccagactgagcacgtga | tggaggaagcagggcttcctttcctctgccatcacgtgctcagtctg | SA (Δ 56nt) |
| HEK3_DF_B_SA_del56nt | gtcaaccagtatcccggtgc | tgatggcagaggaaaggaagccctgcttcctccaccgggatactgg | SA (Δ 56nt) |
| HEK3_DF_A_HA_del64nt | ggcccagactgagcacgtga | tgcaggagctgcatcctctgccatcacgtgctcagtctg | HA (Δ 64nt) |
| HEK3_DF_B_HA_del64nt | gtcaaccagtatcccggtgc | tgatggcagaggatgcagctcctgcaccgggatactgg | HA (Δ 64nt) |
| HEK3_DF_A_PD_del90nt | ggcccagactgagcacgtga | gcccagccaaacttgtcaaccagtatcccggcgtgctcagtctg | PD (Δ 90nt) |
| HEK3_DF_B_PD_del90nt | gtcaaccagtatcccggtgc | gggtcaatccttggggcccagactgagcacgccgggatactgg | PD (Δ 90nt) |
| HEK3_DF_A_SA_del77nt_EvoPreQ1 | ggcccagactgagcacgtga | tcctctgccatcacgtgctcagtctgttaaataacgcggttctatctagttacgcgttaaaccaactagaa | SA-EvoPreQ1 (Δ 77nt) |
| HEK3_DF_B_SA_del77nt_EvoPreQ1 | gtcaaccagtatcccggtgc | tgatggcagaggaccgggatactggaaaatatacgcggttctatctagttacgcgttaaaccaactagaa | SA-EvoPreQ1 (Δ 77nt) |
| HEK3_DF_A_SA_del56nt_EvoPreQ1 | ggcccagactgagcacgtga | tggaggaagcagggcttcctttcctctgccatcacgtgctcagtctgttaataatcgcggttctatctagttacgcgttaaaccaactagaa | SA-EvoPreQ1 (Δ 56nt) |
| HEK3_DF_B_SA_del56nt_EvoPreQ1 | gtcaaccagtatcccggtgc | tgatggcagaggaaaggaagccctgcttcctccaccgggatactggaaaaaaaaacgcggttctatctagttacgcgttaaaccaactagaa | SA-EvoPreQ1 (Δ 56nt) |
| HEK3_DF_A_HA_del64nt_EvoPreQ1 | ggcccagactgagcacgtga | tgcaggagctgcatcctctgccatcacgtgctcagtctgataaataacgcggttctatctagttacgcgttaaaccaactagaa | SA-EvoPreQ1 (Δ 64nt) |
| HEK3_DF_B_HA_del64nt_EvoPreQ1 | gtcaaccagtatcccggtgc | tgatggcagaggatgcagctcctgcaccgggatactggaaaaaaggcgcggttctatctagttacgcgttaaaccaactagaa | SA-EvoPreQ1 (Δ 64nt) |
| HEK3_DF_A_PD_del90nt_EvoPreQ1 | ggcccagactgagcacgtga | gcccagccaaacttgtcaaccagtatcccggcgtgctcagtctgttaaataccgcggttctatctagttacgcgttaaaccaactagaa | PD-EvoPreQ1 (Δ 90nt) |
| HEK3_DF_B_PD_del90nt_EvoPreQ1 | gtcaaccagtatcccggtgc | gggtcaatccttggggcccagactgagcacgccgggatactggaaaataatcgcggttctatctagttacgcgttaaaccaactagaa | PD-EvoPreQ1 (Δ 90nt) |

**Figure 2f**

| pegRNA | spacer sequence | 3' extension | Edits made by the specified pegRNA |
|---|---|---|---|
| DMD-Exon51-A1_a_SA | gattggctttgatttcccta | ttgaataggaagtaaattaatttgaagctggaccctagggaaatcaa | SA-1 (Δ 780nt) |
| DMD-Exon51-B1_A1_b_SA | gcagttgcctaagaactggt | tagggtccagcttcaaattaatttacttcctattcaaagttcttaggc | SA-1 (Δ 780nt) |
| DMD-Exon51-a3-b1_a_PD | gtatatgattgttactgaga | tgctgagagagaaacagttgcctaagaactcagtaacaat | PD-1 (Δ 627nt); PD-2 (Δ 627nt) |

| | | | |
|---|---|---|---|
| DMD-Exon51-b1-a3_a-PD | gcagttgcctaagaactggt | accacttccacaatgtatatgattgttactgagttcttagg | PD-1 (Δ 627nt) |
| DMD-Exon51-b1-a3_b-PD | gcagttgcctaagaactggt | accacttccacaatgtatatgattgttactgagttcttaggcaa | PD-2 (Δ 627nt) |
| DMD-Exon51-A3_b_attB | gtatatgattgttactgaga | atgatcctgacgacggagaccgccgtcgtcgacaagcccagtaacaatc | tPE-1 attB (Δ 589nt); tPE-2 attB (Δ 589nt); tPE-1 attB (Δ 558nt); tPE-2 attB (Δ 558nt) |
| DMD-Exon51-A3_c_attB | gtatatgattgttactgaga | atgatcctgacgacggagaccgccgtcgtcgacaagcccagtaacaatcata | tPE-3 attB (Δ 589nt); tPE-4 attB (Δ 589nt); tPE-3 attB (Δ 558nt); tPE-4 attB (Δ 558nt) |
| DMD-Exon51-B1_b_attB | gcagttgcctaagaactggt | ggcttgtcgacgacggcggtctccgtcgtcaggatcatagttcttaggc | tPE-1 attB (Δ 589nt); tPE-3 attB (Δ 589nt) |
| DMD-Exon51-B1_c_attB | gcagttgcctaagaactggt | ggcttgtcgacgacggcggtctccgtcgtcaggatcatagttcttaggcaac | tPE-2 attB (Δ 589nt); tPE-4 attB (Δ 589nt) |
| DMD-Exon51-B2_b_attB | gaggagagtaaagtgattgg | ggcttgtcgacgacggcggtctccgtcgtcaggatcatatcactttactc | tPE-1 attB (Δ 558nt); tPE-3 attB (Δ 558nt) |
| DMD-Exon51-B2_c_attB | gaggagagtaaagtgattgg | ggcttgtcgacgacggcggtctccgtcgtcaggatcatatcactttactctcc | tPE-2 attB (Δ 558nt); tPE-4 attB (Δ 558nt) |
| **sgRNA** | **spacer sequence** | | **Edits made by the paired sgRNA** |
| DMD-Exon51-A1 | gattggctttgatttcccta | | Cas9 (Δ 818nt) |
| DMD-Exon51-A3 | gtatatgattgttactgaga | | Cas9 (Δ 627nt) |
| DMD-Exon51-B1 | gcagttgcctaagaactggt | | Cas9 (Δ 818nt); Cas9 (Δ 627nt) |

**Figure 3b**

| pegRNA | spacer sequence | 3' extension | Edits made by the specified pegRNA |
|---|---|---|---|
| CCR5_A223c | gtcatcctgataaactgcaaa | atgatcctgacgacggagaccgccgtcgtcgacaagccgcagtttatcagg | indicated on the x-axis |
| CCR5_B272a | gaaggaaaaacaggtcagaga | ggcttgtcgacgacggcggtctccgtcgtcaggatcatctgacctg | indicated on the x-axis |
| CCR5_B291b | gcccagaaggggacagtaaga | ggcttgtcgacgacggcggtctccgtcgtcaggatcattactgtcccct | indicated on the x-axis |
| CCR5_B305a | ggcagcatagtgagcccaga | ggcttgtcgacgacggcggtctccgtcgtcaggatcatgggctcact | indicated on the x-axis |
| CCR5_B326b | gatttccaaagtcccactggg | ggcttgtcgacgacggcggtctccgtcgtcaggatcatagtgggactttg | indicated on the x-axis |
| CCR5_B330b | gttgtatttccaaagtcccac | ggcttgtcgacgacggcggtctccgtcgtcaggatcatggactttggaa | indicated on the x-axis |
| CCR5_A260c | gtgacatctacctgctcaacc | atgatcctgacgacggagaccgccgtcgtcgacaagcctgagcaggtagat | indicated on the x-axis |
| CCR5_B305c | ggcagcatagtgagcccaga | ggcttgtcgacgacggcggtctccgtcgtcaggatcatgggctcactatgc | indicated on the x-axis |
| CCR5_B330c | gttgtatttccaaagtcccac | ggcttgtcgacgacggcggtctccgtcgtcaggatcatggactttggaaat | indicated on the x-axis |
| CCR5_A325b | gctcactatgctgccgcccag | atgatcctgacgacggagaccgccgtcgtcgacaagcccggcggcagcat | indicated on the x-axis |
| CCR5_B414a | ggtacctatcgattgtcagg | ggcttgtcgacgacggcggtctccgtcgtcaggatcatgacaatcga | indicated on the x-axis |
| CCR5_A360b | gacaatgtgtcaactcttgac | atgatcctgacgacggagaccgccgtcgtcgacaagccaagagttgacac | indicated on the x-axis |
| CCR5_A506c | gacaagtgtgatcacttggg | atgatcctgacgacggagaccgccgtcgtcgacaagccaagtgatcacactt | indicated on the x-axis |
| CCR5_B584a | gtatggaaaatgagagctgc | ggcttgtcgacgacggcggtctccgtcgtcaggatcatgctctcatt | indicated on the x-axis |
| CCR5_A509a | gaagtgtgatcacttgggtgg | atgatcctgacgacggagaccgccgtcgtcgacaagcccccaagtga | indicated on the x-axis |
| CCR5_B535c | gatctggtaaagatgattcc | ggcttgtcgacgacggcggtctccgtcgtcaggatcatatcatctttaccag | indicated on the x-axis |
| CCR5_A531c | gctgtgtttgcgtctctccc | atgatcctgacgacggagaccgccgtcgtcgacaagccagagacgcaaaca | indicated on the x-axis |
| CCR5_B584b | gtatggaaaatgagagctgc | ggcttgtcgacgacggcggtctccgtcgtcaggatcatgctctcattttc | indicated on the x-axis |

**Figure 3c**

| pegRNA | spacer sequence | 3' extension | Edits made by the specified pegRNA |
|---|---|---|---|
| AAVS1_A1077b | gcagagccaggaacccctgt | taccgtacaccactgagaccgcggtggttgaccagacaaacctggggttcctgg | indicated on the x-axis |
| AAVS1_B1154b | gtccttggcaagcccaggag | gtctggtcaaccaccgcggtctcagtggtgtacggtacaaacctctgggcttgcc | indicated on the x-axis |
| AAVS1_A1098b | gggaaggggcaggagagcca | taccgtacaccactgagaccgcggtggttgaccagacaaacctctctcctgccc | indicated on the x-axis |
| AAVS1_B1154a | gtccttggcaagcccaggag | gtctggtcaaccaccgcggtctcagtggtgtacggtacaaacctctgggcttg | indicated on the x-axis |
| AAVS1_A1246c | gaatatgtcccagatagcac | taccgtacaccactgagaccgcggtggttgaccagacaaacctctatctgggacat | indicated on the x-axis |
| AAVS1_B1314b | gtgcgtcctaggtgttcacc | gtctggtcaaccaccgcggtctcagtggtgtacggtacaaacctgaacacctagg | indicated on the x-axis |
| AAVS1_B1376a | gtcctggcagggctgtggtg | gtctggtcaaccaccgcggtctcagtggtgtacggtacaaacctcacagccct | indicated on the x-axis |
| AAVS1_A1267c | ggggactctttaaggaaaga | taccgtacaccactgagaccgcggtggttgaccagacaaacctttccttaaagagt | indicated on the x-axis |
| AAVS1_A1293a | gagaaagagaaagggagtag | taccgtacaccactgagaccgcggtggttgaccagacaaacctctccctttc | indicated on the x-axis |
| AAVS1_A1307b | gagtagaggcggccacgacc | taccgtacaccactgagaccgcggtggttgaccagacaaacctcgtggccgcct | indicated on the x-axis |
| AAVS1_B1314a | gtgcgtcctaggtgttcacc | gtctggtcaaccaccgcggtctcagtggtgtacggtacaaacctgaacaccta | indicated on the x-axis |
| AAVS1_A1582c | gatcagtgaaacgcaccaga | taccgtacaccactgagaccgcggtggttgaccagacaaacctggtgcgtttcact | indicated on the x-axis |
| AAVS1_B1640a | gtgacctgcccggttctcag | gtctggtcaaccaccgcggtctcagtggtgtacggtacaaacctagaaccggg | indicated on the x-axis |
| AAVS1_B1676a | gagcttggcagggggtggga | gtctggtcaaccaccgcggtctcagtggtgtacggtacaaacctcaccccctg | indicated on the x-axis |
| AAVS1_B1701a | gagccagagaggatcctggg | gtctggtcaaccaccgcggtctcagtggtgtacggtacaaacctaggatcctc | indicated on the x-axis |
| AAVS1_B1705b | gatggagccagagaggatcc | gtctggtcaaccaccgcggtctcagtggtgtacggtacaaaccttcctctctggc | indicated on the x-axis |
| AAVS1_A1615b | gcagctcaggttctgggaga | taccgtacaccactgagaccgcggtggttgaccagacaaacctcccagaacctg | indicated on the x-axis |
| AAVS1_B1640a | gtgacctgcccggttctcag | gtctggtcaaccaccgcggtctcagtggtgtacggtacaaacctagaaccggg | indicated on the x-axis |
| AAVS1_A1615a | gcagctcaggttctgggaga | taccgtacaccactgagaccgcggtggttgaccagacaaacctcccagaacc | indicated on the x-axis |
| AAVS1_B1676a | gagcttggcagggggtggga | gtctggtcaaccaccgcggtctcagtggtgtacggtacaaacctcaccccctg | indicated on the x-axis |
| AAVS1_A1647b | gtggccactgagaaccgggc | taccgtacaccactgagaccgcggtggttgaccagacaaacctcggttctcagt | indicated on the x-axis |
| AAVS1_B1676b | gagcttggcagggggtggga | gtctggtcaaccaccgcggtctcagtggtgtacggtacaaacctcaccccctgcc | indicated on the x-axis |
| AAVS1_B1701a | gagccagagaggatcctggg | gtctggtcaaccaccgcggtctcagtggtgtacggtacaaacctaggatcctc | indicated on the x-axis |
| AAVS1_B1705a | gatggagccagagaggatcc | gtctggtcaaccaccgcggtctcagtggtgtacggtacaaaccttcctctctg | indicated on the x-axis |
| AAVS1_A1810b | gaatctgcctaacaggaggt | taccgtacaccactgagaccgcggtggttgaccagacaaaccttcctgttaggc | indicated on the x-axis |
| AAVS1_B1883b | ggggccactagggacaggat | gtctggtcaaccaccgcggtctcagtggtgtacggtacaaacctctgtccctagt | indicated on the x-axis |

| | | | |
|---|---|---|---|
| AAVS1_B1902a | gtcccctccaccccacagtg | gtctggtcaaccaccgcggtctcagtggtgtacggtacaaaccttgtggggtg | indicated on the x-axis |
| AAVS1_A1890b | gtcaccaatcctgtccctag | taccgtacaccactgagaccgcggtggttgaccagacaaacctgggacaggatt | indicated on the x-axis |
| AAVS1_A1890c | gtcaccaatcctgtccctag | taccgtacaccactgagaccgcggtggttgaccagacaaacctgggacaggattgg | indicated on the x-axis |
| AAVS1_B1962c | gggaccaccttatattccca | tctggtcaaccaccgcggtctcagtggtgtacggtacaaacctgaatataaggtgg | indicated on the x-axis |
| AAVS1_A3786c | gctggcccccaccgcccca | taccgtacaccactgagaccgcggtggttgaccagacaaacctggcggtggggggc | indicated on the x-axis |
| AAVS1_B3839c | gacctgcccagcacaccctg | tctggtcaaccaccgcggtctcagtggtgtacggtacaaacctggtgtgctgggca | indicated on the x-axis |
| AAVS1_A3786a | gctggcccccaccgcccca | taccgtacaccactgagaccgcggtggttgaccagacaaacctggcggtggg | indicated on the x-axis |
| AAVS1_B3903b | gcgactcctggaagtggcca | gtctggtcaaccaccgcggtctcagtggtgtacggtacaaacctccacttccagg | indicated on the x-axis |
| AAVS1_B3930a | ggacttcccagtgtgcatcg | gtctggtcaaccaccgcggtctcagtggtgtacggtacaaaccttgcacactg | indicated on the x-axis |
| AAVS1_A3835b | gacgtcacggcgctgcccca | taccgtacaccactgagaccgcggtggttgaccagacaaacctggcagcgccgt | indicated on the x-axis |
| AAVS1_B3930b | ggacttcccagtgtgcatcg | gtctggtcaaccaccgcggtctcagtggtgtacggtacaaaccttgcacactggg | indicated on the x-axis |
| AAVS1_A3856a | ggtgtgctgggcaggtcgcg | taccgtacaccactgagaccgcggtggttgaccagacaaacctgacctgccc | indicated on the x-axis |

**Figure 3d**

| **pegRNA** | **spacer sequence** | **3' extension** | **Edits made by the specified pegRNA** |
|---|---|---|---|
| CCR5_A325a | gctcactatgctgccgcccag | atgatcctgacgacggagaccgccgtcgtcgacaagccggcggcagc | 325/414 |
| CCR5_B414b | ggtacctatcgattgtcagg | ggcttgtcgacgacggcggtctccgtcgtcaggatcatgacaatcgata | 325/414 |
| CCR5_A506c | gacaagtgtgatcacttggg | atgatcctgacgacggagaccgccgtcgtcgacaagccaagtgatcacactt | 506/584 |
| CCR5_A509b | gaagtgtgatcacttgggtgg | atgatcctgacgacggagaccgccgtcgtcgacaagcccccaagtgatc | 509/584 |
| CCR5_A531b | gctgtgtttgcgtctctccc | atgatcctgacgacggagaccgccgtcgtcgacaagccagagacgcaaa | 531/584 |
| CCR5_B584b | gtatggaaaatgagagctgc | ggcttgtcgacgacggcggtctccgtcgtcaggatcatgctctcattttc | 506/584, 509/584, 531/584 |
| AAVS1_A1077c | gcagagccaggaacccctgt | taccgtacaccactgagaccgcggtggttgaccagacaaacctggggttcctggct | 1077/1154 |
| AAVS1_B1154c | gtccttggcaagcccaggag | tctggtcaaccaccgcggtctcagtggtgtacggtacaaacctctgggcttgccaa | 1077/1154 |
| AAVS1_A3786c | gctggcccccaccgcccca | taccgtacaccactgagaccgcggtggttgaccagacaaacctggcggtggggggc | 3786/3903, 3786/3930 |
| AAVS1_B3903c | gcgactcctggaagtggcca | tctggtcaaccaccgcggtctcagtggtgtacggtacaaacctccacttccaggag | 3786/3903 |
| AAVS1_B3930c | ggacttcccagtgtgcatcg | tctggtcaaccaccgcggtctcagtggtgtacggtacaaaccttgcacactgggaa | 3786/3930 |

**Figure 3e**

| **pegRNA** | **spacer sequence** | **3' extension** | **Edits made by the specified pegRNA** |
|---|---|---|---|
| CCR5_A531b | gctgtgtttgcgtctctccc | atgatcctgacgacggagaccgccgtcgtcgacaagccagagacgcaaa | attB 38 |
| CCR5_B584b | gtatggaaaatgagagctgc | ggcttgtcgacgacggcggtctccgtcgtcaggatcatgctctcattttc | attB 38 |
| CCR5_A7_attB_30 | gctgtgtttgcgtctctccc | tcctgacgacggagaccgccgtcgtcgacaagccagagacgcaaa | attB 30 |
| CCR5_B8_attB_30 | gtatggaaaatgagagctgc | tgtcgacgacggcggtctccgtcgtcaggatcatgctctcattttc | attB 30 |
| CCR5_A7_attB_20 | gctgtgtttgcgtctctccc | acgacggagaccgccgtcgtcgacaagccagagacgcaaa | attB 20 |
| CCR5_B8_attB_20 | gtatggaaaatgagagctgc | acgacggcggtctccgtcgtcaggatcatgctctcattttc | attB 20 |
| CCR5_A7_attB_GA_38 | gctgtgtttgcgtctctccc | atgatcctgacgacggagtccgccgtcgtcgacaagccagagacgcaaa | attB-GA 38 |
| CCR5_B8_attB_GA_38 | gtatggaaaatgagagctgc | ggcttgtcgacgacggcggactccgtcgtcaggatcatgctctcattttc | attB-GA 38 |
| CCR5_A7_attB_GA_30 | gctgtgtttgcgtctctccc | tcctgacgacggagtccgccgtcgtcgacaagccagagacgcaaa | attB-GA 30 |
| CCR5_B8_attB_GA_30 | gtatggaaaatgagagctgc | tgtcgacgacggcggactccgtcgtcaggatcatgctctcattttc | attB-GA 30 |
| CCR5_A7_attB_GA_20 | gctgtgtttgcgtctctccc | acgacggagtccgccgtcgtcgacaagccagagacgcaaa | attB-GA 20 |
| CCR5_B8_attB_GA_20 | gtatggaaaatgagagctgc | acgacggcggactccgtcgtcaggatcatgctctcattttc | attB-GA 20 |
| CCR5_A7_attP_50 | gctgtgtttgcgtctctccc | gggtttgtaccgtacaccactgagaccgcggtggttgaccagacaaaccaagagacg caaa | attP 50 |
| CCR5_B8_attP_50 | gtatggaaaatgagagctgc | tggtttgtctggtcaaccaccgcggtctcagtggtgtacggtacaaacccgctctcatttt c | attP 50 |
| CCR5_A7_attP_40 | gctgtgtttgcgtctctccc | tgtaccgtacaccactgagaccgcggtggttgaccagacaaaccaagagacgcaaa | attP 40 |
| CCR5_B8_attP_40 | gtatggaaaatgagagctgc | tgtctggtcaaccaccgcggtctcagtggtgtacggtacaaacccgctctcattttc | attP 40 |
| CCR5_A7_attP_30 | gctgtgtttgcgtctctccc | cgtacaccactgagaccgcggtggttgaccagacaaaccaagagacgcaaa | attP 30 |
| CCR5_B8_attP_30 | gtatggaaaatgagagctgc | ggtcaaccaccgcggtctcagtggtgtacggtacaaacccgctctcattttc | attP 30 |
| CCR5_A7_attP_GA_50 | gctgtgtttgcgtctctccc | gggtttgtaccgtacaccactgagtccgcggtggttgaccagacaaaccaagagacg caaa | attP-GA 50 |
| CCR5_B8_attP_GA_50 | gtatggaaaatgagagctgc | tggtttgtctggtcaaccaccgcggactcagtggtgtacggtacaaacccgctctcattt tc | attP-GA 50 |
| CCR5_A7_attP_GA_40 | gctgtgtttgcgtctctccc | tgtaccgtacaccactgagtccgcggtggttgaccagacaaaccaagagacgcaaa | attP-GA 40 |
| CCR5_B8_attP_GA_40 | gtatggaaaatgagagctgc | tgtctggtcaaccaccgcggactcagtggtgtacggtacaaacccgctctcattttc | attP-GA 40 |
| CCR5_A7_attP_GA_30 | gctgtgtttgcgtctctccc | cgtacaccactgagtccgcggtggttgaccagacaaaccaagagacgcaaa | attP-GA 30 |
| CCR5_B8_attP_GA_30 | gtatggaaaatgagagctgc | ggtcaaccaccgcggactcagtggtgtacggtacaaacccgctctcattttc | attP-GA 30 |

**Figure 3f**

| **pegRNA** | **spacer sequence** | **3' extension** | **Edits made by the specified pegRNA** |
|---|---|---|---|
| CCR5_A277b | gactgaaacttcacagaata | aacttcacagaggcttgtcgacgacggcggtctccgtcgtcaggatcat | indicated on the x-axis |
| CCR5_A277c | gactgaaacttcacagaata | gaaacttcacagaggcttgtcgacgacggcggtctccgtcgtcaggatcat | indicated on the x-axis |
| CCR5_B358b | gatttatgagatcaacagcac | ggcttgtcgacgacggcggtctccgtcgtcaggatcatctgttgatctc | indicated on the x-axis |
| CCR5_B358c | gatttatgagatcaacagcac | ggcttgtcgacgacggcggtctccgtcgtcaggatcatctgttgatctcat | indicated on the x-axis |

**Figure 3g**

| **pegRNA** | **spacer sequence** | **3' extension** | **Edits made by the specified pegRNA** |
|---|---|---|---|
| CCR5_A7_attB_20 | gctgtgtttgcgtctctccc | acgacggagaccgccgtcgtcgacaagccagagacgcaaa | CCR5 attB (HEK293T and Huh7) |
| CCR5_B8_attB_20 | gtatggaaaatgagagctgc | acgacggcggtctccgtcgtcaggatcatgctctcattttc | CCR5 attB (HEK293T and Huh7) |
| CCR5_A277c | gactgaaacttcacagaata | gaaacttcacagaggcttgtcgacgacggcggtctccgtcgtcaggatcat | ALB attB (HEK293T and Huh7) |

| | | | |
|---|---|---|---|
| CCR5_B358c | gatttatgagatcaacagcac | ggcttgtcgacgacggcggtctccgtcgtcaggatcatctgttgatctcat | ALB attB (HEK293T and Huh7) |

**Figure 4b**

| **pegRNA** | **spacer sequence** | **3' extension** | **Edits made by the specified pegRNA** |
|---|---|---|---|
| IDS2_DF_A1_a_attP_rev | gacaccaaaaactgccacagg | taccgtacaccactgagaccgcggtggttgaccagacaaacctgtggcagtta | A1B2 (attP rev) |
| IDS2_DF_A1_a_ attP_fwd | gacaccaaaaactgccacagg | gtctggtcaaccaccgcggtctcagtggtgtacggtacaaacctgtggcagtta | A1B2 (attP fwd) |
| IDS2_DF_A1_c_attP_rev | gacaccaaaaactgccacagg | taccgtacaccactgagaccgcggtggttgaccagacaaacctgtggcagtttta | A1B3 (attP rev); A1B4 (attP rev) |
| IDS2_DF_A1_c_ attP_fwd | gacaccaaaaactgccacagg | gtctggtcaaccaccgcggtctcagtggtgtacggtacaaacctgtggcagtttta | A1B3 (attP fwd); A1B4 (attP fwd) |
| IDS2_DF_A4_b_attP_rev | gcactcatttcctccaagctc | taccgtacaccactgagaccgcggtggttgaccagacaaacctcttggaggaaa | A4B7 (attP rev) |
| IDS2_DF_A4_b_ attP_fwd | gcactcatttcctccaagctc | gtctggtcaaccaccgcggtctcagtggtgtacggtacaaacctcttggaggaaa | A4B7 (attP fwd) |
| IDS2_DF_B2_a_attP_rev | gtaggtacaggacagggcag | gtctggtcaaccaccgcggtctcagtggtgtacggtacaaacctccctgtcc | A1B2 (attP rev) |
| IDS2_DF_B2_a_ attP_fwd | gtaggtacaggacagggcag | taccgtacaccactgagaccgcggtggttgaccagacaaacctccctgtcc | A1B2 (attP fwd) |
| IDS2_DF_B3_a_attP_rev | gagataggtaggtacaggaca | tctggtcaaccaccgcggtctcagtggtgtacggtacaaacctcctgtacct | A1B3 (attP rev) |
| IDS2_DF_B3_a_ attP_fwd | gagataggtaggtacaggaca | taccgtacaccactgagaccgcggtggttgaccagacaaacctcctgtacct | A1B3 (attP fwd) |
| IDS2_DF_B4_b_attP_rev | gtgaaaagataggtaggtac | gtctggtcaaccaccgcggtctcagtggtgtacggtacaaacctcctacctatcta | A1B4 (attP rev) |
| IDS2_DF_B4_b_ attP_fwd | gtgaaaagataggtaggtac | taccgtacaccactgagaccgcggtggttgaccagacaaacctcctacctatcta | A1B4 (attP fwd) |
| IDS2_DF_B7_b_attP_rev | gttatggtttactccatcta | gtctggtcaaccaccgcggtctcagtggtgtacggtacaaacctatggagtaaacct | A4B7 (attP rev) |
| IDS2_DF_B7_b_ attP_fwd | gttatggtttactccatcta | taccgtacaccactgagaccgcggtggttgaccagacaaacctatggagtaaacc | A4B7 (attP fwd) |
| IDS_DF_C2_c_attB_rev | gttttggtttaccctatcta | atgatcctgacgacggagaccgccgtcgtcgacaagccatagggtaaacca | C2D2 (attB rev); C2D2 (attB rev) |
| IDS_DF_C2_c_attB_fwd | gttttggtttaccctatcta | ggcttgtcgacgacggcggtctccgtcgtcaggatcatatagggtaaacca | C2D1 (attB fwd); C2D2 (attB fwd) |
| IDS_DF_D1_b_attB_rev | gctgtggaactgcaacacact | ggcttgtcgacgacggcggtctccgtcgtcaggatcatgtgttgcagt | C2D1 (attB rev) |
| IDS_DF_D1_b_attB_fwd | gctgtggaactgcaacacact | atgatcctgacgacggagaccgccgtcgtcgacaagccgtgttgcagt | C2D1 (attB fwd) |
| IDS_DF_D2_c_attB_rev | gtgccacctaacagtgagctg | ggcttgtcgacgacggcggtctccgtcgtcaggatcatctcactgttaggt | C2D2 (attB rev) |
| IDS_DF_D2_c_attB_fwd | gtgccacctaacagtgagctg | atgatcctgacgacggagaccgccgtcgtcgacaagccctcactgttaggt | C2D2 (attB fwd) |

**Figure 4c**

| **pegRNA** | **spacer sequence** | **3' extension** | **Edits made by the specified pegRNA** |
|---|---|---|---|
| IDS2_DF_A4_b_a ttP_rev | gcactcatttcctccaagctc | taccgtacaccactgagaccgcggtggttgaccagacaaacctcttggaggaaa | pegRNA set1 installing attP_rev in IDS2 and attB_fwd in IDS |
| IDS2_DF_B7_b_ attP_rev | gttatggtttactccatcta | gtctggtcaaccaccgcggtctcagtggtgtacggtacaaacctatggagtaaacct | pegRNA set1 installing attP_rev in IDS2 and attB_fwd in IDS |
| IDS_DF_C2_c_ attB_fwd | gttttggtttaccctatcta | ggcttgtcgacgacggcggtctccgtcgtcaggatcatatagggtaaacca | pegRNA set1 installing attP_rev in IDS2 and attB_fwd in IDS |
| IDS_DF_D1_b_ attB_fwd | gctgtggaactgcaacacact | atgatcctgacgacggagaccgccgtcgtcgacaagccgtgttgcagt | pegRNA set1 installing attP_rev in IDS2 and attB_fwd in IDS |
| IDS2_DF_A4_b_ attP_fwd | gcactcatttcctccaagctc | gtctggtcaaccaccgcggtctcagtggtgtacggtacaaacctcttggaggaaa | pegRNA set2 installing attP_fwd in IDS2 and attB_rev in IDS |
| IDS2_DF_B7_b_ attP_fwd | gttatggtttactccatcta | taccgtacaccactgagaccgcggtggttgaccagacaaacctatggagtaaacc | pegRNA set2 installing attP_fwd in IDS2 and attB_rev in IDS |
| IDS_DF_C2_c_attB_rev | gttttggtttaccctatcta | atgatcctgacgacggagaccgccgtcgtcgacaagccatagggtaaacca | pegRNA set2 installing attP_fwd in IDS2 and attB_rev in IDS |
| IDS_DF_D1_b_attB_rev | gctgtggaactgcaacacact | ggcttgtcgacgacggcggtctccgtcgtcaggatcatgtgttgcagt | pegRNA set2 installing attP_fwd in IDS2 and attB_rev in IDS |

**Extended Data Figures**

**ED Figure 1a - Comparison of twinPE- and PE3-mediated *FKBP* insertion at *CCR5* region 1**

| **pegRNA** | **spacer sequence** | **3' extension** | **Edits made by the specified pegRNA** |
|---|---|---|---|
| PE3_CCR5_1_F414a_34 _FKBP108 | ggtacctatcgattgtcagg | tactgtccccttctgggctcactatgctgccgccaaatttctttccatcttcaagcatcccg gtgtagtgcaccacgcaggtctggccgcgcttggggaaggtgcgcccgtctcctggg gagatggtttccacctgcactccgacaatcga | indicated on the x-axis |
| PE3_CCR5_1_F414a_29 _FKBP108 | ggtacctatcgattgtcagg | tccccttctgggctcactatgctgccgccaaatttctttccatcttcaagcatcccggtgta gtgcaccacgcaggtctggccgcgcttggggaaggtgcgcccgtctcctggggaga tggtttccacctgcactccgacaatcga | indicated on the x-axis |
| PE3_CCR5_1_F414a_23 _FKBP108 | ggtacctatcgattgtcagg | tctgggctcactatgctgccgccaaatttctttccatcttcaagcatcccggtgtagtgca ccacgcaggtctggccgcgcttggggaaggtgcgcccgtctcctggggagatggttt ccacctgcactccgacaatcga | indicated on the x-axis |

| | | | |
|---|---|---|---|
| twinPE_CCR5_1_F414a_ FKBP108 | ggtacctatcgattgtcagg | accttccccaagcgcggccagacctgcgtggtgcactacaccgggatgcttgaagat ggaaagaaatttgacaatcga | indicated on the x-axis |
| twinPE_CCR5_1_E325b_ FKBP108 | gctcactatgctgccgcccag | accacgcaggtctggccgcgcttggggaaggtgcgcccgtctcctggggagatggtt tccacctgcactccggcggcagcat | indicated on the x-axis |
| **sgRNA** | **spacer sequence** | | |
| PE3_CCR5_A1_sgRNA | gcatcctgataaactgcaaa | | |
| PE3_CCR5_A2_sgRNA | ggacatctacctgctcaacc | | |
| PE3_CCR5_A4_sgRNA | gcaatgtgtcaactcttgac | | |

**ED Figure 1b - Comparison of twinPE- and PE3-mediated *FKBP* insertion at *CCR5* region 2**

| pegRNA | spacer sequence | 3' extension | Edits made by the specified pegRNA |
|---|---|---|---|
| PE3_CCR5_2_F2_34_ FKBP108 | gtgaaagacagcctggagtc | agattggagaaacccttgaaaagacatcaagcacaaatttctttccatcttcaagcatcc cggtgtagtgcaccacgcaggtctggccgcgcttggggaaggtgcgcccgtctcctg gggagatggtttccacctgcactcctccaggct | indicated on the x-axis |
| PE3_CCR5_2_F2_28_ FKBP108 | gtgaaagacagcctggagtc | gagaaacccttgaaaagacatcaagcacaaatttctttccatcttcaagcatcccggtgt agtgcaccacgcaggtctggccgcgcttggggaaggtgcgcccgtctcctggggag atggtttccacctgcactcctccaggct | indicated on the x-axis |
| PE3_CCR5_2_F2_23_FK BP108 | gtgaaagacagcctggagtc | acccttgaaaagacatcaagcacaaatttctttccatcttcaagcatcccggtgtagtgca ccacgcaggtctggccgcgcttggggaaggtgcgcccgtctcctggggagatggttt ccacctgcactcctccaggct | indicated on the x-axis |
| twinPE_CCR5_2_F2a_ FKBP108 | gtgaaagacagcctggagtc | accttccccaagcgcggccagacctgcgtggtgcactacaccgggatgcttgaagat ggaaagaaattttccaggct | indicated on the x-axis |
| twinPE_CCR5_2_E2c_ FKBP108nt | gaaaagacatcaagcacaga | accacgcaggtctggccgcgcttggggaaggtgcgcccgtctcctggggagatggtt tccacctgcactccgtgcttgatgtctt | indicated on the x-axis |
| **sgRNA** | **spacer sequence** | | |
| PE3_CCR5_C1_sgRNA | gatgcagagtcagcagaact | | |
| PE3_CCR5_C1.5_ sgRNA | ggaagtgagggtcagagagg | | |
| PE3_CCR5_C4_sgRNA | gatggattggtgtaaaagga | | |

**ED Figure 1c - Long twinPE insertions at *CCR5***

| pegRNA | spacer sequence | 3' extension | Edits made by the specified pegRNA |
|---|---|---|---|
| CCR5_A7_attP_spacer27 | gctgtgtttgcgtctctccc | ttcgttatacgcccattcttcgcgaaaggtttgtaccgtacaccactgagaccgcggtgg ttgaccagacaaaccagagacgcaaa | attB-spacer-attP |
| CCR5_B8_spacer_attB | gtatggaaaatgagagctgc | tttcgcgaagaatgggcgtataacgaaggcttgtcgacgacggcggtctccgtcgtca ggatcatgctctcattttc | attB-spacer-attP, attB-spacer-attB |
| CCR5_A7_attB_spacer27 | gctgtgtttgcgtctctccc | ttcgttatacgcccattcttcgcgaaaatgatcctgacgacggagaccgccgtcgtcga caagccagagacgcaaa | attB-spacer-attB |

**ED Figure 2 - *PAH* exon recoding via twinPE**

| pegRNA | spacer sequence | 3' extension | Edits made by the specified pegRNA |
|---|---|---|---|
| PAH_2.1.1 | gctcaaataagcgcaatactt | tgatcttcagcctgaaggaagaggtgggcgccctggcgaaggtattgcgctt | indicated on the x-axis |
| PAH_2.1.2 | gctcaaataagcgcaatactt | tgatcttcagcctgaaggaagaggtgggcgccctggcgaaggtattgcgcttatt | indicated on the x-axis |
| PAH_2.1.3 | gctcaaataagcgcaatactt | tgatcttcagcctgaaggaagaggtgggcgccctggcgaaggtattgcgcttatttg | indicated on the x-axis |
| PAH_2.2.1 | gaagacaactgcaatcaaaa | tcttccttcaggctgaagatcagagagatagcgccgttctgattgcag | indicated on the x-axis |
| PAH_2.2.2 | gaagacaactgcaatcaaaa | tcttccttcaggctgaagatcagagagatagcgccgttctgattgcagttg | indicated on the x-axis |
| PAH_2.2.3 | gaagacaactgcaatcaaaa | tcttccttcaggctgaagatcagagagatagcgccgttctgattgcagttgtc | indicated on the x-axis |
| PAH_4.1.1 | gttctctgtgtttcagtgccc | aagatttgattagcgaacctatcaagttcctgaattgtccgtggaaaccacggcactgaa ac | indicated on the x-axis |
| PAH_4.1.2 | gttctctgtgtttcagtgccc | aagatttgattagcgaacctatcaagttcctgaattgtccgtggaaaccacggcactgaa acac | indicated on the x-axis |
| PAH_4.1.3 | gttctctgtgtttcagtgccc | aagatttgattagcgaacctatcaagttcctgaattgtccgtggaaaccacggcactgaa acacag | indicated on the x-axis |
| PAH_4.1.4 | gttctctgtgtttcagtgccc | gacaagatttgattagcgaacctatcaagttcctgaattgtccgtggaaaccacggcact gaaac | indicated on the x-axis |
| PAH_4.1.5 | gttctctgtgtttcagtgccc | gacaagatttgattagcgaacctatcaagttcctgaattgtccgtggaaaccacggcact gaaacac | indicated on the x-axis |
| PAH_4.1.6 | gttctctgtgtttcagtgccc | gacaagatttgattagcgaacctatcaagttcctgaattgtccgtggaaaccacggcact gaaacacag | indicated on the x-axis |
| PAH_4.2.1 | gcccaagaaccattcaagagc | ggctccgtaggacaagatttgattagcgaacctatcaagttcttgaatgg | indicated on the x-axis |
| PAH_4.2.2 | gcccaagaaccattcaagagc | ggctccgtaggacaagatttgattagcgaacctatcaagttcttgaatggttc | indicated on the x-axis |
| PAH_4.2.3 | gcccaagaaccattcaagagc | ggctccgtaggacaagatttgattagcgaacctatcaagttcttgaatggttcttg | indicated on the x-axis |
| PAH_4.2.4 | gcccaagaaccattcaagagc | gttcggctccgtaggacaagatttgattagcgaacctatcaagttcttgaatgg | indicated on the x-axis |
| PAH_4.2.5 | gcccaagaaccattcaagagc | gttcggctccgtaggacaagatttgattagcgaacctatcaagttcttgaatggttc | indicated on the x-axis |
| PAH_4.2.6 | gcccaagaaccattcaagagc | gttcggctccgtaggacaagatttgattagcgaacctatcaagttcttgaatggttcttg | indicated on the x-axis |
| PAH_4.2.7 | gcccaagaaccattcaagagc | gatgatcagcgtcaagttcggctccgtaggacaagatttgattagcgaacctatcaagtt cttgaatgg | indicated on the x-axis |
| PAH_4.2.8 | gcccaagaaccattcaagagc | gatgatcagcgtcaagttcggctccgtaggacaagatttgattagcgaacctatcaagtt cttgaatggttc | indicated on the x-axis |
| PAH_4.2.9 | gcccaagaaccattcaagagc | gatgatcagcgtcaagttcggctccgtaggacaagatttgattagcgaacctatcaagtt cttgaatggttcttg | indicated on the x-axis |
| PAH_4.3.1 | gcgggccatggactcacaggg | ataggttcgctaatcaaatcttgtcctacggagccgaacttgacgctgatcatcctgtga gtcc | indicated on the x-axis |
| PAH_4.3.2 | gcgggccatggactcacaggg | ataggttcgctaatcaaatcttgtcctacggagccgaacttgacgctgatcatcctgtga gtccat | indicated on the x-axis |

| | | | |
|---|---|---|---|
| PAH_4.3.3 | gcgggccatggactcacaggg | ataggttcgctaatcaaatcttgtcctacggagccgaacttgacgctgatcatcctgtgagtccatgg | indicated on the x-axis |
| PAH_4.3.4 | gcgggccatggactcacaggg | taatcaaatcttgtcctacggagccgaacttgacgctgatcatcctgtgagtcc | indicated on the x-axis |
| PAH_4.3.5 | gcgggccatggactcacaggg | taatcaaatcttgtcctacggagccgaacttgacgctgatcatcctgtgagtccat | indicated on the x-axis |
| PAH_4.3.6 | gcgggccatggactcacaggg | taatcaaatcttgtcctacggagccgaacttgacgctgatcatcctgtgagtccatgg | indicated on the x-axis |
| PAH_4.4.1 | gctacgggccatggactcaca | ggttcgctaatcaaatcttgtcctacggagccgaacttgacgctgatcatcctgtgagtccatg | indicated on the x-axis |
| PAH_4.4.2 | gctacgggccatggactcaca | ggttcgctaatcaaatcttgtcctacggagccgaacttgacgctgatcatcctgtgagtccatggc | indicated on the x-axis |
| PAH_4.4.3 | gctacgggccatggactcaca | ggttcgctaatcaaatcttgtcctacggagccgaacttgacgctgatcatcctgtgagtccatggcc | indicated on the x-axis |
| PAH_4.4.4 | gctacgggccatggactcaca | aaatcttgtcctacggagccgaacttgacgctgatcatcctgtgagtccatg | indicated on the x-axis |
| PAH_4.4.5 | gctacgggccatggactcaca | aaatcttgtcctacggagccgaacttgacgctgatcatcctgtgagtccatggc | indicated on the x-axis |
| PAH_4.4.6 | gctacgggccatggactcaca | aaatcttgtcctacggagccgaacttgacgctgatcatcctgtgagtccatggcc | indicated on the x-axis |
| PAH_4.5.1 | gaaatctcatcctacgggcca | taatcaaatcttgtcctacggagccgaacttgacgctgatcatcctgtgagtccatggcccgtaggat | indicated on the x-axis |
| PAH_4.5.2 | gaaatctcatcctacgggcca | taatcaaatcttgtcctacggagccgaacttgacgctgatcatcctgtgagtccatggcccgtaggatga | indicated on the x-axis |
| PAH_4.5.3 | gaaatctcatcctacgggcca | taatcaaatcttgtcctacggagccgaacttgacgctgatcatcctgtgagtccatggcccgtaggatgag | indicated on the x-axis |
| PAH_5.1.1 | gcaggtgtctcttttctccta | gaattgcttacggcgggcgcgataaacggggtccttgaagccctaggagaaaagag | indicated on the x-axis |
| PAH_5.1.2 | gcaggtgtctcttttctccta | gaattgcttacggcgggcgcgataaacggggtccttgaagccctaggagaaaagagac | indicated on the x-axis |
| PAH_5.1.3 | gcaggtgtctcttttctccta | gaattgcttacggcgggcgcgataaacggggtccttgaagccctaggagaaaagagacac | indicated on the x-axis |
| PAH_5.2.1 | gaagcaaggcagacttactgg | atcgcgcccgccgtaagcaattcgccgatattgcatataattatcgccagtaagtctg | indicated on the x-axis |
| PAH_5.2.2 | gaagcaaggcagacttactgg | atcgcgcccgccgtaagcaattcgccgatattgcatataattatcgccagtaagtctgc | indicated on the x-axis |
| PAH_5.2.3 | gaagcaaggcagacttactgg | atcgcgcccgccgtaagcaattcgccgatattgcatataattatcgccagtaagtctgcc | indicated on the x-axis |
| PAH_7.1.1 | gcttttcatcccagcttgcac | tactgctcaagagcccggcaacggggcggagtcgaaagccagtacaagctggg | indicated on the x-axis |
| PAH_7.1.2 | gcttttcatcccagcttgcac | tactgctcaagagcccggcaacggggcggagtcgaaagccagtacaagctgggat | indicated on the x-axis |
| PAH_7.1.3 | gcttttcatcccagcttgcac | tactgctcaagagcccggcaacggggcggagtcgaaagccagtacaagctgggatga | indicated on the x-axis |
| PAH_7.1.4 | gcttttcatcccagcttgcac | agaaagtctctactgctcaagagcccggcaacggggcggagtcgaaagccagtacaagctggg | indicated on the x-axis |
| PAH_7.1.5 | gcttttcatcccagcttgcac | agaaagtctctactgctcaagagcccggcaacggggcggagtcgaaagccagtacaagctgggat | indicated on the x-axis |
| PAH_7.1.6 | gcttttcatcccagcttgcac | agaaagtctctactgctcaagagcccggcaacggggcggagtcgaaagccagtacaagctgggatga | indicated on the x-axis |
| PAH_7.2.1 | gtggtttccgcctccgacctg | agaaagtctctactgctcaagagcccggcaacgggtcggagg | indicated on the x-axis |
| PAH_7.2.2 | gtggtttccgcctccgacctg | agaaagtctctactgctcaagagcccggcaacgggtcggaggcg | indicated on the x-axis |
| PAH_7.2.3 | gtggtttccgcctccgacctg | agaaagtctctactgctcaagagcccggcaacgggtcggaggcgga | indicated on the x-axis |
| PAH_7.2.4 | gtggtttccgcctccgacctg | gagaccccccagaaagtctctactgctcaagagcccggcaacgggtcggagg | indicated on the x-axis |
| PAH_7.2.5 | gtggtttccgcctccgacctg | gagaccccccagaaagtctctactgctcaagagcccggcaacgggtcggaggcg | indicated on the x-axis |
| PAH_7.2.6 | gtggtttccgcctccgacctg | gagaccccccagaaagtctctactgctcaagagcccggcaacgggtcggaggcgga | indicated on the x-axis |
| PAH_7.2.7 | gtggtttccgcctccgacctg | aacacgcggaatgcgagaccccccagaaagtctctactgctcaagagcccggcaacgggtcggagg | indicated on the x-axis |
| PAH_7.2.8 | gtggtttccgcctccgacctg | aacacgcggaatgcgagaccccccagaaagtctctactgctcaagagcccggcaacgggtcggaggcg | indicated on the x-axis |
| PAH_7.2.9 | gtggtttccgcctccgacctg | aacacgcggaatgcgagaccccccagaaagtctctactgctcaagagcccggcaacgggtcggaggcgga | indicated on the x-axis |
| PAH_7.3.1 | ggctggcctgctttcctctc | gcggaatgcgagaccccccagaaagtctctacaggaaagc | indicated on the x-axis |
| PAH_7.3.2 | ggctggcctgctttcctctc | gcggaatgcgagaccccccagaaagtctctacaggaaagcag | indicated on the x-axis |
| PAH_7.3.3 | ggctggcctgctttcctctc | gcggaatgcgagaccccccagaaagtctctacaggaaagcaggc | indicated on the x-axis |
| PAH_7.3.4 | ggctggcctgctttcctctc | tgggtacaatgaaacacgcggaatgcgagaccccccagaaagtctctacaggaaagc | indicated on the x-axis |
| PAH_7.3.5 | ggctggcctgctttcctctc | tgggtacaatgaaacacgcggaatgcgagaccccccagaaagtctctacaggaaagcag | indicated on the x-axis |
| PAH_7.3.6 | ggctggcctgctttcctctc | tgggtacaatgaaacacgcggaatgcgagaccccccagaaagtctctacaggaaagcaggc | indicated on the x-axis |
| PAH_7.3.7 | ggctggcctgctttcctctc | taatatactgggtacaatgaaacacgcggaatgcgagaccccccagaaagtctctacaggaaagc | indicated on the x-axis |
| PAH_7.3.8 | ggctggcctgctttcctctc | taatatactgggtacaatgaaacacgcggaatgcgagaccccccagaaagtctctacaggaaagcag | indicated on the x-axis |
| PAH_7.3.9 | ggctggcctgctttcctctc | taatatactgggtacaatgaaacacgcggaatgcgagaccccccagaaagtctctacaggaaagcaggc | indicated on the x-axis |
| PAH_7.5.1 | gagtggaagactcggaaggcc | gttgccgggctcttgagcagtagagactttctggggggtctcgccttccgagt | indicated on the x-axis |
| PAH_7.5.2 | gagtggaagactcggaaggcc | gttgccgggctcttgagcagtagagactttctggggggtctcgccttccgagtct | indicated on the x-axis |
| PAH_7.5.3 | gagtggaagactcggaaggcc | gttgccgggctcttgagcagtagagactttctggggggtctcgccttccgagtcttc | indicated on the x-axis |
| PAH_7.5.4 | gagtggaagactcggaaggcc | tcttgagcagtagagactttctggggggtctcgccttccgagt | indicated on the x-axis |
| PAH_7.5.5 | gagtggaagactcggaaggcc | tcttgagcagtagagactttctggggggtctcgccttccgagtct | indicated on the x-axis |
| PAH_7.5.6 | gagtggaagactcggaaggcc | tcttgagcagtagagactttctggggggtctcgccttccgagtcttc | indicated on the x-axis |
| PAH_7.6.1 | gtctgatgtactgtgtgcag | tcttgagcagtagagactttctggggggtctcgcattccgcgtgtttcattgcacacagta | indicated on the x-axis |
| PAH_7.6.2 | gtctgatgtactgtgtgcag | tcttgagcagtagagactttctggggggtctcgcattccgcgtgtttcattgcacacagtaca | indicated on the x-axis |
| PAH_7.6.3 | gtctgatgtactgtgtgcag | tcttgagcagtagagactttctggggggtctcgcattccgcgtgtttcattgcacacagtacatc | indicated on the x-axis |
| PAH_7.6.4 | gtctgatgtactgtgtgcag | agtagagactttctggggggtctcgcattccgcgtgtttcattgcacacagta | indicated on the x-axis |
| PAH_7.6.5 | gtctgatgtactgtgtgcag | agtagagactttctggggggtctcgcattccgcgtgtttcattgcacacagtaca | indicated on the x-axis |

| | | | |
|---|---|---|---|
| PAH_7.6.6 | gtctgatgtactgtgtgcag | agtagagactttctggggggtctcgcattccgcgtgtttcattgcacacagtacatc | indicated on the x-axis |
| PAH_7.6.7 | gtctgatgtactgtgtgcag | tggggggtctcgcattccgcgtgtttcattgcacacagta | indicated on the x-axis |
| PAH_7.6.8 | gtctgatgtactgtgtgcag | tggggggtctcgcattccgcgtgtttcattgcacacagtaca | indicated on the x-axis |
| PAH_7.6.9 | gtctgatgtactgtgtgcag | tggggggtctcgcattccgcgtgtttcattgcacacagtacatc | indicated on the x-axis |
| PAH_7.7.1 | gttcgggggtatacatgggct | gggtctcgcattccgcgtgtttcattgtacccagtatattaggcatggttcaaaacccatgtatac | indicated on the x-axis |
| PAH_7.7.2 | gttcgggggtatacatgggct | gggtctcgcattccgcgtgtttcattgtacccagtatattaggcatggttcaaaacccatgtataccc | indicated on the x-axis |
| PAH_7.7.3 | gttcgggggtatacatgggct | gggtctcgcattccgcgtgtttcattgtacccagtatattaggcatggttcaaaacccatgtatacccc | indicated on the x-axis |
| PAH_7.7.4 | gttcgggggtatacatgggct | attccgcgtgtttcattgtacccagtatattaggcatggttcaaaacccatgtatac | indicated on the x-axis |
| PAH_7.7.5 | gttcgggggtatacatgggct | attccgcgtgtttcattgtacccagtatattaggcatggttcaaaacccatgtataccc | indicated on the x-axis |
| PAH_7.7.6 | gttcgggggtatacatgggct | gtgtttcattgtacccagtatattaggcatggttcaaaacccatgtatacccc | indicated on the x-axis |
| PAH_7.7.7 | gttcgggggtatacatgggct | gtgtttcattgtacccagtatattaggcatggttcaaaacccatgtatac | indicated on the x-axis |
| PAH_7.7.8 | gttcgggggtatacatgggct | gtgtttcattgtacccagtatattaggcatggttcaaaacccatgtataccc | indicated on the x-axis |
| PAH_7.7.9 | gttcgggggtatacatgggct | attccgcgtgtttcattgtacccagtatattaggcatggttcaaaacccatgtatacccc | indicated on the x-axis |
| PAH_7.8.1 | ggacagtactcacggttcgg | gcgtgtttcattgtacccagtatattaggcatggttcaaaaccgatgtacacaccagaaccgtga | indicated on the x-axis |
| PAH_7.8.2 | ggacagtactcacggttcgg | gcgtgtttcattgtacccagtatattaggcatggttcaaaaccgatgtacacaccagaaccgtgagt | indicated on the x-axis |
| PAH_7.8.3 | ggacagtactcacggttcgg | gcgtgtttcattgtacccagtatattaggcatggttcaaaaccgatgtacacaccagaaccgtgagtac | indicated on the x-axis |
| PAH_7.8.4 | ggacagtactcacggttcgg | attgtacccagtatattaggcatggttcaaaaccgatgtacacaccagaaccgtga | indicated on the x-axis |
| PAH_7.8.5 | ggacagtactcacggttcgg | attgtacccagtatattaggcatggttcaaaaccgatgtacacaccagaaccgtgagt | indicated on the x-axis |
| PAH_7.8.6 | ggacagtactcacggttcgg | attgtacccagtatattaggcatggttcaaaaccgatgtacacaccagaaccgtgagtac | indicated on the x-axis |
| PAH_8.1.1 | gtgacatctgccatgagctgt | gaaggacctatcgctgaacagtggcacgtgaccgagaagctcatggc | indicated on the x-axis |
| PAH_8.1.2 | gtgacatctgccatgagctgt | gaaggacctatcgctgaacagtggcacgtgaccgagaagctcatggcag | indicated on the x-axis |
| PAH_8.1.3 | gtgacatctgccatgagctgt | gaaggacctatcgctgaacagtggcacgtgaccgagaagctcatggcagat | indicated on the x-axis |
| PAH_8.2.1 | gtaaaaaatccattccttacc | actgttcagcgataggtccttcgcacagttctctcaggtaaggaatgg | indicated on the x-axis |
| PAH_8.2.2 | gtaaaaaatccattccttacc | actgttcagcgataggtccttcgcacagttctctcaggtaaggaatgga | indicated on the x-axis |
| PAH_8.2.3 | gtaaaaaatccattccttacc | actgttcagcgataggtccttcgcacagttctctcaggtaaggaatggatt | indicated on the x-axis |
| PAH_9.1.1 | gttcccccaattacaggaaat | tatatactcgtctggtgctcccaggcttgccagaccgatctcctgtaatt | indicated on the x-axis |
| PAH_9.1.2 | gttcccccaattacaggaaat | tatatactcgtctggtgctcccaggcttgccagaccgatctcctgtaattgg | indicated on the x-axis |
| PAH_9.1.3 | gttcccccaattacaggaaat | tatatactcgtctggtgctcccaggcttgccagaccgatctcctgtaattggg | indicated on the x-axis |
| PAH_9.1.4 | gttcccccaattacaggaaat | ttctctatatactcgtctggtgctcccaggcttgccagaccgatctcctgtaatt | indicated on the x-axis |
| PAH_9.1.5 | gttcccccaattacaggaaat | ttctctatatactcgtctggtgctcccaggcttgccagaccgatctcctgtaattgg | indicated on the x-axis |
| PAH_9.1.6 | gttcccccaattacaggaaat | ttctctatatactcgtctggtgctcccaggcttgccagaccgatctcctgtaattggg | indicated on the x-axis |
| PAH_9.2.1 | gggagagaagggacttactg | tgggagcaccagacgagtatatagagaaacttgctacagtaagtccctt | indicated on the x-axis |
| PAH_9.2.2 | gggagagaagggacttactg | tgggagcaccagacgagtatatagagaaacttgctacagtaagtcccttc | indicated on the x-axis |
| PAH_9.2.3 | gggagagaagggacttactg | tgggagcaccagacgagtatatagagaaacttgctacagtaagtcccttctc | indicated on the x-axis |
| PAH_9.3.1 | gaccatccacccagggagaga | accagacgagtatatagagaaacttgctacagtaagtcctttctctccctgggt | indicated on the x-axis |
| PAH_9.3.2 | gaccatccacccagggagaga | accagacgagtatatagagaaacttgctacagtaagtcctttctctccctgggtg | indicated on the x-axis |
| PAH_9.3.3 | gaccatccacccagggagaga | accagacgagtatatagagaaacttgctacagtaagtcctttctctccctgggtgga | indicated on the x-axis |
| PAH_10.2.1_EvoPreQ1 | gccagatttactggtttactg | gtaggctttaatgctatcaccctgcttacacaatccgaattcgacggtaaaccagtaaatctctctcttgacgcggttctatctagttacgcgttaaaccaactagaaa | indicated on the x-axis |
| PAH_10.2.2_EvoPreQ1 | gccagatttactggtttactg | gtaggctttaatgctatcaccctgcttacacaatccgaattcgacggtaaaccagtaaatctctctctcttgacgcggttctatctagttacgcgttaaaccaactagaaa | indicated on the x-axis |
| PAH_10.2.3_EvoPreQ1 | gccagatttactggtttactg | gtaggctttaatgctatcaccctgcttacacaatccgaattcgacggtaaaccagtaaatctgtctctctcttgacgcggttctatctagttacgcgttaaaccaactagaaa | indicated on the x-axis |
| PAH_10.3.1_EvoPreQ1 | gtaattcaccaaaggatgac | taagcagggtgatagcattaaagcctacggagcaggtttgctctcatcctttggtgtctctctcttgacgcggttctatctagttacgcgttaaaccaactagaaa | indicated on the x-axis |
| PAH_10.3.2_EvoPreQ1 | gtaattcaccaaaggatgac | taagcagggtgatagcattaaagcctacggagcaggtttgctctcatcctttggtgatctctctcttgacgcggttctatctagttacgcgttaaaccaactagaaa | indicated on the x-axis |
| PAH_10.3.3_EvoPreQ1 | gtaattcaccaaaggatgac | taagcagggtgatagcattaaagcctacggagcaggtttgctctcatcctttggtgaatctctctcttgacgcggttctatctagttacgcgttaaaccaactagaaa | indicated on the x-axis |
| PAH_11.1.1_EvoPreQ1 | gaagccaaagcttctccccc | tgtcactgtatagttctgaattgcagtcttttcgagttcgagtgggagaagctctctcttgacgcggttctatctagttacgcgttaaaccaactagaaa | indicated on the x-axis |
| PAH_11.1.2_EvoPreQ1 | gaagccaaagcttctccccc | tgtcactgtatagttctgaattgcagtcttttcgagttcgagtgggagaagctttctctcttgacgcggttctatctagttacgcgttaaaccaactagaaa | indicated on the x-axis |
| PAH_11.1.3_EvoPreQ1 | gaagccaaagcttctccccc | tgtcactgtatagttctgaattgcagtcttttcgagttcgagtgggagaagctttggtctctctcttgacgcggttctatctagttacgcgttaaaccaactagaaa | indicated on the x-axis |
| PAH_11.2.1_EvoPreQ1 | gaaagcttctcccccctggagc | tgtcactgtatagttctgaattgcagtcttttcgagttccagggggtctctctcttgacgcggttctatctagttacgcgttaaaccaactagaaa | indicated on the x-axis |
| PAH_11.2.2_EvoPreQ1 | gaaagcttctcccccctggagc | tgtcactgtatagttctgaattgcagtcttttcgagttccagggggagtctctctcttgacgcggttctatctagttacgcgttaaaccaactagaaa | indicated on the x-axis |
| PAH_11.2.3_EvoPreQ1 | gaaagcttctcccccctggagc | tgtcactgtatagttctgaattgcagtcttttcgagttccagggggagaatctctctcttgacgcggttctatctagttacgcgttaaaccaactagaaa | indicated on the x-axis |
| PAH_11.4.1_EvoPreQ1 | gaactctctgccacgtaatag | tgcaattcagaactatacagtgacagaatttcaaccattgtattacgtggctctctctcttgacgcggttctatctagttacgcgttaaaccaactagaaa | indicated on the x-axis |
| PAH_11.4.2_EvoPreQ1 | gaactctctgccacgtaatag | tgcaattcagaactatacagtgacagaatttcaaccattgtattacgtggcagtctctctcttgacgcggttctatctagttacgcgttaaaccaactagaaa | indicated on the x-axis |
| PAH_11.4.3_EvoPreQ1 | gaactctctgccacgtaatag | tgcaattcagaactatacagtgacagaatttcaaccattgtattacgtggcagagtctctctcttgacgcggttctatctagttacgcgttaaaccaactagaaa | indicated on the x-axis |
| PAH_12.2.1_EvoPreQ1 | gactttgctgccacaatacct | ttcgatgcgctgtgtataaggatcatatctcacgctaaatggtcttggtattgtggctctctctcttgacgcggttctatctagttacgcgttaaaccaactagaaa | indicated on the x-axis |
| PAH_12.2.2_EvoPreQ1 | gactttgctgccacaatacct | ttcgatgcgctgtgtataaggatcatatctcacgctaaatggtcttggtattgtggcagtctctctcttgacgcggttctatctagttacgcgttaaaccaactagaaa | indicated on the x-axis |

| | | | |
|---|---|---|---|
| PAH_12.2.3_EvoPreQ1 | gactttgctgccacaatacct | ttcgatgcgctgtgtataaggatcatatctcacgctaaatggtcttggtattgtggcagctctctctcttgacgcggttctatctagttacgcgttaaaccaactagaaa | indicated on the x-axis |
| PAH_12.3.1_EvoPreQ1 | gctacgacccatacacccaa | gatcttcaattgttgagtgttatcgagcacttcgatgcgctgggtgtatggtctctctcttgacgcggttctatctagttacgcgttaaaccaactagaaa | indicated on the x-axis |
| PAH_12.3.2_EvoPreQ1 | gctacgacccatacacccaa | gatcttcaattgttgagtgttatcgagcacttcgatgcgctgggtgtatgggttctctctcttgacgcggttctatctagttacgcgttaaaccaactagaaa | indicated on the x-axis |
| PAH_12.3.3_EvoPreQ1 | gctacgacccatacacccaa | gatcttcaattgttgagtgttatcgagcacttcgatgcgctgggtgtatgggtctctctctcttgacgcggttctatctagttacgcgttaaaccaactagaaa | indicated on the x-axis |
| PAH_12.4.1_EvoPreQ1 | gcagccaaaatcttaagctgc | tatgatccttatacacagcgcatcgaagtgctcgataacactcaacagcttaagatttctctctcttgacgcggttctatctagttacgcgttaaaccaactagaaa | indicated on the x-axis |
| PAH_12.4.2_EvoPreQ1 | gcagccaaaatcttaagctgc | tatgatccttatacacagcgcatcgaagtgctcgataacactcaacagcttaagatttagtctctctcttgacgcggttctatctagttacgcgttaaaccaactagaaa | indicated on the x-axis |
| PAH_12.4.3_EvoPreQ1 | gcagccaaaatcttaagctgc | tatgatccttatacacagcgcatcgaagtgctcgataacactcaacagcttaagatttaggtctctctcttgacgcggttctatctagttacgcgttaaaccaactagaaa | indicated on the x-axis |
| PAH_12.5.1_EvoPreQ1 | gtgtaaattacttactgttaa | gataacactcaacaattgaagatcctcgcagacagtatcaacagtaagtaatctctctcttgacgcggttctatctagttacgcgttaaaccaactagaaa | indicated on the x-axis |
| PAH_12.5.2_EvoPreQ1 | gtgtaaattacttactgttaa | gataacactcaacaattgaagatcctcgcagacagtatcaacagtaagtaatttctctctcttgacgcggttctatctagttacgcgttaaaccaactagaaa | indicated on the x-axis |
| PAH_12.5.3_EvoPreQ1 | gtgtaaattacttactgttaa | gataacactcaacaattgaagatcctcgcagacagtatcaacagtaagtaatttatctctctcttgacgcggttctatctagttacgcgttaaaccaactagaaa | indicated on the x-axis |

**ED Figure 3 - pegRNA screen at *AAVS1***

| **pegRNA** | **spacer sequence** | **3' extension** | **Edits made by the specified pegRNA** |
|---|---|---|---|
| AAVS1_A1077a | gcagagccaggaacccctgt | taccgtacaccactgagaccgcggtggttgaccagacaaacctggggttcct | indicated on the x-axis |
| AAVS1_A1077b | gcagagccaggaacccctgt | taccgtacaccactgagaccgcggtggttgaccagacaaacctggggttcctgg | indicated on the x-axis |
| AAVS1_A1077c | gcagagccaggaacccctgt | taccgtacaccactgagaccgcggtggttgaccagacaaacctggggttcctggct | indicated on the x-axis |
| AAVS1_A1098a | gggaaggggcaggagagcca | taccgtacaccactgagaccgcggtggttgaccagacaaacctctctcctgc | indicated on the x-axis |
| AAVS1_A1098b | gggaaggggcaggagagcca | taccgtacaccactgagaccgcggtggttgaccagacaaacctctctcctgccc | indicated on the x-axis |
| AAVS1_A1098c | gggaaggggcaggagagcca | taccgtacaccactgagaccgcggtggttgaccagacaaacctctctcctgcccct | indicated on the x-axis |
| AAVS1_A1246a | gaatatgtcccagatagcac | taccgtacaccactgagaccgcggtggttgaccagacaaacctctatctggg | indicated on the x-axis |
| AAVS1_A1246b | gaatatgtcccagatagcac | taccgtacaccactgagaccgcggtggttgaccagacaaacctctatctgggac | indicated on the x-axis |
| AAVS1_A1246c | gaatatgtcccagatagcac | taccgtacaccactgagaccgcggtggttgaccagacaaacctctatctgggacat | indicated on the x-axis |
| AAVS1_A1267a | ggggactctttaaggaaaga | taccgtacaccactgagaccgcggtggttgaccagacaaacctttccttaaa | indicated on the x-axis |
| AAVS1_A1267b | ggggactctttaaggaaaga | taccgtacaccactgagaccgcggtggttgaccagacaaacctttccttaaaga | indicated on the x-axis |
| AAVS1_A1267c | ggggactctttaaggaaaga | taccgtacaccactgagaccgcggtggttgaccagacaaacctttccttaaagagt | indicated on the x-axis |
| AAVS1_A1293a | gagaaagagaaagggagtag | taccgtacaccactgagaccgcggtggttgaccagacaaacctctccctttc | indicated on the x-axis |
| AAVS1_A1293b | gagaaagagaaagggagtag | taccgtacaccactgagaccgcggtggttgaccagacaaacctctccctttctc | indicated on the x-axis |
| AAVS1_A1293c | gagaaagagaaagggagtag | taccgtacaccactgagaccgcggtggttgaccagacaaacctctccctttctctt | indicated on the x-axis |
| AAVS1_A1307a | gagtagaggcggccacgacc | taccgtacaccactgagaccgcggtggttgaccagacaaacctcgtggccgc | indicated on the x-axis |
| AAVS1_A1307b | gagtagaggcggccacgacc | taccgtacaccactgagaccgcggtggttgaccagacaaacctcgtggccgcct | indicated on the x-axis |
| AAVS1_A1307c | gagtagaggcggccacgacc | taccgtacaccactgagaccgcggtggttgaccagacaaacctcgtggccgcctct | indicated on the x-axis |
| AAVS1_A1582a | gatcagtgaaacgcaccaga | taccgtacaccactgagaccgcggtggttgaccagacaaacctggtgcgttt | indicated on the x-axis |
| AAVS1_A1582b | gatcagtgaaacgcaccaga | taccgtacaccactgagaccgcggtggttgaccagacaaacctggtgcgtttca | indicated on the x-axis |
| AAVS1_A1582c | gatcagtgaaacgcaccaga | taccgtacaccactgagaccgcggtggttgaccagacaaacctggtgcgtttcact | indicated on the x-axis |
| AAVS1_A1615a | gcagctcaggttctgggaga | taccgtacaccactgagaccgcggtggttgaccagacaaacctcccagaacc | indicated on the x-axis |
| AAVS1_A1615b | gcagctcaggttctgggaga | taccgtacaccactgagaccgcggtggttgaccagacaaacctcccagaacctg | indicated on the x-axis |
| AAVS1_A1615c | gcagctcaggttctgggaga | taccgtacaccactgagaccgcggtggttgaccagacaaacctcccagaacctgag | indicated on the x-axis |
| AAVS1_A1647a | gtggccactgagaaccgggc | taccgtacaccactgagaccgcggtggttgaccagacaaacctcggttctca | indicated on the x-axis |
| AAVS1_A1647b | gtggccactgagaaccgggc | taccgtacaccactgagaccgcggtggttgaccagacaaacctcggttctcagt | indicated on the x-axis |
| AAVS1_A1647c | gtggccactgagaaccgggc | taccgtacaccactgagaccgcggtggttgaccagacaaacctcggttctcagtgg | indicated on the x-axis |
| AAVS1_A1810a | gaatctgcctaacaggaggt | taccgtacaccactgagaccgcggtggttgaccagacaaaccttcctgttag | indicated on the x-axis |
| AAVS1_A1810b | gaatctgcctaacaggaggt | taccgtacaccactgagaccgcggtggttgaccagacaaaccttcctgttaggc | indicated on the x-axis |
| AAVS1_A1810c | gaatctgcctaacaggaggt | taccgtacaccactgagaccgcggtggttgaccagacaaaccttcctgttaggcag | indicated on the x-axis |
| AAVS1_A1890a | gtcaccaatcctgtccctag | taccgtacaccactgagaccgcggtggttgaccagacaaacctgggacagga | indicated on the x-axis |
| AAVS1_A1890b | gtcaccaatcctgtccctag | taccgtacaccactgagaccgcggtggttgaccagacaaacctgggacaggatt | indicated on the x-axis |
| AAVS1_A1890c | gtcaccaatcctgtccctag | taccgtacaccactgagaccgcggtggttgaccagacaaacctgggacaggattgg | indicated on the x-axis |
| AAVS1_A3786a | gctggcccccaccgcccca | taccgtacaccactgagaccgcggtggttgaccagacaaacctggcggtggg | indicated on the x-axis |
| AAVS1_A3786b | gctggcccccaccgcccca | taccgtacaccactgagaccgcggtggttgaccagacaaacctggcggtggggg | indicated on the x-axis |
| AAVS1_A3786c | gctggcccccaccgcccca | taccgtacaccactgagaccgcggtggttgaccagacaaacctggcggtgggggc | indicated on the x-axis |
| AAVS1_A3835a | gacgtcacggcgctgcccca | taccgtacaccactgagaccgcggtggttgaccagacaaacctggcagcgcc | indicated on the x-axis |
| AAVS1_A3835b | gacgtcacggcgctgcccca | taccgtacaccactgagaccgcggtggttgaccagacaaacctggcagcgccgt | indicated on the x-axis |
| AAVS1_A3835c | gacgtcacggcgctgcccca | taccgtacaccactgagaccgcggtggttgaccagacaaacctggcagcgccgtga | indicated on the x-axis |
| AAVS1_A3856a | ggtgtgctgggcaggtcgcg | taccgtacaccactgagaccgcggtggttgaccagacaaacctgacctgccc | indicated on the x-axis |
| AAVS1_A3856b | ggtgtgctgggcaggtcgcg | taccgtacaccactgagaccgcggtggttgaccagacaaacctgacctgcccag | indicated on the x-axis |
| AAVS1_A3856c | ggtgtgctgggcaggtcgcg | taccgtacaccactgagaccgcggtggttgaccagacaaacctgacctgcccagca | indicated on the x-axis |
| AAVS1_B1154a | gtccttggcaagcccaggag | gtctggtcaaccaccgcggtctcagtggtgtacggtacaaacctctgggcttg | indicated on the x-axis |
| AAVS1_B1154b | gtccttggcaagcccaggag | gtctggtcaaccaccgcggtctcagtggtgtacggtacaaacctctgggcttgcc | indicated on the x-axis |
| AAVS1_B1154c | gtccttggcaagcccaggag | tctggtcaaccaccgcggtctcagtggtgtacggtacaaacctctgggcttgccaa | indicated on the x-axis |
| AAVS1_B1314a | gtgcgtcctaggtgttcacc | gtctggtcaaccaccgcggtctcagtggtgtacggtacaaacctgaacaccta | indicated on the x-axis |
| AAVS1_B1314b | gtgcgtcctaggtgttcacc | gtctggtcaaccaccgcggtctcagtggtgtacggtacaaacctgaacacctagg | indicated on the x-axis |
| AAVS1_B1314c | gtgcgtcctaggtgttcacc | tctggtcaaccaccgcggtctcagtggtgtacggtacaaacctgaacacctaggac | indicated on the x-axis |
| AAVS1_B1376a | gtcctggcagggctgtggtg | gtctggtcaaccaccgcggtctcagtggtgtacggtacaaacctcacagccct | indicated on the x-axis |
| AAVS1_B1376b | gtcctggcagggctgtggtg | gtctggtcaaccaccgcggtctcagtggtgtacggtacaaacctcacagccctgc | indicated on the x-axis |
| AAVS1_B1376c | gtcctggcagggctgtggtg | tctggtcaaccaccgcggtctcagtggtgtacggtacaaacctcacagccctgcca | indicated on the x-axis |
| AAVS1_B1640a | gtgacctgcccggttctcag | gtctggtcaaccaccgcggtctcagtggtgtacggtacaaacctagaaccggg | indicated on the x-axis |
| AAVS1_B1640b | gtgacctgcccggttctcag | gtctggtcaaccaccgcggtctcagtggtgtacggtacaaacctagaaccgggca | indicated on the x-axis |
| AAVS1_B1640c | gtgacctgcccggttctcag | tctggtcaaccaccgcggtctcagtggtgtacggtacaaacctagaaccgggcagg | indicated on the x-axis |

| | | | |
|---|---|---|---|
| AAVS1_B1676a | gagcttggcagggggtggga | gtctggtcaaccaccgcggtctcagtggtgtacggtacaaacctcaccccctg | indicated on the x-axis |
| AAVS1_B1676b | gagcttggcagggggtggga | gtctggtcaaccaccgcggtctcagtggtgtacggtacaaacctcaccccctgcc | indicated on the x-axis |
| AAVS1_B1676c | gagcttggcagggggtggga | tctggtcaaccaccgcggtctcagtggtgtacggtacaaacctcaccccctgccaa | indicated on the x-axis |
| AAVS1_B1701a | gagccagagaggatcctggg | gtctggtcaaccaccgcggtctcagtggtgtacggtacaaacctaggatcctc | indicated on the x-axis |
| AAVS1_B1701b | gagccagagaggatcctggg | gtctggtcaaccaccgcggtctcagtggtgtacggtacaaacctaggatcctctc | indicated on the x-axis |
| AAVS1_B1701c | gagccagagaggatcctggg | tctggtcaaccaccgcggtctcagtggtgtacggtacaaacctaggatcctctctg | indicated on the x-axis |
| AAVS1_B1705a | gatggagccagagaggatcc | gtctggtcaaccaccgcggtctcagtggtgtacggtacaaaccttcctctctg | indicated on the x-axis |
| AAVS1_B1705b | gatggagccagagaggatcc | gtctggtcaaccaccgcggtctcagtggtgtacggtacaaaccttcctctctggc | indicated on the x-axis |
| AAVS1_B1705c | gatggagccagagaggatcc | tctggtcaaccaccgcggtctcagtggtgtacggtacaaaccttcctctctggctc | indicated on the x-axis |
| AAVS1_B1883a | ggggccactagggacaggat | gtctggtcaaccaccgcggtctcagtggtgtacggtacaaacctctgtccctа | indicated on the x-axis |
| AAVS1_B1883b | ggggccactagggacaggat | gtctggtcaaccaccgcggtctcagtggtgtacggtacaaacctctgtccctagt | indicated on the x-axis |
| AAVS1_B1883c | ggggccactagggacaggat | tctggtcaaccaccgcggtctcagtggtgtacggtacaaacctctgtccctagtgg | indicated on the x-axis |
| AAVS1_B1902a | gtcccctccaccccacagtg | gtctggtcaaccaccgcggtctcagtggtgtacggtacaaaccttgtggggtg | indicated on the x-axis |
| AAVS1_B1902b | gtcccctccaccccacagtg | gtctggtcaaccaccgcggtctcagtggtgtacggtacaaaccttgtggggtgga | indicated on the x-axis |
| AAVS1_B1902c | gtcccctccaccccacagtg | tctggtcaaccaccgcggtctcagtggtgtacggtacaaaccttgtggggtggagg | indicated on the x-axis |
| AAVS1_B1962a | gggaccaccttatattccca | gtctggtcaaccaccgcggtctcagtggtgtacggtacaaacctgaatataag | indicated on the x-axis |
| AAVS1_B1962b | gggaccaccttatattccca | gtctggtcaaccaccgcggtctcagtggtgtacggtacaaacctgaatataaggt | indicated on the x-axis |
| AAVS1_B1962c | gggaccaccttatattccca | tctggtcaaccaccgcggtctcagtggtgtacggtacaaacctgaatataaggtgg | indicated on the x-axis |
| AAVS1_B3839a | gacctgcccagcacaccctg | gtctggtcaaccaccgcggtctcagtggtgtacggtacaaacctggtgtgctg | indicated on the x-axis |
| AAVS1_B3839b | gacctgcccagcacaccctg | gtctggtcaaccaccgcggtctcagtggtgtacggtacaaacctggtgtgctggg | indicated on the x-axis |
| AAVS1_B3839c | gacctgcccagcacaccctg | tctggtcaaccaccgcggtctcagtggtgtacggtacaaacctggtgtgctgggca | indicated on the x-axis |
| AAVS1_B3903a | gcgactcctggaagtggcca | gtctggtcaaccaccgcggtctcagtggtgtacggtacaaacctccacttcca | indicated on the x-axis |
| AAVS1_B3903b | gcgactcctggaagtggcca | gtctggtcaaccaccgcggtctcagtggtgtacggtacaaacctccacttccagg | indicated on the x-axis |
| AAVS1_B3903c | gcgactcctggaagtggcca | tctggtcaaccaccgcggtctcagtggtgtacggtacaaacctccacttccaggag | indicated on the x-axis |
| AAVS1_B3930a | ggacttcccagtgtgcatcg | gtctggtcaaccaccgcggtctcagtggtgtacggtacaaaccttgcacactg | indicated on the x-axis |
| AAVS1_B3930b | ggacttcccagtgtgcatcg | gtctggtcaaccaccgcggtctcagtggtgtacggtacaaaccttgcacactggg | indicated on the x-axis |
| AAVS1_B3930c | ggacttcccagtgtgcatcg | tctggtcaaccaccgcggtctcagtggtgtacggtacaaaccttgcacactgggaa | indicated on the x-axis |

**ED Figure 4 - pegRNA screen in *CCR5* region 1**

| **pegRNA** | **spacer sequence** | **3' extension** | **Edits made by the specified pegRNA** |
|---|---|---|---|
| CCR5_A223a | gtcatcctgataaactgcaaa | atgatcctgacgacggagaccgccgtcgtcgacaagccgcagtttatc | indicated on the x-axis |
| CCR5_A223b | gtcatcctgataaactgcaaa | atgatcctgacgacggagaccgccgtcgtcgacaagccgcagtttatcag | indicated on the x-axis |
| CCR5_A223c | gtcatcctgataaactgcaaa | atgatcctgacgacggagaccgccgtcgtcgacaagccgcagtttatcagg | indicated on the x-axis |
| CCR5_A260a | gtgacatctacctgctcaacc | atgatcctgacgacggagaccgccgtcgtcgacaagcctgagcaggt | indicated on the x-axis |
| CCR5_A260b | gtgacatctacctgctcaacc | atgatcctgacgacggagaccgccgtcgtcgacaagcctgagcaggtag | indicated on the x-axis |
| CCR5_A260c | gtgacatctacctgctcaacc | atgatcctgacgacggagaccgccgtcgtcgacaagcctgagcaggtagat | indicated on the x-axis |
| CCR5_A325a | gctcactatgctgccgcccag | atgatcctgacgacggagaccgccgtcgtcgacaagccggcggcagc | indicated on the x-axis |
| CCR5_A325b | gctcactatgctgccgcccag | atgatcctgacgacggagaccgccgtcgtcgacaagccggcggcagcat | indicated on the x-axis |
| CCR5_A325c | gctcactatgctgccgcccag | atgatcctgacgacggagaccgccgtcgtcgacaagccggcggcagcatag | indicated on the x-axis |
| CCR5_A360a | gacaatgtgtcaactcttgac | atgatcctgacgacggagaccgccgtcgtcgacaagccaagagttgac | indicated on the x-axis |
| CCR5_A360b | gacaatgtgtcaactcttgac | atgatcctgacgacggagaccgccgtcgtcgacaagccaagagttgacac | indicated on the x-axis |
| CCR5_A360c | gacaatgtgtcaactcttgac | atgatcctgacgacggagaccgccgtcgtcgacaagccaagagttgacaca | indicated on the x-axis |
| CCR5_A506a | gacaagtgtgatcacttggg | atgatcctgacgacggagaccgccgtcgtcgacaagccaagtgatcac | indicated on the x-axis |
| CCR5_A506b | gacaagtgtgatcacttggg | atgatcctgacgacggagaccgccgtcgtcgacaagccaagtgatcacac | indicated on the x-axis |
| CCR5_A506c | gacaagtgtgatcacttggg | atgatcctgacgacggagaccgccgtcgtcgacaagccaagtgatcacactt | indicated on the x-axis |
| CCR5_A509a | gaagtgtgatcacttgggtgg | atgatcctgacgacggagaccgccgtcgtcgacaagcccccaagtga | indicated on the x-axis |
| CCR5_A509b | gaagtgtgatcacttgggtgg | atgatcctgacgacggagaccgccgtcgtcgacaagcccccaagtgatc | indicated on the x-axis |
| CCR5_A509c | gaagtgtgatcacttgggtgg | atgatcctgacgacggagaccgccgtcgtcgacaagcccccaagtgatcac | indicated on the x-axis |
| CCR5_A531a | gctgtgtttgcgtctctccc | atgatcctgacgacggagaccgccgtcgtcgacaagccagagacgca | indicated on the x-axis |
| CCR5_A531b | gctgtgtttgcgtctctccc | atgatcctgacgacggagaccgccgtcgtcgacaagccagagacgcaaa | indicated on the x-axis |
| CCR5_A531c | gctgtgtttgcgtctctccc | atgatcctgacgacggagaccgccgtcgtcgacaagccagagacgcaaaca | indicated on the x-axis |
| CCR5_B272a | gaaggaaaaacaggtcagaga | ggcttgtcgacgacggcggtctccgtcgtcaggatcatctgacctg | indicated on the x-axis |
| CCR5_B272b | gaaggaaaaacaggtcagaga | ggcttgtcgacgacggcggtctccgtcgtcaggatcatctgacctgt | indicated on the x-axis |
| CCR5_B272c | gaaggaaaaacaggtcagaga | ggcttgtcgacgacggcggtctccgtcgtcaggatcatctgacctgttt | indicated on the x-axis |
| CCR5_B291a | gcccagaaggggacagtaaga | ggcttgtcgacgacggcggtctccgtcgtcaggatcattactgtccc | indicated on the x-axis |
| CCR5_B291b | gcccagaaggggacagtaaga | ggcttgtcgacgacggcggtctccgtcgtcaggatcattactgtcccct | indicated on the x-axis |
| CCR5_B291c | gcccagaaggggacagtaaga | ggcttgtcgacgacggcggtctccgtcgtcaggatcattactgtccccttc | indicated on the x-axis |
| CCR5_B305a | ggcagcatagtgagcccaga | ggcttgtcgacgacggcggtctccgtcgtcaggatcatgggctcact | indicated on the x-axis |
| CCR5_B305b | ggcagcatagtgagcccaga | ggcttgtcgacgacggcggtctccgtcgtcaggatcatgggctcactat | indicated on the x-axis |
| CCR5_B305c | ggcagcatagtgagcccaga | ggcttgtcgacgacggcggtctccgtcgtcaggatcatgggctcactatgc | indicated on the x-axis |
| CCR5_B326a | gatttccaaagtcccactggg | ggcttgtcgacgacggcggtctccgtcgtcaggatcatagtgggact | indicated on the x-axis |
| CCR5_B326b | gatttccaaagtcccactggg | ggcttgtcgacgacggcggtctccgtcgtcaggatcatagtgggactttg | indicated on the x-axis |
| CCR5_B326c | gatttccaaagtcccactggg | ggcttgtcgacgacggcggtctccgtcgtcaggatcatagtgggactttgg | indicated on the x-axis |
| CCR5_B330a | gttgtatttccaaagtcccac | ggcttgtcgacgacggcggtctccgtcgtcaggatcatggactttgg | indicated on the x-axis |
| CCR5_B330b | gttgtatttccaaagtcccac | ggcttgtcgacgacggcggtctccgtcgtcaggatcatggactttggaa | indicated on the x-axis |
| CCR5_B330c | gttgtatttccaaagtcccac | ggcttgtcgacgacggcggtctccgtcgtcaggatcatggactttggaaat | indicated on the x-axis |
| CCR5_B414a | ggtacctatcgattgtcagg | ggcttgtcgacgacggcggtctccgtcgtcaggatcatgacaatcga | indicated on the x-axis |
| CCR5_B414b | ggtacctatcgattgtcagg | ggcttgtcgacgacggcggtctccgtcgtcaggatcatgacaatcgata | indicated on the x-axis |
| CCR5_B414c | ggtacctatcgattgtcagg | ggcttgtcgacgacggcggtctccgtcgtcaggatcatgacaatcgatagg | indicated on the x-axis |
| CCR5_B535a | gatctggtaaagatgattcc | ggcttgtcgacgacggcggtctccgtcgtcaggatcatatcatcttta | indicated on the x-axis |
| CCR5_B535b | gatctggtaaagatgattcc | ggcttgtcgacgacggcggtctccgtcgtcaggatcatatcatctttacc | indicated on the x-axis |
| CCR5_B535c | gatctggtaaagatgattcc | ggcttgtcgacgacggcggtctccgtcgtcaggatcatatcatctttaccag | indicated on the x-axis |
| CCR5_B584a | gtatggaaaatgagagctgc | ggcttgtcgacgacggcggtctccgtcgtcaggatcatgctctcatt | indicated on the x-axis |
| CCR5_B584b | gtatggaaaatgagagctgc | ggcttgtcgacgacggcggtctccgtcgtcaggatcatgctctcattttc | indicated on the x-axis |
| CCR5_B584c | gtatggaaaatgagagctgc | ggcttgtcgacgacggcggtctccgtcgtcaggatcatgctctcattttcca | indicated on the x-axis |
| CCR5_B601a | gcagaattgatactgactgta | ggcttgtcgacgacggcggtctccgtcgtcaggatcatagtcagtat | indicated on the x-axis |
| CCR5_B601b | gcagaattgatactgactgta | ggcttgtcgacgacggcggtctccgtcgtcaggatcatagtcagtatca | indicated on the x-axis |

| | | | |
|---|---|---|---|
| CCR5_B601c | gcagaattgatactgactgta | ggcttgtcgacgacggcggtctccgtcgtcaggatcatagtcagtatcaat | indicated on the x-axis |

**ED Figure 5a - Comparison of twinPE- and PE3-mediated *attB* insertion at *CCR5* region 1**

| **pegRNA** | **spacer sequence** | **3' extension** | **Edits made by the specified pegRNA** |
|---|---|---|---|
| CCR5_A325b_attB | gctcactatgctgccgcccag | atgatcctgacgacggagaccgccgtcgtcgacaagccggcggcagcat | indicated on the x-axis |
| CCR5_B414a_attB | ggtacctatcgattgtcagg | ggcttgtcgacgacggcggtctccgtcgtcaggatcatgacaatcga | indicated on the x-axis |
| CCR5_B414_23 | ggtacctatcgattgtcagg | tctgggctcactatgctgccgccggcttgtcgacgacggcggtctccgtcgtcaggatcatgacaatcga | indicated on the x-axis |
| CCR5_B414_29 | ggtacctatcgattgtcagg | tcccttctgggctcactatgctgccgccggcttgtcgacgacggcggtctccgtcgtcaggatcatgacaatcga | indicated on the x-axis |
| CCR5_B414_34 | ggtacctatcgattgtcagg | tactgtccccttctgggctcactatgctgccgccggcttgtcgacgacggcggtctccgtcgtcaggatcatgacaatcga | indicated on the x-axis |
| **sgRNA** | **spacer sequence** | | |
| PE3_CCR5_A1_sgRNA | gcatcctgataaactgcaaa | | |
| PE3_CCR5_A2_sgRNA | ggacatctacctgctcaacc | | |
| PE3_CCR5_A4_sgRNA | gcaatgtgtcaactcttgac | | |

**ED Figure 5b - Comparison of twinPE- and PE3-mediated *attB* insertion at *CCR5* region 2**

| **pegRNA** | **spacer sequence** | **3' extension** | **Edits made by the specified pegRNA** |
|---|---|---|---|
| CCR5_C2c_attB | gaaaagacatcaagcacaga | atgatcctgacgacggagaccgccgtcgtcgacaagccgtgcttgatgtctt | indicated on the x-axis |
| CCR5_D2a_attB | gtgaaagacagcctggagtc | ggcttgtcgacgacggcggtctccgtcgtcaggatcattccaggct | indicated on the x-axis |
| CCR5_D2_23 | gtgaaagacagcctggagtc | acccttgaaaagacatcaagcacggcttgtcgacgacggcggtctccgtcgtcaggatcattccaggct | indicated on the x-axis |
| CCR5_D2_28 | gtgaaagacagcctggagtc | gagaaacccttgaaaagacatcaagcacggcttgtcgacgacggcggtctccgtcgtcaggatcattccaggct | indicated on the x-axis |
| CCR5_D2_34 | gtgaaagacagcctggagtc | agattggagaaacccttgaaaagacatcaagcacggcttgtcgacgacggcggtctccgtcgtcaggatcattccaggct | indicated on the x-axis |
| **sgRNA** | **spacer sequence** | | |
| PE3_CCR5_C1_ sgRNA | gatgcagagtcagcagaact | | |
| PE3_CCR5_C1.5_ sgRNA | ggaagtgagggtcagagagg | | |
| PE3_CCR5_C4_sgRNA | gatggattggtgtaaaagga | | |

**ED Figure 6b - HTS junction purity**

| **pegRNA** | **spacer sequence** | **3' extension** | **Edits made by the specified pegRNA** |
|---|---|---|---|
| CCR5_A325a | gctcactatgctgccgcccag | atgatcctgacgacggagaccgccgtcgtcgacaagccggcggcagc | 325/414 |
| CCR5_B414b | ggtacctatcgattgtcagg | ggcttgtcgacgacggcggtctccgtcgtcaggatcatgacaatcgata | 325/414 |
| CCR5_A506c | gacaagtgtgatcacttggg | atgatcctgacgacggagaccgccgtcgtcgacaagccaagtgatcacactt | 506/584 |
| CCR5_A509b | gaagtgtgatcacttgggtgg | atgatcctgacgacggagaccgccgtcgtcgacaagcccccaagtgatc | 509/584 |
| CCR5_A531b | gctgtgtttgcgtctctccc | atgatcctgacgacggagaccgccgtcgtcgacaagccagagacgcaaa | 531/584 |
| CCR5_B584b | gtatggaaaatgagagctgc | ggcttgtcgacgacggcggtctccgtcgtcaggatcatgctctcattttc | 506/584, 509/584, 531/584 |
| AAVS1_A1077c | gcagagccaggaacccctgt | taccgtacaccactgagaccgcggtggttgaccagacaaacctggggttcctggct | 1077/1154 |
| AAVS1_B1154c | gtccttggcaagcccaggag | tctggtcaaccaccgcggtctcagtggtgtacggtacaaacctctgggcttgccaa | 1077/1154 |
| AAVS1_A3786c | gctggccccccaccgcccca | taccgtacaccactgagaccgcggtggttgaccagacaaacctggcggtggggggc | 3786/3903, 3786/3930 |
| AAVS1_B3903c | gcgactcctggaagtggcca | tctggtcaaccaccgcggtctcagtggtgtacggtacaaacctccacttccaggag | 3786/3903 |
| AAVS1_B3930c | ggacttcccagtgtgcatcg | tctggtcaaccaccgcggtctcagtggtgtacggtacaaaccttgcacactgggaa | 3786/3930 |

**ED Figure 6c - Multiplex single transfection knock-in**

| **pegRNA** | **spacer sequence** | **3' extension** | **Edits made by the specified pegRNA** |
|---|---|---|---|
| AAVS1_A1077c | gcagagccaggaacccctgt | taccgtacaccactgagaccgcggtggttgaccagacaaacctggggttcctggct | AAVS1 attP, all samples on x-axis |
| AAVS1_B1154c | gtccttggcaagcccaggag | tctggtcaaccaccgcggtctcagtggtgtacggtacaaacctctgggcttgccaa | AAVS1 attP, all samples on x-axis |
| CCR5_A7_attB_20 | gctgtgtttgcgtctctccc | acgacggagaccgccgtcgtcgacaagccagagacgcaaa | CCR5 attB |
| CCR5_B8_attB_20 | gtatggaaaatgagagctgc | acgacggcggtctccgtcgtcaggatcatgctctcattttc | CCR5 attB |
| CCR5_A7_attB_GA_20 | gctgtgtttgcgtctctccc | acgacggagtccgccgtcgtcgacaagccagagacgcaaa | CCR5 attB-GA |
| CCR5_B8_attB_GA_20 | gtatggaaaatgagagctgc | acgacggcggactccgtcgtcaggatcatgctctcattttc | CCR5 attB-GA |
| CCR5_A7_attP_30 | gctgtgtttgcgtctctccc | cgtacaccactgagaccgcggtggttgaccagacaaaccaagagacgcaaa | CCR5 attP |
| CCR5_B8_attP_30 | gtatggaaaatgagagctgc | ggtcaaccaccgcggtctcagtggtgtacggtacaaacccgctctcattttc | CCR5 attP |
| CCR5_A7_attP_GA_30 | gctgtgtttgcgtctctccc | cgtacaccactgagtccgcggtggttgaccagacaaaccaagagacgcaaa | CCR5 attP-GA |
| CCR5_B8_attP_GA_30 | gtatggaaaatgagagctgc | ggtcaaccaccgcggactcagtggtgtacggtacaaacccgctctcattttc | CCR5 attP-GA |

**ED Figure 6d - Overlap reduction**

| **pegRNA** | **spacer sequence** | **3' extension** | **Edits made by the specified pegRNA** |
|---|---|---|---|
| CCR5_A531b | gctgtgtttgcgtctctccc | atgatcctgacgacggagaccgccgtcgtcgacaagccagagacgcaaa | attB 38 |
| CCR5_B584b | gtatggaaaatgagagctgc | ggcttgtcgacgacggcggtctccgtcgtcaggatcatgctctcattttc | attB 38 |
| CCR5_A7_attB_30 | gctgtgtttgcgtctctccc | tcctgacgacggagaccgccgtcgtcgacaagccagagacgcaaa | attB 30 |
| CCR5_B8_attB_30 | gtatggaaaatgagagctgc | tgtcgacgacggcggtctccgtcgtcaggatcatgctctcattttc | attB 30 |
| CCR5_A7_attB_20 | gctgtgtttgcgtctctccc | acgacggagaccgccgtcgtcgacaagccagagacgcaaa | attB 20 |
| CCR5_B8_attB_20 | gtatggaaaatgagagctgc | acgacggcggtctccgtcgtcaggatcatgctctcattttc | attB 20 |

**ED figure 7 - Factor 9 knock-in**

| pegRNA | spacer sequence | 3' extension | Edits made by the specified pegRNA |
|---|---|---|---|
| CCR5_A7_attB_20 | gctgtgtttgcgtctctccc | acgacggagaccgccgtcgtcgacaagccagagacgcaaa | CCR5 attB |
| CCR5_B8_attB_20 | gtatggaaaatgagagctgc | acgacggcggtctccgtcgtcaggatcatgctctcattttc | CCR5 attB |
| CCR5_A277c | gactgaaacttcacagaata | gaaacttcacagaggcttgtcgacgacggcggtctccgtcgtcaggatcat | ALB attB |
| CCR5_B358c | gatttatgagatcaacagcac | ggcttgtcgacgacggcggtctccgtcgtcaggatcatctgttgatctcat | ALB attB |

**ED figure 8 - TwinPE+Bxb1 mediated inversion in the GFP reporter cells**

| pegRNA | spacer sequence | 3' extension | Edits made by the specified pegRNA |
|---|---|---|---|
| AAVS1_A1077b_attB_rev | gcagagccaggaacccctgt | ggcttgtcgacgacggcggtctccgtcgtcaggatcatggggttcctgg | twinPE-mediated insertion of attB_rev in AAVS1 upstream of H2B_EGFP sequences |
| AAVS1_B1154b_attB_fwd | gtccttggcaagcccaggag | atgatcctgacgacggagaccgccgtcgtcgacaagccctgggcttgcc | twinPE-mediated insertion of attB_rev in AAVS1 upstream of H2B_EGFP sequences |
| AAVS1_A3835b | gacgtcacggcgctgcccca | taccgtacaccactgagaccgcggtggttgaccagacaaacctggcagcgccgt | twinPE-mediated insertion of attP_fwd in AAVS1 downstream of H2B_EGFP sequences |
| AAVS1_B3903b | gcgactcctggaagtggcca | gtctggtcaaccaccgcggtctcagtggtgtacggtacaaacctccacttccagg | twinPE-mediated insertion of attP_fwd in AAVS1 downstream of H2B_EGFP sequences |

**ED Figure 9c - *IDS* and *IDS2* pegRNA screen**

| pegRNA | spacer sequence | 3' extension | Edits made by the specified pegRNA |
|---|---|---|---|
| IDS2_A1_a_attP_rev | gacaccaaaaactgccacagg | taccgtacaccactgagaccgcggtggttgaccagacaaacctgtggcagtta | A1B2 (attP rev) |
| IDS2_A1_a_attP_fwd | gacaccaaaaactgccacagg | gtctggtcaaccaccgcggtctcagtggtgtacggtacaaacctgtggcagtta | A1B2 (attP fwd) |
| IDS2_A1_c_attP_rev | gacaccaaaaactgccacagg | taccgtacaccactgagaccgcggtggttgaccagacaaacctgtggcagtttta | A1B3 (attP rev); A1B4 (attP rev) |
| IDS2_A1_c_attP_fwd | gacaccaaaaactgccacagg | gtctggtcaaccaccgcggtctcagtggtgtacggtacaaacctgtggcagtttta | A1B3 (attP fwd); A1B4 (attP fwd) |
| IDS2_A4_b_attP_rev | gcactcatttcctccaagctc | taccgtacaccactgagaccgcggtggttgaccagacaaacctcttggaggaaa | A4B7 (attP rev) |
| IDS2_A4_b_attP_fwd | gcactcatttcctccaagctc | gtctggtcaaccaccgcggtctcagtggtgtacggtacaaacctcttggaggaaa | A4B7 (attP fwd) |
| IDS2_B2_a_attP_rev | gtaggtacaggacagggcag | gtctggtcaaccaccgcggtctcagtggtgtacggtacaaacctccctgtcc | A1B2 (attP rev) |
| IDS2_B2_a_attP_fwd | gtaggtacaggacagggcag | taccgtacaccactgagaccgcggtggttgaccagacaaacctccctgtcc | A1B2 (attP fwd) |
| IDS2_B3_a_attP_rev | gagataggtaggtacaggaca | tctggtcaaccaccgcggtctcagtggtgtacggtacaaacctcctgtacct | A1B3 (attP rev) |
| IDS2_B3_a_attP_fwd | gagataggtaggtacaggaca | taccgtacaccactgagaccgcggtggttgaccagacaaacctcctgtacct | A1B3 (attP fwd) |
| IDS2_B4_b_attP_rev | gtgaaaagataggtaggtac | gtctggtcaaccaccgcggtctcagtggtgtacggtacaaacctcctacctatcta | A1B4 (attP rev) |
| IDS2_B4_b_attP_fwd | gtgaaaagataggtaggtac | taccgtacaccactgagaccgcggtggttgaccagacaaacctcctacctatcta | A1B4 (attP fwd) |
| IDS2_B7_b_attP_rev | gttatggtttactccatcta | gtctggtcaaccaccgcggtctcagtggtgtacggtacaaacctatggagtaaacct | A4B7 (attP rev) |
| IDS2_B7_b_attP_fwd | gttatggtttactccatcta | taccgtacaccactgagaccgcggtggttgaccagacaaacctatggagtaaacc | A4B7 (attP fwd) |
| IDS_C2_c_attB_rev | gttttggtttaccctatcta | atgatcctgacgacggagaccgccgtcgtcgacaagccatagggtaaacca | C2D2 (attB rev); C2D2 (attB rev) |
| IDS_C2_c_attB_fwd | gttttggtttaccctatcta | ggcttgtcgacgacggcggtctccgtcgtcaggatcatatagggtaaacca | C2D1 (attB fwd); C2D2 (attB fwd) |
| IDS_D1_b_attB_rev | gctgtggaactgcaacacact | ggcttgtcgacgacggcggtctccgtcgtcaggatcatgtgttgcagt | C2D1 (attB rev) |
| IDS_D1_b_attB_fwd | gctgtggaactgcaacacact | atgatcctgacgacggagaccgccgtcgtcgacaagccgtgttgcagt | C2D1 (attB fwd) |
| IDS_D2_c_attB_rev | gtgccacctaacagtgagctg | ggcttgtcgacgacggcggtctccgtcgtcaggatcatctcactgttaggt | C2D2 (attB rev) |
| IDS_D2_c_attB_fwd | gtgccacctaacagtgagctg | atgatcctgacgacggagaccgccgtcgtcgacaagccctcactgttaggt | C2D2 (attB fwd) |
| IDS2_A1_a_attP_rev_EvoPreQ1_motif | gacaccaaaaactgccacagg | taccgtacaccactgagaccgcggtggttgaccagacaaacctgtggcagttaaattattccgcggttctatctagttacgcgttaaaccaactagaa | A1B2 (attP rev) |
| IDS2_A1_a_attP_fwd_EvoPreQ1_motif | gacaccaaaaactgccacagg | gtctggtcaaccaccgcggtctcagtggtgtacggtacaaacctgtggcagttaaaattatgcgcggttctatctagttacgcgttaaaccaactagaa | A1B2 (attP fwd) |
| IDS2_A1_c_attP_rev_EvoPreQ1_motif | gacaccaaaaactgccacagg | taccgtacaccactgagaccgcggtggttgaccagacaaacctgtggcagttttatacatacacgcggttctatctagttacgcgttaaaccaactagaa | A1B3 (attP rev); A1B4 (attP rev) |
| IDS2_A1_c_attP_fwd_EvoPreQ1_motif | gacaccaaaaactgccacagg | gtctggtcaaccaccgcggtctcagtggtgtacggtacaaacctgtggcagttttaaaatatttcgcggttctatctagttacgcgttaaaccaactagaa | A1B3 (attP fwd); A1B4 (attP fwd) |
| IDS2_A4_b_attP_rev_EvoPreQ1_motif | gcactcatttcctccaagctc | taccgtacaccactgagaccgcggtggttgaccagacaaacctcttggaggaaacataataacgcggttctatctagttacgcgttaaaccaactagaa | A4B7 (attP rev) |
| IDS2_A4_b_attP_fwd_EvoPreQ1_motif | gcactcatttcctccaagctc | gtctggtcaaccaccgcggtctcagtggtgtacggtacaaacctcttggaggaaaacataatacgcggttctatctagttacgcgttaaaccaactagaa | A4B7 (attP fwd) |
| IDS2_B2_a_attP_rev_EvoPreQ1_motif | gtaggtacaggacagggcag | gtctggtcaaccaccgcggtctcagtggtgtacggtacaaacctccctgtccctcatagtcgcggttctatctagttacgcgttaaaccaactagaa | A1B2 (attP rev) |
| IDS2_B2_a_attP_fwd_EvoPreQ1_motif | gtaggtacaggacagggcag | taccgtacaccactgagaccgcggtggttgaccagacaaacctccctgtccataaattccgcggttctatctagttacgcgttaaaccaactagaa | A1B2 (attP fwd) |
| IDS2_B3_a_attP_rev_EvoPreQ1_motif | gagataggtaggtacaggaca | gtctggtcaaccaccgcggtctcagtggtgtacggtacaaacctcctgtaccttaaagaatcgcggttctatctagttacgcgttaaaccaactagaa | A1B3 (attP rev) |
| IDS2_B3_a_attP_fwd_EvoPreQ1_motif | gagataggtaggtacaggaca | taccgtacaccactgagaccgcggtggttgaccagacaaacctcctgtaccttgaaagaacgcggttctatctagttacgcgttaaaccaactagaa | A1B3 (attP fwd) |
| IDS2_B4_b_attP_rev_EvoPreQ1_motif | gtgaaaagataggtaggtac | gtctggtcaaccaccgcggtctcagtggtgtacggtacaaacctcctacctatctaaataatcccgcggttctatctagttacgcgttaaaccaactagaa | A1B4 (attP rev) |

| | | | |
|---|---|---|---|
| IDS2_B4_b_attP_fwd_EvoPreQ1_motif | gtgaaaagataggtaggtac | taccgtacaccactgagaccgcggtggttgaccagacaaacctcctacctatctaataaatgccgcggttctatctagttacgcgttaaaccaactagaa | A1B4 (attP fwd) |
| IDS2_B7_b_attP_rev_EvoPreQ1_motif | gttatggtttactccatcta | gtctggtcaaccaccgcggtctcagtggtgtacggtacaaacctatggagtaaaccccctttgccgcggttctatctagttacgcgttaaaccaactagaa | A4B7 (attP rev) |
| IDS2_B7_b_attP_fwd_EvoPreQ1_motif | gttatggtttactccatcta | taccgtacaccactgagaccgcggtggttgaccagacaaacctatggagtaaaccccctttctcgcggttctatctagttacgcgttaaaccaactagaa | A4B7 (attP fwd) |
| IDS_C2_c_attB_rev_EvoPreQ1_motif | gttttggtttaccctatcta | atgatcctgacgacggagaccgccgtcgtcgacaagccatagggtaaaccacatttaaccgcggttctatctagttacgcgttaaaccaactagaa | C2D2 (attB rev); C2D2 (attB rev) |
| IDS_C2_c_attB_fwd_EvoPreQ1_motif | gttttggtttaccctatcta | ggcttgtcgacgacggcggtctccgtcgtcaggatcatatagggtaaaccatttaaagacgcggttctatctagttacgcgttaaaccaactagaa | C2D1 (attB fwd); C2D2 (attB fwd) |
| IDS_D1_b_attB_rev_EvoPreQ1_motif | gctgtggaactgcaacacact | ggcttgtcgacgacggcggtctccgtcgtcaggatcatgtgttgcagtagaaatttcgcggttctatctagttacgcgttaaaccaactagaa | C2D1 (attB rev) |
| IDS_D1_b_attB_fwd_EvoPreQ1_motif | gctgtggaactgcaacacact | atgatcctgacgacggagaccgccgtcgtcgacaagccgtgttgcagtaattaatccgcggttctatctagttacgcgttaaaccaactagaa | C2D1 (attB fwd) |
| IDS_D2_c_attB_rev_EvoPreQ1_motif | gtgccacctaacagtgagctg | ggcttgtcgacgacggcggtctccgtcgtcaggatcatctcactgttaggtattaaattcgcggttctatctagttacgcgttaaaccaactagaa | C2D2 (attB rev) |
| IDS_D2_c_attB_fwd_EvoPreQ1_motif | gtgccacctaacagtgagctg | atgatcctgacgacggagaccgccgtcgtcgacaagccctcactgttaggtaataaatccgcggttctatctagttacgcgttaaaccaactagaa | C2D2 (attB fwd) |

**ED Figure 10a - twinPE mediated *attB* insertion in *CCR5* region 2**

| pegRNA | spacer sequence | 3' extension | Edits made by the specified pegRNA |
|---|---|---|---|
| CCR5_ 2_C1c | gatgcagagtcagcagaact | atgatcctgacgacggagaccgccgtcgtcgacaagcctctgctgactctg | indicated on the x-axis |
| CCR5_ 2_D1c | ggtgcttgatgtcttttcaa | ggcttgtcgacgacggcggtctccgtcgtcaggatcataaaagacatcaagc | indicated on the x-axis |
| CCR5_ 2_C2b | gaaaagacatcaagcacaga | atgatcctgacgacggagaccgccgtcgtcgacaagccgtgcttgatgtc | indicated on the x-axis |
| CCR5_ 2_D4c | gacccctcagtatttcagct | ggcttgtcgacgacggcggtctccgtcgtcaggatcattgaaatactgaggg | indicated on the x-axis |
| CCR5_ 2_C2c | gaaaagacatcaagcacaga | atgatcctgacgacggagaccgccgtcgtcgacaagccgtgcttgatgtctt | indicated on the x-axis |
| CCR5_ 2_D2a | gtgaaagacagcctggagtc | ggcttgtcgacgacggcggtctccgtcgtcaggatcattccaggct | indicated on the x-axis |
| CCR5_ 2_C3b | ggtttaggtcaagaagaaga | atgatcctgacgacggagaccgccgtcgtcgacaagcctcttcttgaccta | indicated on the x-axis |
| CCR5_ 2_D2b | gtgaaagacagcctggagtc | ggcttgtcgacgacggcggtctccgtcgtcaggatcattccaggctgt | indicated on the x-axis |
| CCR5_ 2_D4a | gacccctcagtatttcagct | ggcttgtcgacgacggcggtctccgtcgtcaggatcattgaaatactg | indicated on the x-axis |
| CCR5_ 2_C5c | cacagtctcacccagactcc | atgatcctgacgacggagaccgccgtcgtcgacaagccgtctgggtgagac | indicated on the x-axis |
| CCR5_ 2_D3b | gtatttcagctgggatggga | ggcttgtcgacgacggcggtctccgtcgtcaggatcatcatcccagct | indicated on the x-axis |
| CCR5_ 2_D4b | gacccctcagtatttcagct | ggcttgtcgacgacggcggtctccgtcgtcaggatcattgaaatactgag | indicated on the x-axis |
| CCR5_ 2_C6c | gctagatttatgaatacacg | atgatcctgacgacggagaccgccgtcgtcgacaagccgtattcataaatct | indicated on the x-axis |
| CCR5_ 2_D5c | gaacccatgaaatgactact | ggcttgtcgacgacggcggtctccgtcgtcaggatcatagtcatttcatggg | indicated on the x-axis |
| CCR5_ 2_C7c | gcacacatgagatctaggtg | atgatcctgacgacggagaccgccgtcgtcgacaagccctagatctcatgt | indicated on the x-axis |
| CCR5_ 2_D6b | gatgtgctaaatgctgcctg | ggcttgtcgacgacggcggtctccgtcgtcaggatcatgcagcatttag | indicated on the x-axis |

**ED Figure 10b - twinPE in multiple human cells**

| pegRNA | spacer sequence | 3' extension | Edits made by the specified pegRNA |
|---|---|---|---|
| IDS2_A4_b_attP_rev | gcactcatttcctccaagctc | taccgtacaccactgagaccgcggtggttgaccagacaaacctcttggaggaaa | IDS2 (attP rev) |
| IDS2_B7_b_attP_rev | gttatggtttactccatcta | gtctggtcaaccaccgcggtctcagtggtgtacggtacaaacctatggagtaaacct | IDS2 (attP rev) |
| IDS_C2_c_attB_fwd | gttttggtttaccctatcta | ggcttgtcgacgacggcggtctccgtcgtcaggatcatatagggtaaacca | IDS (attB fwd) |
| IDS_D2_c_attB_fwd | gtgccacctaacagtgagctg | atgatcctgacgacggagaccgccgtcgtcgacaagccctcactgttaggt | IDS (attB fwd) |
| MYC_pegRNA_F1_22nt-insert | gaggctattctgcccatttg | aaatgatggtgatgatggtgttatgggcaga | MYC (22-nt) |
| MYC_pegRNA_R1_22nt-insert | gctttaccccgatccagttc | aacaccatcatcaccatcatttctggatcggggt | MYC (22-nt) |
| TIMM44_pegRNA_F1_22nt-insert | gctggccagcctttctccag | ttcagagaagagatggaagacagagaaaggctgg | TIMM44 (22-nt) |
| TIMM44_pegRNA_R1_22nt-insert | gtcctgtcctggggccatcg | tgtcttccatctcttctctgaatggccccag | TIMM44 (22-nt) |
| CCR5_ 2_C5c | cacagtctcacccagactcc | atgatcctgacgacggagaccgccgtcgtcgacaagccgtctgggtgagac | CCR5_a (attB fwd) |
| CCR5_ 2_D3b | gtatttcagctgggatggga | ggcttgtcgacgacggcggtctccgtcgtcaggatcatcatcccagct | CCR5_a (attB fwd) |
| CCR5_ 2_C2c | gaaaagacatcaagcacaga | atgatcctgacgacggagaccgccgtcgtcgacaagccgtgcttgatgtctt | CCR5_b (attB fwd) |
| CCR5_ 2_D2a | gtgaaagacagcctggagtc | ggcttgtcgacgacggcggtctccgtcgtcaggatcattccaggct | CCR5_b (attB fwd) |

**ED figure 10c – Editing activity of Cas9 nickase, PE2-dead RT variant, and PE2**

| pegRNA | spacer sequence | 3' extension | Edits made by the specified pegRNA |
|---|---|---|---|
| PAH_E7.2_55_EvoPreQ1 | gtggtttccgcctccgacctg | acgcggaatgcgagacccccagaaagtctctactgctcaagagcccggcaacgggtcggaggcgtctctctcttgacgcggttctatctagttacgcgttaaaccaactagaaa | Also used in Fig. 2c |
| PAH_E7.6_56_EvoPreQ1 | gtctgatgtactgtgtgcag | gggctcttgagcagtagagactttctggggggtctcgcattccgcgtgtttcattgcacacagtacatctctctcttgacgcggttctatctagttacgcgttaaaccaactagaaa | Also used in Fig. 2c |
| AAVS1_A3835a | gacgtcacggcgctgcccca | taccgtacaccactgagaccgcggtggttgaccagacaaacctggcagcgcc | Also used in ED Fig. 3 |
| AAVS1_B3930a | ggacttcccagtgtgcatcg | gtctggtcaaccaccgcggtctcagtggtgtacggtacaaaccttgcacactg | Also used in ED Fig. 3 |
| CCR5_2_C2_c_attP fwd_pegRNA1 | gaaaagacatcaagcacaga | aggtttgtaccgtacaccactgagaccgcggtggttgaccagacaaacctgtgcttgatgtctt | |
| CCR5_2_D2_a_attP rev_pegRNA2 | gtgaaagacagcctggagtc | aggtttgtctggtcaaccaccgcggtctcagtggtgtacggtacaaaccttccaggct | |
| PE3_CCR5_1_F414a_34_FKBP108 | ggtacctatcgattgtcagg | tactgtccccttctgggctcactatgctgccgccaaatttctttccatcttcaagcatcccggtgtagtgcaccacgcaggtctggccgcgcttggggaaggtgcgcccgtctcctggggagatggtttccacctgcactccgacaatcga | Also used in ED Fig. 1a |

| | | | |
|---|---|---|---|
| PE3_CCR5_1_F414a_29_FKBP108 | ggtacctatcgattgtcagg | tccccttctgggctcactatgctgccgccaaatttctttccatcttcaagcatcccggtgtagtgcaccacgcaggtctggccgcgcttggggaaggtgcgcccgtctcctggggagatggtttccacctgcactccgacaatcga | Also used in ED Fig. 1a |
| IDS2_DF_A4_b_attP_fwd | gcactcatttcctccaagctc | gtctggtcaaccaccgcggtctcagtggtgtacggtacaaacctcttggaggaaa | Also used in Fig. 4b |
| IDS2_DF_B7_b_attP_fwd | gttatggtttactccatcta | taccgtacaccactgagaccgcggtggttgaccagacaaacctatggagtaaacc | Also used in Fig. 4b |

**Supplementary Note 1 - TwinPE PCR bias assessment**

| **pegRNA** | **spacer sequence** | **3' extension** | **Edits made by the specified pegRNA** |
|---|---|---|---|
| CCR5_A223c | gtcatcctgataaactgcaaa | atgatcctgacgacggagaccgccgtcgtcgacaagccgcagtttatcagg | A223c+B272a; A223c+B291b; A223c+B305a; A223c+B326b; A223c+B330b |
| CCR5_B272a | gaaggaaaaacaggtcagaga | ggcttgtcgacgacggcggtctccgtcgtcaggatcatctgacctg | A223c+B272a |
| CCR5_B291b | gcccagaaggggacagtaaga | ggcttgtcgacgacggcggtctccgtcgtcaggatcattactgtcccct | A223c+B291b |
| CCR5_B305a | ggcagcatagtgagcccaga | ggcttgtcgacgacggcggtctccgtcgtcaggatcatgggctcact | A223c+B305a |
| CCR5_B326b | gatttccaaagtcccactggg | ggcttgtcgacgacggcggtctccgtcgtcaggatcatagtgggactttg | A223c+B326b; A260c+B326b |
| CCR5_B330b | gttgtatttccaaagtcccac | ggcttgtcgacgacggcggtctccgtcgtcaggatcatggactttggaa | A223c+B330b |
| CCR5_A260c | gtgacatctacctgctcaacc | atgatcctgacgacggagaccgccgtcgtcgacaagcctgagcaggtagat | A260c+B305c; A260c+B326b; A260c+B330c |
| CCR5_B305c | ggcagcatagtgagcccaga | ggcttgtcgacgacggcggtctccgtcgtcaggatcatgggctcactatgc | A260c+B305c |
| CCR5_B330c | gttgtatttccaaagtcccac | ggcttgtcgacgacggcggtctccgtcgtcaggatcatggactttggaaat | A260+B330c |
| CCR5_A325b | gctcactatgctgccgcccag | atgatcctgacgacggagaccgccgtcgtcgacaagccggcggcagcat | A325b+B414a |
| CCR5_B414a | ggtacctatcgattgtcagg | ggcttgtcgacgacggcggtctccgtcgtcaggatcatgacaatcga | A325b+B414a; A360b+B414a |
| CCR5_A360b | gacaatgtgtcaactcttgac | atgatcctgacgacggagaccgccgtcgtcgacaagccaagagttgacac | A360b+B414a |
| CCR5_A506c | gacaagtgtgatcacttggg | atgatcctgacgacggagaccgccgtcgtcgacaagccaagtgatcacactt | A506c+B584a |
| CCR5_B584a | gtatggaaaatgagagctgc | ggcttgtcgacgacggcggtctccgtcgtcaggatcatgctctcatt | A506c+B584a |
| CCR5_A509a | gaagtgtgatcacttgggtgg | atgatcctgacgacggagaccgccgtcgtcgacaagcccccaagtga | A509a+B535c; A509a+B584a |
| CCR5_B535c | gatctggtaaagatgattcc | ggcttgtcgacgacggcggtctccgtcgtcaggatcatatcatctttaccag | A509_B535c |
| CCR5_A531c | gctgtgtttgcgtctctccc | atgatcctgacgacggagaccgccgtcgtcgacaagccagagacgcaaaca | A531c+B584b |
| CCR5_B584b | gtatggaaaatgagagctgc | ggcttgtcgacgacggcggtctccgtcgtcaggatcatgctctcattttc | A531c+B584b |
| CCR5_ 2_C2b | gaaaagacatcaagcacaga | atgatcctgacgacggagaccgccgtcgtcgacaagccgtgcttgatgtc | C2b+D4c |
| CCR5_ 2_D4c | gacccctcagtatttcagct | ggcttgtcgacgacggcggtctccgtcgtcaggatcattgaaatactgaggg | C2b+D4c |
| CCR5_ 2_C2c | gaaaagacatcaagcacaga | atgatcctgacgacggagaccgccgtcgtcgacaagccgtgcttgatgtctt | C2c+D2a |
| CCR5_ 2_D2a | gtgaaagacagcctggagtc | ggcttgtcgacgacggcggtctccgtcgtcaggatcattccaggct | C2c+D2a |
| CCR5_ 2_C3b | ggtttaggtcaagaagaaga | atgatcctgacgacggagaccgccgtcgtcgacaagcctcttcttgaccta | C3b+D2b; C3b+D4a |
| CCR5_ 2_D2b | gtgaaagacagcctggagtc | ggcttgtcgacgacggcggtctccgtcgtcaggatcattccaggctgt | C3b+D2b |
| CCR5_ 2_D4a | gacccctcagtatttcagct | ggcttgtcgacgacggcggtctccgtcgtcaggatcattgaaatactg | C3b+D4a |
| CCR5_2_C4b | gatggattggtgtaaaagga | atgatcctgacgacggagaccgccgtcgtcgacaagccttttacaccaatc | C4b+D2a |
| CCR5_2_D2a | gtgaaagacagcctggagtc | ggcttgtcgacgacggcggtctccgtcgtcaggatcattccaggct | C4b+D2a |
| CCR5_ 2_C5c | cacagtctcacccagactcc | atgatcctgacgacggagaccgccgtcgtcgacaagccgtctgggtgagac | C5c+D3b; C5c+D4b |
| CCR5_ 2_D3b | gtatttcagctgggatggga | ggcttgtcgacgacggcggtctccgtcgtcaggatcatcatcccagct | C5c+D3b |
| CCR5_ 2_D4b | gacccctcagtatttcagct | ggcttgtcgacgacggcggtctccgtcgtcaggatcattgaaatactgag | C5c+D4b |
| HEK3_DF_A_SA_del77nt | ggcccagactgagcacgtga | tcctctgccatcacgtgctcagtctg | SA (Δ 77nt) |
| HEK3_DF_B_SA_del77nt | gtcaaccagtatcccggtgc | tgatggcagaggaccgggatactgg | SA (Δ 77nt) |
| HEK3_DF_A_SA_del56nt | ggcccagactgagcacgtga | tggaggaagcagggcttcctttcctctgccatcacgtgctcagtctg | SA (Δ 56nt) |
| HEK3_DF_B_SA_del56nt | gtcaaccagtatcccggtgc | tgatggcagaggaaaggaagccctgcttcctccaccgggatactgg | SA (Δ 56nt) |
| HEK3_DF_A_HA_del64nt | ggcccagactgagcacgtga | tgcaggagctgcatcctctgccatcacgtgctcagtctg | HA (Δ 64nt) |
| HEK3_DF_B_HA_del64nt | gtcaaccagtatcccggtgc | tgatggcagaggatgcagctcctgcaccgggatactgg | HA (Δ 64nt) |
| HEK3_DF_A_PD_del90nt | ggcccagactgagcacgtga | gcccagccaaacttgtcaaccagtatcccggcgtgctcagtctg | PD (Δ 90nt) |
| HEK3_DF_B_PD_del90nt | gtcaaccagtatcccggtgc | gggtcaatccttggggcccagactgagcacgccgggatactgg | PD (Δ 90nt) |

**Supplementary Table 2**. Sequences of primers used for mammalian cell genomic DNA amplification and HTS

| | |
|---|---|
| **Primers (All sequences are shown in 5' to 3' orientation)** | |
| **Figure 1** | |
| HEK3_fwd | ACACTCTTTCCCTACACGACGCTCTTCCGATCTNNNNATGTGGGCTGCCTAGAAAGG |
| HEK3_rev | TGGAGTTCAGACGTGTGCTCTTCCGATCTCCCAGCCAAACTTGTCAACC |
| **Figure 2** | |
| PAH_AVA1686 | ACACTCTTTCCCTACACGACGCTCTTCCGATCTNNNNCCATCACCATTGGCTGGGAT |
| PAH_AVA1687 | TGGAGTTCAGACGTGTGCTCTTCCGATCTAGTGGAGGAGAGGCACTGAA |
| PAH_AVA1696 | ACACTCTTTCCCTACACGACGCTCTTCCGATCTNNNNACCTAAAGGTCTCCTAGTGCCT |
| PAH_AVA1697 | TGGAGTTCAGACGTGTGCTCTTCCGATCTCCAGCAATGAACCCAAACCTC |
| HEK3_fwd | ACACTCTTTCCCTACACGACGCTCTTCCGATCTNNNNATGTGGGCTGCCTAGAAAGG |
| HEK3_rev | TGGAGTTCAGACGTGTGCTCTTCCGATCTCCCAGCCAAACTTGTCAACC |
| DMD_UMI_fwd1 | ACACTCTTTCCCTACACGACGCTCTTCCGATCTNNNNNNNNNNNNNNNNTGCTGGCCAGTTTACTAACAAT |
| DMD_UMI_fwd2 | ACACTCTTTCCCTACACGACGCTCTTCCGATCTNNNNNNNNNNNNNNNNCAGAAAGAAGATCTTATCCCATC TTG |
| DMD_rev0 | TGGAGTTCAGACGTGTGCTCTTCCGATCTGGCTACTTTTGTTATTTGCATT |
| **Figure 3** | |
| AAVS1_AVA1713 | TGGAGTTCAGACGTGTGCTCTTCCGATCTCCAGAGCAGGGTCCCGCTTC |
| AAVS1_AVA1717 | ACACTCTTTCCCTACACGACGCTCTTCCGATCTNNNNACGGGGCTCAGTCTGAAGAG |
| AAVS1_AVA1651 | ACACTCTTTCCCTACACGACGCTCTTCCGATCTNNNNGCCAAGGACTCAAACCCAGA |
| AAVS1_AVA1652 | TGGAGTTCAGACGTGTGCTCTTCCGATCTTCCGTGCGTCAGTTTTACCT |
| AAVS1_AVA1653 | ACACTCTTTCCCTACACGACGCTCTTCCGATCTNNNNAACTGCTTCTCCTCTTGGGAA |
| AAVS1_AVA1715 | TGGAGTTCAGACGTGTGCTCTTCCGATCTTCCTTGCCAGAACCTCTAAGGT |
| AAVS1_AVA1655 | ACACTCTTTCCCTACACGACGCTCTTCCGATCTNNNNATCCTCTCTGGCTCCATCGTA |
| AAVS1_AVA1656 | TGGAGTTCAGACGTGTGCTCTTCCGATCTTCCACTTCAGGACAGCATGTTT |
| AAVS1_AVA1707 | ACACTCTTTCCCTACACGACGCTCTTCCGATCTNNNNCGCCGGGAACTGCCGCTGGC |
| AAVS1_AVA1710 | TGGAGTTCAGACGTGTGCTCTTCCGATCTGAGGAGGCCCTCATCTGGCG |
| CCR5_AVA1678 | ACACTCTTTCCCTACACGACGCTCTTCCGATCTNNNNAATCAATGTGAAGCAAATCGCAGC |
| CCR5_AVA1679 | TGGAGTTCAGACGTGTGCTCTTCCGATCTTCGATTGTCAGGAGGATGATGAA |
| CCR5_AVA1680 | ACACTCTTTCCCTACACGACGCTCTTCCGATCTNNNNTCCTTCTTACTGTCCCCTTCTGGGC |
| CCR5_AVA1681 | TGGAGTTCAGACGTGTGCTCTTCCGATCTGCAAACACAGCATGGACGAC |
| CCR5_AVA1682 | ACACTCTTTCCCTACACGACGCTCTTCCGATCTNNNNACAATCGATAGGTACCTGGCTGTC |
| CCR5_AVA1683 | TGGAGTTCAGACGTGTGCTCTTCCGATCTACCAGCCCCAAGATGACTAT |
| ALB_AVA1760 | ACACTCTTTCCCTACACGACGCTCTTCCGATCTNNNNTTGGCATTTATTTCTAAAATGGCATA |
| ALB_AVA1759 | TGGAGTTCAGACGTGTGCTCTTCCGATCTTCTATCAACAGCAACCAAGAAGACAGACT |
| **Figure 4** | |
| IDS_AVA1763 | ACACTCTTTCCCTACACGACGCTCTTCCGATCTNNNNCTGAAAACCTGAGCTTGGAGG |
| IDS_AVA1764 | TGGAGTTCAGACGTGTGCTCTTCCGATCTGTCTACTCCAGCTTAATGGAAGTGG |
| IDS_AVA1765 | ACACTCTTTCCCTACACGACGCTCTTCCGATCTNNNNGAGAAGATGTGGAAATGCCTCAC |
| IDS_AVA1766 | TGGAGTTCAGACGTGTGCTCTTCCGATCTAATCAACATGAAGGGTTGTGTTGT |
| IDS_AVA1769 | ACACTCTTTCCCTACACGACGCTCTTCCGATCTNNNNGTTCCCACACATGCGTTCCTC |
| IDS_AVA1770 | TGGAGTTCAGACGTGTGCTCTTCCGATCTGGCATGAAGGGTTGTTTTTAATTGA |
| IDS_UMI_junc1_fwd | ACACTCTTTCCCTACACGACGCTCTTCCGATCTNNNNNNNNNNNNNNNNCCCTGTCCTGTACCTACCTAT |
| IDS_junc2_fwd | ACACTCTTTCCCTACACGACGCTCTTCCGATCTNNNNTTGACTCATGCCCTACGAGG |
| IDS_universal_rev | TGGAGTTCAGACGTGTGCTCTTCCGATCTCTCAAATTTACCCGTGGCAGC |
| **ED Figure 1 - Long tPE insertions at CCR5** | |
| CCR5_AVA1682 | ACACTCTTTCCCTACACGACGCTCTTCCGATCTNNNNACAATCGATAGGTACCTGGCTGTC |
| CCR5_AVA1683 | TGGAGTTCAGACGTGTGCTCTTCCGATCTACCAGCCCCAAGATGACTAT |
| **ED Figure 2 - pegRNAs screen at PAH** | |
| PAH_AVA1684 | ACACTCTTTCCCTACACGACGCTCTTCCGATCTNNNNTGTCCATGGAGGTTTAACAGGA |
| PAH_AVA1685 | TGGAGTTCAGACGTGTGCTCTTCCGATCTACATGGAAGTTTGCTACGACAT |
| PAH_AVA1686 | ACACTCTTTCCCTACACGACGCTCTTCCGATCTNNNNCCATCACCATTGGCTGGGAT |
| PAH_AVA1687 | TGGAGTTCAGACGTGTGCTCTTCCGATCTAGTGGAGGAGAGGCACTGAA |
| PAH_AVA1689 | TGGAGTTCAGACGTGTGCTCTTCCGATCTGGTAAGAGGAAGGGAGGGGA |
| PAH_AVA1690 | ACACTCTTTCCCTACACGACGCTCTTCCGATCTNNNNGAGAGCCCCCATTCAAAGCA |
| PAH_AVA1691 | TGGAGTTCAGACGTGTGCTCTTCCGATCTGCAGGACTCTTCATGCTGGT |
| PAH_AVA1696 | ACACTCTTTCCCTACACGACGCTCTTCCGATCTNNNNACCTAAAGGTCTCCTAGTGCCT |
| PAH_AVA1697 | TGGAGTTCAGACGTGTGCTCTTCCGATCTCCAGCAATGAACCCAAACCTC |
| PAH_AVA1702 | ACACTCTTTCCCTACACGACGCTCTTCCGATCTNNNNGGCCAAGTACTAGGTTGGTTCT |
| PAH_AVA1703 | TGGAGTTCAGACGTGTGCTCTTCCGATCTTAACCTGGCTTCCAGGGGAGT |
| PAH_exon10_fwd | ACACTCTTTCCCTACACGACGCTCTTCCGATCTNNNNGACACACCCCAAAATAATGC |
| PAH_exon10_rev | TGGAGTTCAGACGTGTGCTCTTCCGATCTTTGAAAGCACAATAATGGTTTT |
| PAH_exon11_fwd | ACACTCTTTCCCTACACGACGCTCTTCCGATCTNNNNCAGGGAATACTGATCCTGAT |
| PAH_exon11_rev | TGGAGTTCAGACGTGTGCTCTTCCGATCTCAACCACCCACAGATGAGT |
| PAH_exon12_fwd | ACACTCTTTCCCTACACGACGCTCTTCCGATCTNNNNGAGGTGTCCGTGTTCCTAA |
| PAH_exon12_rev | TGGAGTTCAGACGTGTGCTCTTCCGATCTCGATGGTAGGGAAAGACAGT |
| **ED Figure 3 - pegRNA screen at AAVS1** | |
| AAVS1_AVA1713 | TGGAGTTCAGACGTGTGCTCTTCCGATCTCCAGAGCAGGGTCCCGCTTC |

| | |
|---|---|
| AAVS1_AVA1717 | ACACTCTTTCCCTACACGACGCTCTTCCGATCTNNNNACGGGGCTCAGTCTGAAGAG |
| AAVS1_AVA1651 | ACACTCTTTCCCTACACGACGCTCTTCCGATCTNNNNGCCAAGGACTCAAACCCAGA |
| AAVS1_AVA1652 | TGGAGTTCAGACGTGTGCTCTTCCGATCTTCCGTGCGTCAGTTTTACCT |
| AAVS1_AVA1653 | ACACTCTTTCCCTACACGACGCTCTTCCGATCTNNNNAACTGCTTCTCCTCTTGGGAA |
| AAVS1_AVA1715 | TGGAGTTCAGACGTGTGCTCTTCCGATCTTCCTTGCCAGAACCTCTAAGGT |
| AAVS1_AVA1655 | ACACTCTTTCCCTACACGACGCTCTTCCGATCTNNNNATCCTCTCTGGCTCCATCGTA |
| AAVS1_AVA1656 | TGGAGTTCAGACGTGTGCTCTTCCGATCTTCCACTTCAGGACAGCATGTTT |
| AAVS1_AVA1707 | ACACTCTTTCCCTACACGACGCTCTTCCGATCTNNNNCGCCGGGAACTGCCGCTGGC |
| AAVS1_AVA1710 | TGGAGTTCAGACGTGTGCTCTTCCGATCTGAGGAGGCCCTCATCTGGCG |

**ED Figure 4 - pegRNA screen at CCR5 region 1**

| | |
|---|---|
| CCR5_AVA1678 | ACACTCTTTCCCTACACGACGCTCTTCCGATCTNNNNAATCAATGTGAAGCAAATCGCAGC |
| CCR5_AVA1679 | TGGAGTTCAGACGTGTGCTCTTCCGATCTTCGATTGTCAGGAGGATGATGAA |
| CCR5_AVA1680 | ACACTCTTTCCCTACACGACGCTCTTCCGATCTNNNNTCCTTCTTACTGTCCCCTTCTGGGC |
| CCR5_AVA1681 | TGGAGTTCAGACGTGTGCTCTTCCGATCTGCAAACACAGCATGGACGAC |
| CCR5_AVA1682 | ACACTCTTTCCCTACACGACGCTCTTCCGATCTNNNNACAATCGATAGGTACCTGGCTGTC |
| CCR5_AVA1683 | TGGAGTTCAGACGTGTGCTCTTCCGATCTACCAGCCCCAAGATGACTAT |

**ED Figure 5 - Comparison of twinPE- and PE3-mediated attB insertion at CCR5 region 1**

| | |
|---|---|
| CCR5_AVA1680 | ACACTCTTTCCCTACACGACGCTCTTCCGATCTNNNNTCCTTCTTACTGTCCCCTTCTGGGC |
| CCR5_AVA1681 | TGGAGTTCAGACGTGTGCTCTTCCGATCTGCAAACACAGCATGGACGAC |
| CCR5-2_fwd2 | ACACTCTTTCCCTACACGACGCTCTTCCGATCTNNNNAGAGGAGTCAGAGAGAATCCC |
| CCR5-2_rev2 | TGGAGTTCAGACGTGTGCTCTTCCGATCTTTCCTAGACCTCATACCTCGT |

**ED Figure 6b - HTS Junction Purity**

| | |
|---|---|
| CCR5_AVA1680 | ACACTCTTTCCCTACACGACGCTCTTCCGATCTNNNNTCCTTCTTACTGTCCCCTTCTGGGC |
| CCR5_AVA1682 | ACACTCTTTCCCTACACGACGCTCTTCCGATCTNNNNACAATCGATAGGTACCTGGCTGTC |
| AAVS1_AVA1717 | ACACTCTTTCCCTACACGACGCTCTTCCGATCTNNNNACGGGGCTCAGTCTGAAGAG |
| AAVS1_AVA1707 | ACACTCTTTCCCTACACGACGCTCTTCCGATCTNNNNCGCCGGGAACTGCCGCTGGC |
| Donor_CJP140 | TGGAGTTCAGACGTGTGCTCTTCCGATCTGAACTTCAGGGTCAGCTTGC |

**ED Figure 6c - HTS Junction Purity (the other donor-genome junction)**

| | |
|---|---|
| CCR5_AVA1681 | TGGAGTTCAGACGTGTGCTCTTCCGATCTGCAAACACAGCATGGACGAC |
| CCR5_AVA1683 | TGGAGTTCAGACGTGTGCTCTTCCGATCTACCAGCCCCAAGATGACTAT |
| AAVS1_AVA1713 | TGGAGTTCAGACGTGTGCTCTTCCGATCTCCAGAGCAGGGTCCCGCTTC |
| AAVS1_AVA1710 | TGGAGTTCAGACGTGTGCTCTTCCGATCTGAGGAGGCCCTCATCTGGCG |
| Donor_other_fwd | ACACTCTTTCCCTACACGACGCTCTTCCGATCTNNNNGGCAAGCTTACATCGAGATCC |

**ED Figure 6e - Overlap reduction**

| | |
|---|---|
| CCR5_AVA1682 | ACACTCTTTCCCTACACGACGCTCTTCCGATCTNNNNACAATCGATAGGTACCTGGCTGTC |
| CCR5_AVA1683 | TGGAGTTCAGACGTGTGCTCTTCCGATCTACCAGCCCCAAGATGACTAT |

**ED Figure 7b**

| | |
|---|---|
| OT1_fwd | ACACTCTTTCCCTACACGACGCTCTTCCGATCTNNNNGGAAATAAGTTATCACAATGGGAAAT |
| OT1_rev | TGGAGTTCAGACGTGTGCTCTTCCGATCTCGCGATTCTTAAAAGGAGAGG |
| OT2_fwd | ACACTCTTTCCCTACACGACGCTCTTCCGATCTNNNNCCATTCATATTTTGAAACAAAAGG |
| OT2_rev | TGGAGTTCAGACGTGTGCTCTTCCGATCTGCATTGCACTCCTACATACAACA |
| OT3_fwd | ACACTCTTTCCCTACACGACGCTCTTCCGATCTNNNNGCTGTGGTTATTCCCAGCTC |
| OT3_rev | TGGAGTTCAGACGTGTGCTCTTCCGATCTGGGAACACTGGACAAAATCC |
| OT4_fwd | ACACTCTTTCCCTACACGACGCTCTTCCGATCTNNNNGGAAAGCTTTGACAAGTGGAA |
| OT4_rev | TGGAGTTCAGACGTGTGCTCTTCCGATCTGCCTACTTGCCCTTCTTCCT |
| OT5_fwd | ACACTCTTTCCCTACACGACGCTCTTCCGATCTNNNNGCATTGCACTCCTACATACAACA |
| OT5_rev | TGGAGTTCAGACGTGTGCTCTTCCGATCTCCATTCATATTTTGAAACAAAAGG |

**ED Figure 7c**

| | |
|---|---|
| Donor_CJP140 (with OT1_fwd or OT2_fwd) | TGGAGTTCAGACGTGTGCTCTTCCGATCTGAACTTCAGGGTCAGCTTGC |
| Donor_fwd_primer (with OT3_rev, or OT4_rev, or OT5_rev) | ACACTCTTTCCCTACACGACGCTCTTCCGATCTNNNNGAACTTCAGGGTCAGCTTGC |

**ED Figure 9**

| | |
|---|---|
| IDS_AVA1763 | ACACTCTTTCCCTACACGACGCTCTTCCGATCTNNNNCTGAAAACCTGAGCTTGGAGG |
| IDS_AVA1764 | TGGAGTTCAGACGTGTGCTCTTCCGATCTGTCTACTCCAGCTTAATGGAAGTGG |
| IDS_AVA1765 | ACACTCTTTCCCTACACGACGCTCTTCCGATCTNNNNGAGAAGATGTGGAAATGCCTCAC |
| IDS_AVA1766 | TGGAGTTCAGACGTGTGCTCTTCCGATCTAATCAACATGAAGGGTTGTGTTGT |
| IDS_AVA1769 | ACACTCTTTCCCTACACGACGCTCTTCCGATCTNNNNGTTCCCACACATGCGTTCCTC |
| IDS_AVA1770 | TGGAGTTCAGACGTGTGCTCTTCCGATCTGGCATGAAGGGTTGTTTTTAATTGA |

**ED Figure 10**

| | |
|---|---|
| CCR5-2_Fwd1 | ACACTCTTTCCCTACACGACGCTCTTCCGATCTNNNNGGTATTCGTGCAGCATATGAG |
| CCR5-2_Rev1 | TGGAGTTCAGACGTGTGCTCTTCCGATCTTATTTCAGCTGGGATGGGAAGG |
| CCR5-2_fwd2 | ACACTCTTTCCCTACACGACGCTCTTCCGATCTNNNNAGAGGAGTCAGAGAGAATCCC |
| CCR5-2_rev2 | TGGAGTTCAGACGTGTGCTCTTCCGATCTTTCCTAGACCTCATACCTCGT |
| CCR5-2_Fwd3 | ACACTCTTTCCCTACACGACGCTCTTCCGATCTNNNNCACTGAATGCTTCTGACTTCATAG |
| CCR5-2_Rev3 | TGGAGTTCAGACGTGTGCTCTTCCGATCTTTTGCTCAATGCTTTGCTCAGT |
| IDS_AVA1765 | ACACTCTTTCCCTACACGACGCTCTTCCGATCTNNNNGAGAAGATGTGGAAATGCCTCAC |
| IDS_AVA1766 | TGGAGTTCAGACGTGTGCTCTTCCGATCTAATCAACATGAAGGGTTGTGTTGT |

| | |
|---|---|
| IDS_AVA1769 | ACACTCTTTCCCTACACGACGCTCTTCCGATCTNNNNGTTCCCACACATGCGTTCCTC |
| IDS_AVA1770 | TGGAGTTCAGACGTGTGCTCTTCCGATCTGGCATGAAGGGTTGTTTTTAATTGA |
| CCR5-2_fwd2 | ACACTCTTTCCCTACACGACGCTCTTCCGATCTNNNNAGAGGAGTCAGAGAGAATCCC |
| CCR5-2_rev2 | TGGAGTTCAGACGTGTGCTCTTCCGATCTTTCCTAGACCTCATACCTCGT |
| MYC_fwd | ACACTCTTTCCCTACACGACGCTCTTCCGATCTNNNNTTGCACTGGAACTTACAACAC |
| MYC_rev | TGGAGTTCAGACGTGTGCTCTTCCGATCTGGCAGAAATCTCGAAAGG |
| TIMM44_fwd | ACACTCTTTCCCTACACGACGCTCTTCCGATCTNNNNAGACCTGTACATTCGGGC |
| TIMM44_rev | TGGAGTTCAGACGTGTGCTCTTCCGATCTAAAAGCCCAGTGCTGCTC |

**Supplementary Table 5 - TwinPE-mediated off-target genome editing**

| | |
|---|---|
| HEK3_OT1_fwd | ACACTCTTTCCCTACACGACGCTCTTCCGATCTNNNNTCCCCTGTTGACCTGGAGAA |
| HEK3_OT1_rev | TGGAGTTCAGACGTGTGCTCTTCCGATCTCACTGTACTTGCCCTGACCA |
| HEK3_OT2_fwd | ACACTCTTTCCCTACACGACGCTCTTCCGATCTNNNNTTGGTGTTGACAGGGAGCAA |
| HEK3_OT2_rev | TGGAGTTCAGACGTGTGCTCTTCCGATCTCTGAGATGTGGGCAGAAGGG |
| HEK3_OT3_fwd | ACACTCTTTCCCTACACGACGCTCTTCCGATCTNNNNTGAGAGGGAACAGAAGGGCT |
| HEK3_OT3_rev | TGGAGTTCAGACGTGTGCTCTTCCGATCTGTCCAAAGGCCCAAGAACCT |
| HEK3_OT4_fwd | ACACTCTTTCCCTACACGACGCTCTTCCGATCTNNNNTCCTAGCACTTTGGAAGGTCG |
| HEK3_OT4_rev | TGGAGTTCAGACGTGTGCTCTTCCGATCTGCTCATCTTAATCTGCTCAGCC |

**Supplementary Note 1 - Analysis of editing quantification bias**

| | |
|---|---|
| UMI_HEK3_fwd | ACACTCTTTCCCTACACGACGCTCTTCCGATCTNNNNNNNNNNNNNNNNATGTGGGCTGCCTAGAAAGG |
| UMI_CCR5-2_fwd | ACACTCTTTCCCTACACGACGCTCTTCCGATCTNNNNNNNNNNNNNNNNAGAGGAGTCAGAGAGAATCCC |
| UMI_AVA1678_fwd | ACACTCTTTCCCTACACGACGCTCTTCCGATCTNNNNNNNNNNNNNNNNAATCAATGTGAAGCAAATCGCAGC |
| UMI_AVA1680_fwd | ACACTCTTTCCCTACACGACGCTCTTCCGATCTNNNNNNNNNNNNNNNNTCCTTCTTACTGTCCCCTTCTGGGC |
| UMI_AVA1682_fwd | ACACTCTTTCCCTACACGACGCTCTTCCGATCTNNNNNNNNNNNNNNNNACAATCGATAGGTACCTGGCTGTC |
| 5p constant primer | ACACTCTTTCCCTACACG |

**Supplementary Table 3**. Sequences of primers and probes used for ddPCR assays

**Fig 3d**

| Assay | fwd | rev | Probe | Samples |
|---|---|---|---|---|
| CCR5_B6 | catctctgacctgtttttcc | tcctcgcccttgctcac | /56-FAM/ACGACGGCG/ZEN/GTCTCAGTGGTG/3IABkFQ/ | 325/414 |
| CCR5_B8_1 | gccaggacggtcaccttg | tcctcgcccttgctcac | /56-FAM/ACGACGGCG/ZEN/GTCTCAGTGGTG/3IABkFQ/ | 507/584, 510/584, 532/584 |
| AAVS1_1077 | ggaacggggctcagtct | tcctcgcccttgctcac | /56-FAM/ACCACCGCG/ZEN/GTCTCCGTCGT/3IABkFQ/ | 1077/1154 |
| AAVS1_3786 | ggcaagcttacatcgagatcc | gaggaggccctcatctggcg | /56-FAM/ACGACGGCG/ZEN/GTCTCAGTGGTG/3IABkFQ/ | 3786/3903, 3786/3930 |
| ACTB | acactgtgcccatctac | aatgtcacgcacgatttc | /5HEX/CGGGACCTG/ZEN/ACTGACTACCTCAT/3IABkFQ/ | all |

**Fig 3e**

| Assay | fwd | rev | Probe | Samples |
|---|---|---|---|---|
| CCR5_B8_2 | gccaggacggtcaccttg | tcctcgcccttgctcac | /56-FAM/CTCAGTGGT/ZEN/GTACGGTACAAACCC/3IABkFQ/ | all |
| ACTB | acactgtgcccatctac | aatgtcacgcacgatttc | /5HEX/CGGGACCTG/ZEN/ACTGACTACCTCAT/3IABkFQ/ | all |

**Fig 3g**

| Assay | fwd | rev | Probe | Samples |
|---|---|---|---|---|
| CCR5_B8_1 | gccaggacggtcaccttg | tcctcgcccttgctcac | /56-FAM/ACGACGGCG/ZEN/GTCTCAGTGGTG/3IABkFQ/ | CCR5 samples |
| ALB | gtgactgtaattttcttttgcg | tcctcgcccttgctcac | /56-FAM/ACGACGGCG/ZEN/GTCTCAGTGGTG/3IABkFQ/ | ALB samples |
| ACTB | acactgtgcccatctac | aatgtcacgcacgatttc | /5HEX/CGGGACCTG/ZEN/ACTGACTACCTCAT/3IABkFQ/ | all |

**ED Fig 6a**

| Assay | fwd | rev | Probe | Samples |
|---|---|---|---|---|
| CCR5_B8_2 | gccaggacggtcaccttg | tcctcgcccttgctcac | /56-FAM/CTCAGTGGT/ZEN/GTACGGTACAAACCC/3IABkFQ/ | all |
| ACTB | acactgtgcccatctac | aatgtcacgcacgatttc | /5HEX/CGGGACCTG/ZEN/ACTGACTACCTCAT/3IABkFQ/ | all |

**ED Fig 6c**

| Assay | fwd | rev | Probe | Samples |
|---|---|---|---|---|
| AAVS1_1077 | ggaacggggctcagtct | tcctcgcccttgctcac | /56-FAM/ACCACCGCG/ZEN/GTCTCCGTCGT/3IABkFQ/ | all |
| CCR5_B8_1 | gccaggacggtcaccttg | tcctcgcccttgctcac | /56-FAM/ACGACGGCG/ZEN/GTCTCAGTGGTG/3IABkFQ/ | attB |
| CCR5_B8_2 | gccaggacggtcaccttg | tcctcgcccttgctcac | /56-FAM/CTCAGTGGT/ZEN/GTACGGTACAAACCC/3IABkFQ/ | attB-GA |
| CCR5_B8_3 | gccaggacggtcaccttg | tcctcgcccttgctcac | /56-FAM/ACCACCGCG/ZEN/GTCTCCGTCGT/3IABkFQ/ | attP |
| CCR5_B8_4 | gccaggacggtcaccttg | tcctcgcccttgctcac | /56-FAM/CTCCGTCGT/ZEN/CAGGATCATCCGT/3IABkFQ/ | attP-GA |
| ACTB | acactgtgcccatctac | aatgtcacgcacgatttc | /5HEX/CGGGACCTG/ZEN/ACTGACTACCTCAT/3IABkFQ/ | all |

**Supplementary Table 4**. Sequence of recoded exonic *PAH* sequences

Nucleotides labeled in red indicate positions where silent mutations were introduced.

| Spacer 1 | Spacer 2 | Recoded allele product |
|---|---|---|
| 2.1 | 2.2 | GAAGACAACTGCAATCAGAACGGCGCTATCTCTCTGATCTTCAGCCTGAAGGAAGAGGTGGGCGCCCTGGCGAAGGTATTGCGCTTATTTGAG |
| 4.1 | 4.3 | TTCTCTGTGTTTCAGTGCCGTGGTTTCCACGGACAATTCAGGAACTTGATAGGTTCGCTAATCAAATCTTGTCCTACGGAGCCGAACTTGACGCTGATCATCCTGTGAGTCCATGGCCCG |
| 4.1 | 4.4 | TTCTCTGTGTTTCAGTGCCGTGGTTTCCACGGACAATTCAGGAACTTGATAGGTTCGCTAATCAAATCTTGTCCTACGGAGCCGAACTTGACGCTGATCATCCTGTGAGTCCATGGCCCGTAG |
| 4.2 | 4.3 | CCCAAGAACCATTCAAGAACTTGATAGGTTCGCTAATCAAATCTTGTCCTACGGAGCCGAACTTGACGCTGATCATCCTGTGAGTCCATGGCCCG |
| 4.2 | 4.4 | CCCAAGAACCATTCAAGAACTTGATAGGTTCGCTAATCAAATCTTGTCCTACGGAGCCGAACTTGACGCTGATCATCCTGTGAGTCCATGGCCCGTAG |
| 4.2 | 4.5 | CCCAAGAACCATTCAAGAACTTGATAGGTTCGCTAATCAAATCTTGTCCTACGGAGCCGAACTTGACGCTGATCATCCTGTGAGTCCATGGCCCGTAGGATGAGATTT |
| 5.1 | 5.2 | CAGGTGTCTCTTTTCTCCTAGGGCTTCAAGGACCCCGTTTATCGCGCCCGCCGTAAGCAATTCGCCGATATTGCATATAATTATCGCCAGTAAGTCTGCCTTGCTT |
| 7.1 | 7.5 | CTTTTCATCCCAGCTTGTACTGGCTTTCGACTCCGCCCCGTTGCCGGGCTCTTGAGCAGTAGAGACTTTCTGGGGGGTCTCGCCTTCCGAGTCTTCCACT |
| 7.1 | 7.6 | CTTTTCATCCCAGCTTGTACTGGCTTTCGACTCCGCCCCGTTGCCGGGCTCTTGAGCAGTAGAGACTTTCTGGGGGGTCTCGCATTCCGCGTGTTTCATTGCACACAGTACATCAGAC |
| 7.2 | 7.5 | TGGTTTCCGCCTCCGACCCGTTGCCGGGCTCTTGAGCAGTAGAGACTTTCTGGGGGGTCTCGCCTTCCGAGTCTTCCACT |
| 7.2 | 7.6 | TGGTTTCCGCCTCCGACCCGTTGCCGGGCTCTTGAGCAGTAGAGACTTTCTGGGGGGTCTCGCATTCCGCGTGTTTCATTGCACACAGTACATCAGAC |
| 7.2 | 7.7 | TGGTTTCCGCCTCCGACCCGTTGCCGGGCTCTTGAGCAGTAGAGACTTTCTGGGGGGTCTCGCATTCCGCGTGTTTCATTGTACCCAGTATATTAGGCATGGTTCAAAACCGATGTACACACCAGAA |
| 7.3 | 7.6 | GGCTGGCCTGCTTTCCTGTAGAGACTTTCTGGGGGGTCTCGCATTCCGCGTGTTTCATTGCACACAGTACATCAGAC |
| 7.3 | 7.7 | GGCTGGCCTGCTTTCCTGTAGAGACTTTCTGGGGGGTCTCGCATTCCGCGTGTTTCATTGTACCCAGTATATTAGGCATGGTTCAAAACCGATGTACACACCAGAA |
| 7.3 | 7.8 | GGCTGGCCTGCTTTCCTGTAGAGACTTTCTGGGGGGTCTCGCATTCCGCGTGTTTCATTGTACCCAGTATATTAGGCATGGTTCAAAACCGATGTACACACCAGAACCGTGAGTACTGTCC |
| 9.1 | 9.2 | TTCCCCCAATTACAGGAGATCGGTCTGGCAAGCCTGGGAGCACCAGACGAGTATATAGAGAAACTTGCTACAGTAAGTCCCTTCTCTCCC |
| 9.1 | 9.3 | TTCCCCCAATTACAGGAGATCGGTCTGGCAAGCCTGGGAGCACCAGACGAGTATATAGAGAAACTTGCTACAGTAAGTCCCTTCTCTCCCTGGGTGGATGGT |
| 10.2 | 10.3 | CCAGATTTACTGGTTTACCGTCGAATTCGGATTGTGTAAGCAGGGTGATAGCATTAAAGCCTACGGAGCAGGTTTGCTCTCATCCTTTGGTGAATTAC |
| 11.1 | 11.4 | GAAGCCAAAGCTTCTCCCACTCGAACTCGAAAAGACTGCAATTCAGAACTATACAGTGACAGAATTTCAACCATTGTATTACGTGGCAGAGAGTT |
| 11.2 | 11.4 | AAAGCTTCTCCCCCTGGAACTCGAAAAGACTGCAATTCAGAACTATACAGTGACAGAATTTCAACCATTGTATTACGTGGCAGAGAGTT |
| 12.2 | 12.4 | ACTTTGCTGCCACAATACCAAGACCATTTAGCGTGAGATATGATCCTTATACACAGCGCATCGAAGTGCTCGATAACACTCAACAGCTTAAGATTTTGGCTG |
| 12.3 | 12.5 | GCTACGACCCATACACCCAGCGCATCGAAGTGCTCGATAACACTCAACAATTGAAGATCCTCGCAGACAGTATCAACAGTAAGTAATTTACAC |

**Supplementary Table 5**. TwinPE-mediated off-target genome editing.

**twinPE-mediated % editing at *HEK3* on-target site and off-target sites**

| ***HEK3*** | | **standerd pegRNAs** | | | | **enhanced pegRNAs** | | | |
|---|---|---|---|---|---|---|---|---|---|
| | Untreated | SA (Δ77 nt) | SA (Δ56 nt) | HA (Δ64 nt) | PD (Δ90 nt) | SA (Δ77 nt) | SA (Δ56 nt) | HA (Δ64 nt) | PD (Δ90 nt) |
| on-target site | <0.1 | 14.8 | 18.8 | 11.7 | 40.4 | 29.6 | 28.5 | 29.5 | 59.5 |
| off-target site 1 | <0.1 | <0.1 | <0.1 | <0.1 | <0.1 | <0.1 | <0.1 | <0.1 | <0.1 |
| off-target site 2 | <0.1 | <0.1 | <0.1 | <0.1 | <0.1 | <0.1 | <0.1 | <0.1 | <0.1 |
| off-target site 3 | <0.1 | <0.1 | <0.1 | <0.1 | <0.1 | <0.1 | <0.1 | <0.1 | <0.1 |
| off-target site 4 | <0.1 | <0.1 | <0.1 | <0.1 | <0.1 | <0.1 | <0.1 | <0.1 | <0.1 |

**Supplementary Note 1**. Analysis of editing quantification bias

To analyze quantification bias, we applied unique molecular identifiers (UMIs) to index individual allele copies using linear amplification followed by bead-based purification, PCR amplification, and Illumina MiSeq amplicon sequencing (see method section for details) similar to the method used by Choi, et al[1] and Bolukbasi, et al[2]. Reads containing identical UMIs are first aligned and then collapsed (deduplicated) to a single consensus sequence. This workflow allows for quantification of allele frequencies based on UMI counts instead of total read counts, which may be subject to PCR amplification bias based on sequence composition or amplicon size differences between different allele products.

To ensure the UMI-sequencing protocol can faithfully quantify the editing efficiency, we mixed chemically synthesized gene fragments (IDT) replicating either an edited or wild-type allele for the given edits (insertion of *attB* at *CCR5* or 77-nt deletion at *HEK3*) to simulate 0%, 20%, 40%, 60%, 80%, and 100% editing efficiencies before diluting the mixed standards to 2000 genetic copies/µL, which is the estimated allele copy number amplified from our cell editing experiments. Diluted standards were then subjected to UMI barcoding, MiSeq analysis, and deduplication. We plotted the percentage of observed edited alleles after sequencing and UMI deduplication against the expected edited allele percentage based on the input allele fraction.

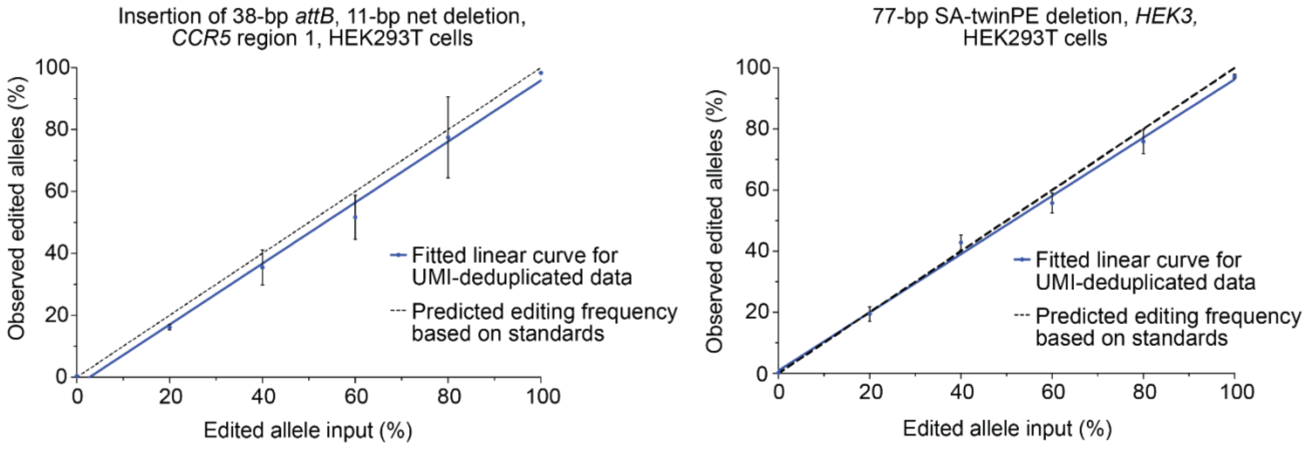

Next, we prepared libraries of twinPE-edited samples from *CCR5* region 1 (15 samples), *CCR5* region 2 (7 samples), *HEK3* (4 samples), and *IDS* (1 sample). By comparing

the editing efficiency calculated from total reads and the editing efficiency calculated from reads deduplicated based on UMIs, we estimated the bias in editing efficiency quantification using the linear amplification with UMIs as the “true” editing efficiency. When plotting quantification bias vs. absolute size difference between the starting and edited alleles (net deletion size), we observed a positive correlation between bias and twinPE-induced deletion size. Although bias in this workflow could still exist in the linear amplification stage due to differences in sequence composition at the target DNA locus, this bias does not propagate over the course of repeated PCR cycles and should therefore more accurately represent the true editing efficiency in twinPE editing experiments analyzed by amplicon sequencing.

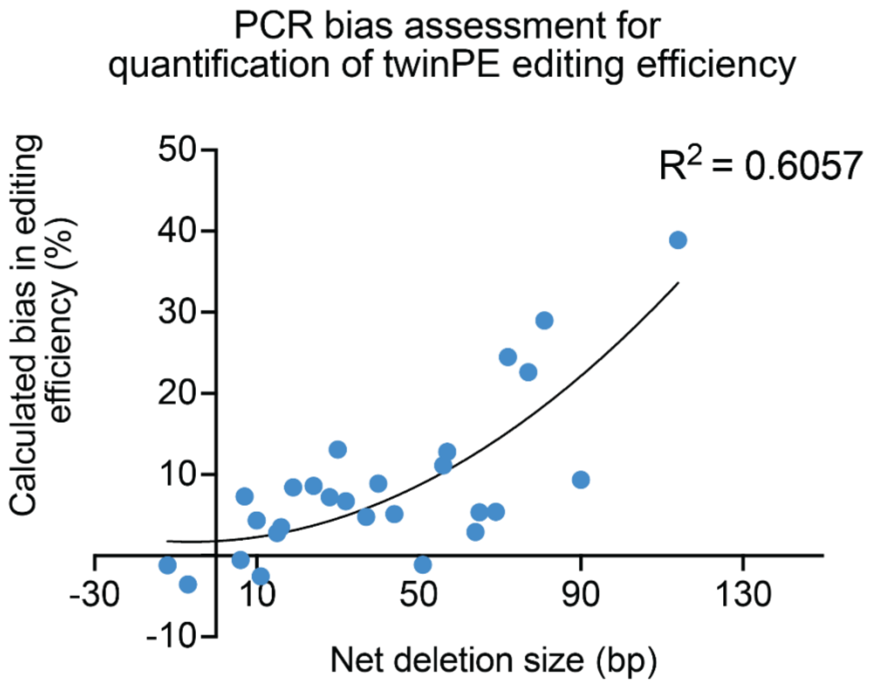

Each point in the above graph represents the mean value of bias calculated from three independent biological replicates. The percentage of bias in editing efficiency was defined by the following equation: 100 x (A – B) ÷ B, where “A” represents the editing efficiency calculated from total read counts without UMI-based deduplication, and “B” represents the editing efficiency calculated after UMI-based deduplication. For example, if the non-deduplicated editing efficiency is 60% and the deduplicated editing efficiency is 46%, the calculated bias is 30%, i.e., the deduplicated editing efficiency is 1.3-fold lower than the non-deduplicated editing efficiency. The bias in editing efficiency is plotted above as a function of the net deletion between the starting and edited alleles, which shows a positive correlation (a second-order polynomial trend line, $R^2$ = 0.6057).

**Supplementary Note 2.** Representative plot of FACS gating for GFP reporter assay

25,000 HEK293T stable GFP reporter cells were seeded into 48-well poly-D-lysine coated plates (Corning). 16-24 h post-seeding, cells were transfected with 1uL of Lipofectamine 2000 (Thermo Fisher Scientific) using the protocol described in the methods section and 750 ng of PE2, 250 ng AAVS1 targeting pegRNAs (62.5 ng each), and Bxb1 plasmid DNA (100 ng, 200 ng, 500 ng, or 1000 ng). The untreated and twinPE+Bxb1 treated cells were cultured for 72 hours, and then collected for flow cytometry analysis. HEK293T stable GFP reporter cells were first gated (Gate A) based on forward (FSC-A) and side scattering (SSC-A) to remove dead cells and other debris. A second gate (Gate B) was used to select singlets based on FSC-H and FSC-A. Finally, GFP positive and GFP negative cells were gated (Gate C) and analyzed via FITC channel.

**a**

A: Gating on population - live cells

B: Doublet Exclusion

C: Gate GFP positive cells

Untreated sample

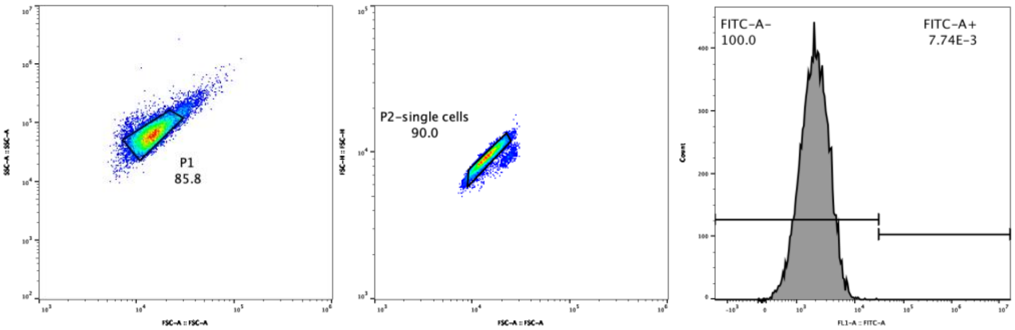

Gates and statistics

| | | |
|---|---|---|
| ▼ SS35.fcs | | 16659 |
| ▼ P1 | 85.8 | 14286 |
| ▼ P2-single cells | 90.0 | 12853 |
| FITC-A+ | 7.78E-3 | 1 |
| FITC-A- | 100.0 | 12852 |

**b**

C: Gate GFP positive cells

twinPE + Bxb1 (500ng) treated sample

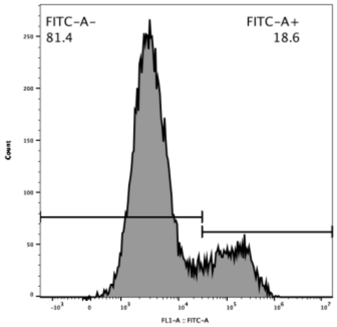

Gates and statistics

| | | |
|---|---|---|
| ▼ S7.fcs | | 18633 |
| ▼ P1 | 73.0 | 13609 |
| ▼ P2-single cells | 85.8 | 11671 |
| FITC-A+ | 18.6 | 2170 |
| FITC-A- | 81.4 | 9501 |

**Supplementary Note 3.** Analysis of twinPE design principles

To identify design principles that lead to efficient twinPE editing, we analyzed our dataset of *attP* insertions at *AAVS1* and *attB* insertions at *CCR5*. For each spacer pair, we calculated nick-to-nick distances between predicted pegRNA-induced nick sites, the DeepCas9 score[3] for each spacer, and the predicted $T_m$ of the pegRNA primer binding sequence (PBS) for *AAVS1*. For comparing nick-to-nick distances and DeepCas9 scores, the highest efficiency pegRNA pair for each spacer pair was chosen for analysis (*i.e.*, the optimal individual pegRNAs were used). For analyzing the influence of predicted spacer activities, the average of the individual DeepCas9 scores for each spacer was used. For comparing predicted $T_m$ values for PBS variants, the editing efficiency of each pegRNA was normalized to the maximum editing achieved within the spacer pair group. The editing efficiency for a given PBS variant was then averaged across the three paired pegRNA editing efficiencies (each experiment with the paired spacer). $T_m$ was calculated according to the following formula[4]: $T_m = 4N_{G \cdot C} + 2N_{A \cdot T}$.

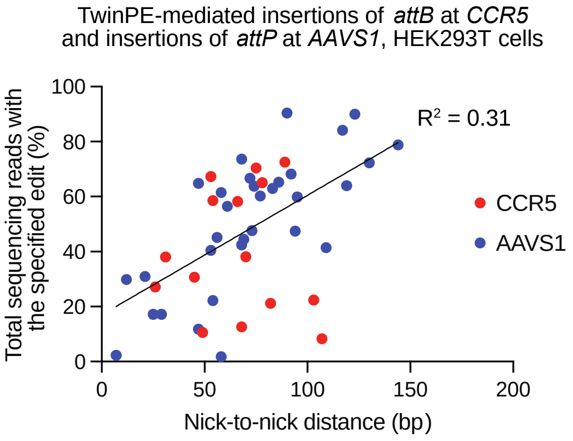

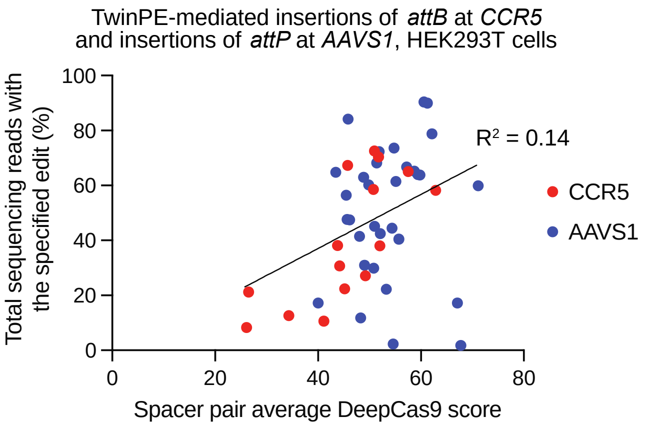

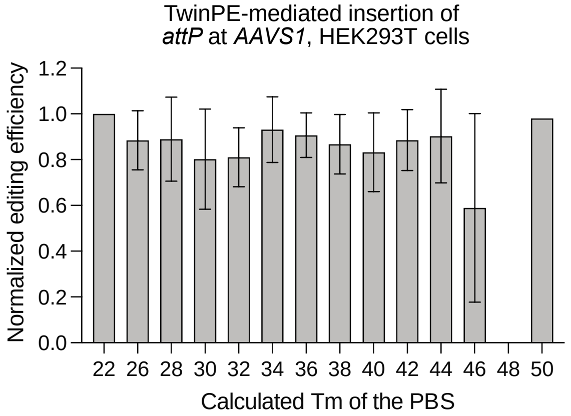

**Supplementary Sequences 1**. Sequences of plasmid donor DNA sequences harboring Bxb1 recombination sites

*att* site: highlighted in yellow
Promoter-less EGFP: highlighted in green
EF1a promoter: underlined
PuroR: highlighted in red
BFP: highlighted in blue
KanR: highlighted in grey

***attB*-Puro-GA donor DNA:**
gatgccagctcattcctcccactcatgatctatagatcccccgggctgcaggaattctacccactctgtcgataccccaccgagacc
ccattggggccaatacgcccgcgtttcttccttttccccaccccaccccccaagttcgggtgaaggcccagggctcgcagccaacg
tcggggcggcaagcttacatcgagatcccggcttgtcgacgacggcggactccgtcgtcaggatcatccgtgagcaagggcga
ggagctgttcaccggggtggtgcccatcctggtcgagctggacggcgacgtaaacggccacaagttcagcgtgtccggcgagg
gcgagggcgatgccacctacggcaagctgaccctgaagttcatctgcaccaccggcaagctgcccgtgccctggcccaccctc
gtgaccaccctgacctacggcgtgcagtgcttcagccgctaccccgaccacatgaagcagcacgacttcttcaagtccgccatgc
ccgaaggctacgtccaggagcgcaccatcttcttcaaggacgacggcaactacaagacccgcgccgaggtgaagttcgaggg
cgacaccctggtgaaccgcatcgagctgaagggcatcgacttcaaggaggacggcaacatcctggggcacaagctggagtac
aactacaacagccacaacgtctatatcatggccgacaagcagaagaacggcatcaaggtgaacttcaagatccgccacaaca
tcgaggacggcagcgtgcagctcgccgaccactaccagcagaacacccccatcggcgacggccccgtgctgctgcccgaca
accactacctgagcacccagtccgccctgagcaaagaccccaacgagaagcgcgatcacatggtcctgctggagttcgtgacc
gccgccgggatcactctcggcatggacgagctgtacaagagcggcctgaggagcagagcccaggcgagcaacagcgccgt
ggacgccaccatgggcgatcgcccgggaattgactagtgcggccgcctaggatccatgccgatagcgttggttgagtggataac
cgtattaccgccaagcttatgcatgtgcccgtcagtgggcagagcgcacatcgcccacagtccccgagaagttggggggaggg
gtcggcaattgaaccggtgcctagagaaggtggcgcggggtaaactgggaaagtgatgtcgtgtactggctccgcctttttcccga
gggtgggggagaaccgtatataagtgcagtagtcgccgtgaacgttctttttcgcaacgggtttgccgccagaacacaggtaagtg
ccgtgtgtggttcccgcgggcctggcctctttacgggttatggcccttgcgtgccttgaattacttccacctggctgcagtacgtgattctt
gatcccgagcttcgggttggaagtgggtgggagagttcgaggccttgcgcttaaggagccccttcgcctcgtgcttgagttgaggcc
tggcctgggcgctggggccgccgcgtgcgaatctggtggcaccttcgcgcctgtctcgctgctttcgataagtctctagccatttaaa
atttttgatgacctgctgcgacgctttttttctggcaagatagtcttgtaaatgcgggccaagatctgcacactggtatttcggtttttgggg
ccgcgggcggcgacggggcccgtgcgtcccagcgcacatgttcggcgaggcggggcctgcgagcgcggccaccgagaatc
ggacgggggtagtctcaagctggccggcctgctctggtgcctggcctcgcgccgccgtgtatcgccccgccctgggcggcaagg
ctggcccggtcggcaccagttgcgtgagcggaaagatggccgcttcccggccctgctgcagggagctcaaaatggaggacgc
ggcgctcgggagagcgggcgggtgagtcacccacacaaaggaaaagggcctttccgtcctcagccgtcgcttcatgtgactcca
cggagtaccgggcgccgtccaggcacctcgattagttctcgcgcttttggagtacgtcgtctttaggttggggggaggggttttatgc
gatggagtttccccacactgagtgggtggagactgaagttaggccagcttggcacttgatgtaattctccttggaatttgccctttttga
gtttggatcttggttcattctcaagcctcagacagtggttcaaagtttttttcttccatttcaggtgtcgtgagctagcccaccatgaccga
gtacaagcccacggtgcgcctcgccacccgcgacgacgtccccgggccgtacgcaccctcgccgccgcgttcgccgactacc
ccgccacgcgccacaccgtcgacccggaccgccacatcgagcgggtcaccgagctgcaagaactcttcctcacgcgcgtcgg
gctcgacatcggcaaggtgtgggtcgcggacgacggcgccgcggtggcggtctggaccacgccggagagcgtcgaagcggg
ggcggtgttcgccgagatcggcccgcgcatggccgagttgagcggttcccggctggccgcgcagcaacagatggaaggcctcc
tggcgccgcaccggcccaaggagcccgcgtggttcctggccaccgtcggcgtctcgcccgaccaccagggcaagggtctggg
cagcgccgtcgtgctccccggagtggaggcggccgagcgcgccggggtgcccgccttcctggagacctccgcgccccgcaac
ctccccttctacgagcggctcggcttcaccgtcaccgccgacgtcgaggtgcccgaaggaccgcgcacctggtgcatgacccgc
aagcccggtgccggatcgggagagggcagaggaagtctgctaacatgcggtgacgtcgaggagaatcctggcccaccggtcg
ccaccagcgagctgattaaggagaacatgcacatgaagctgtacatggagggcaccgtggacaaccatcacttcaagtgcaca
tccgagggcgaaggcaagccctacgagggcacccagaccatgagaatcaaggtggtcgagggcggccctctccccttcgcctt
cgacatcctggctactagcttcctctacggcagcaagaccttcatcaaccacacccagggcatccccgacttcttcaagcagtcctt

ccctgagggcttcacatgggagagagtcaccacatacgaagacgggggcgtgctgaccgctacccaggacaccagcctccag
gacggctgcctcatctacaacgtcaagatcagaggggtgaacttcacatccaacggccctgtgatgcagaagaaaacactcgg
ctgggaggccttcaccgagacgctgtaccccgctgacggcggcctggaaggcagaaacgacatggccctgaagctcgtgggc
gggagccatctgatcgcaaacatcaagaccacatatagatccaagaaacccgctaagaacctcaagatgcctggcgtctactat
gtggactacagactggaaagaatcaaggaggccaacaacgagacctacgtcgagcagcacgaggtggcagtggccagata
ctgcgacctccctagcaaactggggcacaagcttaattaagaattctctagaggatccagacatgataagatacattgatgagtttg
gacaaaccacaactagaatgcagtgaaaaaaatgctttatttgtgaaatttgtgatgctattgctttatttgtaaccattataagctgca
ataaacaagttaacaacaacaattgcattcattttatgtttcaggttcagggggaggtgtgggaggttttttaaagcaagtaaaacctc
tacaaatgtggtatggctgattatgatcctgcaagcctcgtcgccgcggtttattctgttgacaattaatcatcggcatagtatatcggc
atagtataatacgacaaggtgaggaactaaaccatgggatcggccattgaacaagatggattgcacgcaggttctccggccgctt
gggtggagaggctattcggctatgactgggcacaacagacgatcggctgctctgatgccgccgtgttccggctgtcagcgcaggg
gcgcccggttctttttgtcaagaccgacctgtccggtgccctgaatgaactgcaggacgaggcagcgcggctatcgtggctggcca
cgacgggcgttccttgcgcagctgtgctcgacgttgtcactgaagcgggaagggactggctgctattgggcgaagtgccggggca
ggatctcctgtcatctcaccttgctcctgccgagaaagtatccatcatggctgatgcaatgcggcggctgcatacgcttgatccggct
acctgcccattcgaccaccaagcgaaacatcgcatcgagcgagcacgtactcggatggaagccggtcttgtcgatcaggatgat
ctggacgaagagcatcaggggctcgcgccagccgaactgttcgccaggctcaaggcgcgcatgcccgacggcgaggatctcg
tcgtgacccatggcgatgcctgcttgccgaatatcatggtggaaaatggccgcttttctggattcatcgactgtggccggctgggtgtg
gcggaccgctatcaggacatagcgttggctacccgtgatattgctgaagagcttggcggcgaatgggctgaccgcttcctcgtgctt
tacggtatcgccgcccccgattcgcagcgcatcgccttctatcgccttcttgacgagttcttctgagcgggactctggggttcgaataa
agaccgaccaagcgacgtctgagagctccctggcgaattcggtaccaataaaagagctttattttcatgatctgtgtgttggtttttgg
ccgcgttgctggcgtttttccataggctccgcccccctgacgagcatcacaaaaatcgacgctcaagtcagaggtggcgaaaccc
gacaggactataaagataccaggcgtttccccctggaagctccctcgtgcgctctcctgttccgaccctgccgcttaccggatacct
gtccgcctttctcccttcgggaagcgtggcgctttctcatagctcacgctgtaggtatctcagttcggtgtaggtcgttcgctccaagctg
ggctgtgtgcacgaaccccccgttcagcccgaccgctgcgccttatccggtaactatcgtcttgagtccaacccggtaagacacga
cttatcgccactggcagcagccactggtaacaggattagcagagcgaggtatgtaggcggtgctacagagttcttgaagtggtgg
cctaactacggctacactagaagaacagtatttggtatctgcgctctgctgaagccagttaccttcggaaaaagagttggtagctctt
gatccggcaaacaaaccaccgctggtagcggtggtttttttgtttgcaagcagcagattacgcgcagaaaaaaaggatctcaaga
agatcctttgatcttttctagtgtgcg

***attB*-Puro donor DNA:**
gatgccagctcattcctcccactcatgatctatagatcccccgggctgcaggaattctacccactctgtcgataccccaccgagacc
ccattggggccaatacgcccgcgtttcttccttttccccaccccaccccccaagttcgggtgaaggcccagggctcgcagccaacg
tcggggcggcaagcttacatcgagatcccggcttgtcgacgacggcggtctccgtcgtcaggatcatccgtgagcaagggcgag
gagctgttcaccggggtggtgcccatcctggtcgagctggacggcgacgtaaacggccacaagttcagcgtgtccggcgaggg
cgagggcgatgccacctacggcaagctgaccctgaagttcatctgcaccaccggcaagctgcccgtgccctggcccaccctcgt
gaccaccctgacctacggcgtgcagtgcttcagccgctaccccgaccacatgaagcagcacgacttcttcaagtccgccatgcc
cgaaggctacgtccaggagcgcaccatcttcttcaaggacgacggcaactacaagacccgcgccgaggtgaagttcgagggc
gacaccctggtgaaccgcatcgagctgaagggcatcgacttcaaggaggacggcaacatcctggggcacaagctggagtaca
actacaacagccacaacgtctatatcatggccgacaagcagaagaacggcatcaaggtgaacttcaagatccgccacaacat
cgaggacggcagcgtgcagctcgccgaccactaccagcagaacacccccatcggcgacggccccgtgctgctgcccgacaa
ccactacctgagcacccagtccgccctgagcaaagaccccaacgagaagcgcgatcacatggtcctgctggagttcgtgaccg
ccgccgggatcactctcggcatggacgagctgtacaagagcggcctgaggagcagagcccaggcgagcaacagcgccgtg
gacgccaccatgggcgatcgcccgggaattgactagtgcggccgcctaggatccatgccgatagcgttggttgagtggataacc
gtattaccgccaagcttatgcatgtgcccgtcagtgggcagagcgcacatcgcccacagtccccgagaagttggggggaggggt
cggcaattgaaccggtgcctagagaaggtggcgcggggtaaactgggaaagtgatgtcgtgtactggctccgcctttttcccgag
ggtgggggagaaccgtatataagtgcagtagtcgccgtgaacgttctttttcgcaacgggtttgccgccagaacacaggtaagtgc
cgtgtgtggttcccgcgggcctggcctctttacgggttatggcccttgcgtgccttgaattacttccacctggctgcagtacgtgattcttg
atcccgagcttcgggttggaagtgggtgggagagttcgaggccttgcgcttaaggagccccttcgcctcgtgcttgagttgaggcct
ggcctgggcgctggggccgccgcgtgcgaatctggtggcaccttcgcgcctgtctcgctgctttcgataagtctctagccatttaaaa

tttttgatgacctgctgcgacgcttttttctggcaagatagtcttgtaaatgcgggccaagatctgcacactggtatttcggtttttggggc
cgcgggcggcgacggggcccgtgcgtcccagcgcacatgttcggcgaggcggggcctgcgagcgcggccaccgagaatcg
gacgggggtagtctcaagctggccggcctgctctggtgcctggcctcgcgccgccgtgtatcgccccgccctgggcggcaaggct
ggcccggtcggcaccagttgcgtgagcggaaagatggccgcttcccggccctgctgcagggagctcaaaatggaggacgcgg
cgctcgggagagcgggcgggtgagtcacccacacaaaggaaaagggcctttccgtcctcagccgtcgcttcatgtgactccacg
gagtaccgggcgccgtccaggcacctcgattagttctcgcgcttttggagtacgtcgtctttaggttggggggaggggttttatgcgat
ggagtttccccacactgagtgggtggagactgaagttaggccagcttggcacttgatgtaattctccttggaatttgccctttttgagttt
ggatcttggttcattctcaagcctcagacagtggttcaaagtttttttcttccatttcaggtgtcgtgagctagcccaccatgaccgagta
caagcccacggtgcgcctcgccacccgcgacgacgtccccgggccgtacgcaccctcgccgccgcgttcgccgactacccc
gccacgcgccacaccgtcgacccggaccgccacatcgagcgggtcaccgagctgcaagaactcttcctcacgcgcgtcgggc
tcgacatcggcaaggtgtgggtcgcggacgacggcgccgcggtggcggtctggaccacgccggagagcgtcgaagcggggg
cggtgttcgccgagatcggcccgcgcatggccgagttgagcggttcccggctggccgcgcagcaacagatggaaggcctcctg
gcgccgcaccggcccaaggagcccgcgtggttcctggccaccgtcggcgtctcgcccgaccaccagggcaagggtctgggca
gcgccgtcgtgctccccggagtggaggcggccgagcgcgccggggtgcccgccttcctggagacctccgcgccccgcaacctc
cccttctacgagcggctcggcttcaccgtcaccgccgacgtcgaggtgcccgaaggaccgcgcacctggtgcatgacccgcaa
gcccggtgccggatcgggagagggcagaggaagtctgctaacatgcggtgacgtcgaggagaatcctggcccaccggtcgcc
accagcgagctgattaaggagaacatgcacatgaagctgtacatggagggcaccgtggacaaccatcacttcaagtgcacatc
cgagggcgaaggcaagccctacgagggcacccagaccatgagaatcaaggtggtcgagggcggccctctcccctttcgccttc
gacatcctggctactagcttcctctacggcagcaagaccttcatcaaccacacccagggcatccccgacttcttcaagcagtccttc
cctgagggcttcacatgggagagagtcaccacatacgaagacgggggcgtgctgaccgctacccaggacaccagcctccagg
acggctgcctcatctacaacgtcaagatcagaggggtgaacttcacatccaacggccctgtgatgcagaagaaaacactcggct
gggaggccttcaccgagacgctgtaccccgctgacggcggcctggaaggcagaaacgacatggccctgaagctcgtgggcg
ggagccatctgatcgcaaacatcaagaccacatatagatccaagaaacccgctaagaacctcaagatgcctggcgtctactatg
tggactacagactggaaagaatcaaggaggccaacaacgagacctacgtcgagcagcacgaggtggcagtggccagatact
gcgacctccctagcaaactggggcacaagcttaattaagaattctctagaggatccagacatgataagatacattgatgagtttgg
acaaaccacaactagaatgcagtgaaaaaaatgctttatttgtgaaatttgtgatgctattgctttatttgtaaccattataagctgcaat
aaacaagttaacaacaacaattgcattcattttatgtttcaggttcagggggaggtgtgggaggttttttaaagcaagtaaaacctcta
caaatgtggtatggctgattatgatcctgcaagcctcgtcgccgcggtttattctgttgacaattaatcatcggcatagtatatcggcat
agtataatacgacaaggtgaggaactaaaccatgggatcggccattgaacaagatggattgcacgcaggttctccggccgcttg
ggtggagaggctattcggctatgactgggcacaacagacgatcggctgctctgatgccgccgtgttccggctgtcagcgcagggg
cgcccggttctttttgtcaagaccgacctgtccggtgccctgaatgaactgcaggacgaggcagcgcggctatcgtggctggccac
gacgggcgttccttgcgcagctgtgctcgacgttgtcactgaagcgggaagggactggctgctattgggcgaagtgccggggca
ggatctcctgtcatctcaccttgctcctgccgagaaagtatccatcatggctgatgcaatgcggcggctgcatacgcttgatccggct
acctgcccattcgaccaccaagcgaaacatcgcatcgagcgagcacgtactcggatggaagccggtcttgtcgatcaggatgat
ctggacgaagagcatcaggggctcgcgccagccgaactgttcgccaggctcaaggcgcgcatgcccgacggcgaggatctcg
tcgtgacccatggcgatgcctgcttgccgaatatcatggtggaaaatggccgcttttctggattcatcgactgtggccggctgggtgtg
gcggaccgctatcaggacatagcgttggctacccgtgatattgctgaagagcttggcggcgaatgggctgaccgcttcctcgtgctt
tacggtatcgccgctcccgattcgcagcgcatcgccttctatcgccttcttgacgagttcttctgagcgggactctggggttcgaataa
agaccgaccaagcgacgtctgagagctccctggcgaattcggtaccaataaaagagctttattttcatgatctgtgtgttggtttttgg
ccgcgttgctggcgtttttccataggctccgcccccctgacgagcatcacaaaaatcgacgctcaagtcagaggtggcgaaaccc
gacaggactataaagataccaggcgtttccccctggaagctccctcgtgcgctctcctgttccgaccctgccgcttaccggatacct
gtccgcctttctcccttcgggaagcgtggcgctttctcatagctcacgctgtaggtatctcagttcggtgtaggtcgttcgctccaagctg
ggctgtgtgcacgaaccccccgttcagcccgaccgctgcgccttatccggtaactatcgtcttgagtccaacccggtaagacacga
cttatcgccactggcagcagccactggtaacaggattagcagagcgaggtatgtaggcggtgctacagagttcttgaagtggtgg
cctaactacggctacactagaagaacagtatttggtatctgcgctctgctgaagccagttaccttcggaaaaagagttggtagctctt
gatccggcaaacaaaccaccgctggtagcggtggtttttttgtttgcaagcagcagattacgcgcagaaaaaaaggatctcaaga
agatcctttgatcttttctagtgtgcg

***attP*-Puro-GA donor DNA:**

gatgccagctcattcctcccactcatgatctatagatcccccgggctgcaggaattctacccactctgtcgataccccaccgagacc
ccattggggccaatacgcccgcgtttcttccttttccccaccccaccccccaagttcgggtgaaggcccagggctcgcagccaacg
tcggggcggcaagcttacatcgagatcccggtttgtctggtcaaccaccgcggactcagtggtgtacggtacaaacccgtgagc
aagggcgaggagctgttcaccggggtggtgcccatcctggtcgagctggacggcgacgtaaacggccacaagttcagcgtgtc
cggcgagggcgagggcgatgccacctacggcaagctgaccctgaagttcatctgcaccaccggcaagctgcccgtgccctgg
cccaccctcgtgaccaccctgacctacggcgtgcagtgcttcagccgctaccccgaccacatgaagcagcacgacttcttcaagt
ccgccatgcccgaaggctacgtccaggagcgcaccatcttcttcaaggacgacggcaactacaagacccgcgccgaggtgaa
gttcgagggcgacaccctggtgaaccgcatcgagctgaagggcatcgacttcaaggaggacggcaacatcctggggcacaag
ctggagtacaactacaacagccacaacgtctatatcatggccgacaagcagaagaacggcatcaaggtgaacttcaagatccg
ccacaacatcgaggacggcagcgtgcagctcgccgaccactaccagcagaacacccccatcggcgacggccccgtgctgct
gcccgacaaccactacctgagcacccagtccgccctgagcaaagaccccaacgagaagcgcgatcacatggtcctgctggag
ttcgtgaccgccgccgggatcactctcggcatggacgagctgtacaagagcggcctgaggagcagagcccaggcgagcaac
agcgccgtggacgccaccatgggcgatcgcccgggaattgactagtgcggccgcctaggatccatgccgatagcgttggttgag
tggataaccgtattaccgccaagcttatgcatgtgcccgtcagtgggcagagcgcacatcgcccacagtccccgagaagttggg
gggaggggtcggcaattgaaccggtgcctagagaaggtggcgcggggtaaactgggaaagtgatgtcgtgtactggctccgcct
ttttcccgagggtgggggagaaccgtatataagtgcagtagtcgccgtgaacgttctttttcgcaacgggtttgccgccagaacaca
ggtaagtgccgtgtgtggttcccgcgggcctggcctctttacgggttatggcccttgcgtgccttgaattacttccacctggctgcagta
cgtgattcttgatcccgagcttcgggttggaagtgggtgggagagttcgaggccttgcgcttaaggagccccttcgcctcgtgcttga
gttgaggcctggcctgggcgctggggccgccgcgtgcgaatctggtggcaccttcgcgcctgtctcgctgctttcgataagtctctag
ccatttaaaatttttgatgacctgctgcgacgctttttttctggcaagatagtcttgtaaatgcgggccaagatctgcacactggtatttcg
gtttttggggccgcgggcggcgacggggcccgtgcgtcccagcgcacatgttcggcgaggcggggcctgcgagcgcggccac
cgagaatcggacgggggtagtctcaagctggccggcctgctctggtgcctggcctcgcgccgccgtgtatcgccccgccctgggc
ggcaaggctggcccggtcggcaccagttgcgtgagcggaaagatggccgcttcccggccctgctgcagggagctcaaaatgg
aggacgcggcgctcgggagagcgggcgggtgagtcacccacacaaaggaaaagggcctttccgtcctcagccgtcgcttcat
gtgactccacggagtaccgggcgccgtccaggcacctcgattagttctcgcgcttttggagtacgtcgtctttaggttgggggagg
ggttttatgcgatggagtttccccacactgagtgggtggagactgaagttaggccagcttggcacttgatgtaattctccttggaatttg
ccctttttgagtttggatcttggttcattctcaagcctcagacagtggttcaaagtttttttcttccatttcaggtgtcgtgagctagcccacc
atgaccgagtacaagcccacggtgcgcctcgccacccgcgacgacgtccccgggccgtacgcaccctcgccgccgcgttcg
ccgactaccccgccacgcgccacaccgtcgacccggaccgccacatcgagcgggtcaccgagctgcaagaactcttcctcac
gcgcgtcgggctcgacatcggcaaggtgtgggtcgcggacgacggcgccgcggtggcggtctggaccacgccggagagcgt
cgaagcgggggcggtgttcgccgagatcggcccgcgcatggccgagttgagcggttcccggctggccgcgcagcaacagatg
gaaggcctcctggcgccgcaccggcccaaggagcccgcgtggttcctggccaccgtcggcgtctcgcccgaccaccagggca
agggtctgggcagcgccgtcgtgctccccggagtggaggcggccgagcgcgccggggtgcccgccttcctggagacctccgc
gccccgcaacctccccttctacgagcggctcggcttcaccgtcaccgccgacgtcgaggtgcccgaaggaccgcgcacctggtg
catgacccgcaagcccggtgccggatcgggagagggcagaggaagtctgctaacatgcggtgacgtcgaggagaatcctggc
ccaccggtcgccaccagcgagctgattaaggagaacatgcacatgaagctgtacatggagggcaccgtggacaaccatcactt
caagtgcacatccgagggcgaaggcaagccctacgagggcacccagaccatgagaatcaaggtggtcgagggcggccctct
ccccttcgccttcgacatcctggctactagcttcctctacggcagcaagaccttcatcaaccacacccagggcatccccgacttcttc
aagcagtccttccctgagggcttcacatgggagagagtcaccacatacgaagacgggggcgtgctgaccgctacccaggaca
ccagcctccaggacggctgcctcatctacaacgtcaagatcagaggggtgaacttcacatccaacggccctgtgatgcagaaga
aaacactcggctgggaggccttcaccgagacgctgtaccccgctgacggcggcctggaaggcagaaacgacatggccctgaa
gctcgtgggcgggagccatctgatcgcaaacatcaagaccacatatagatccaagaaacccgctaagaacctcaagatgcctg
gcgtctactatgtggactacagactggaaagaatcaaggaggccaacaacgagacctacgtcgagcagcacgaggtggcagt
ggccagatactgcgacctccctagcaaactggggcacaagcttaattaagaattctctagaggatccagacatgataagatacat
tgatgagtttggacaaaccacaactagaatgcagtgaaaaaaatgctttatttgtgaaatttgtgatgctattgctttatttgtaaccatta
taagctgcaataaacaagttaacaacaacaattgcattcattttatgtttcaggttcagggggaggtgtgggaggttttttaaagcaag
taaaacctctacaaatgtggtatggctgattatgatcctgcaagcctcgtcgccgcggtttattctgttgacaattaatcatcggcatag
tatatcggcatagtataatacgacaaggtgaggaactaaaccatgggatcggccattgaacaagatggattgcacgcaggttctc
cggccgcttgggtggagaggctattcggctatgactgggcacaacagacgatcggctgctctgatgccgccgtgttccggctgtca

gcgcaggggcgcccggttctttttgtcaagaccgacctgtccggtgccctgaatgaactgcaggacgaggcagcgcggctatcgt
ggctggccacgacgggcgttccttgcgcagctgtgctcgacgttgtcactgaagcgggaagggactggctgctattgggcgaagt
gccggggcaggatctcctgtcatctcaccttgctcctgccgagaaagtatccatcatggctgatgcaatgcggcggctgcatacgct
tgatccggctacctgcccattcgaccaccaagcgaaacatcgcatcgagcgagcacgtactcggatggaagccggtcttgtcgat
caggatgatctggacgaagagcatcaggggctcgcgccagccgaactgttcgccaggctcaaggcgcgcatgcccgacggcg
aggatctcgtcgtgacccatggcgatgcctgcttgccgaatatcatggtggaaaatggccgcttttctggattcatcgactgtggccg
gctgggtgtggcggaccgctatcaggacatagcgttggctacccgtgatattgctgaagagcttggcggcgaatgggctgaccgc
ttcctcgtgctttacggtatcgccgcccccgattcgcagcgcatcgccttctatcgccttcttgacgagttcttctgagcgggactctggg
gttcgaataaagaccgaccaagcgacgtctgagagctccctggcgaattcggtaccaataaaagagctttattttcatgatctgtgt
gttggtttttggccgcgttgctggcgtttttccataggctccgcccccctgacgagcatcacaaaaatcgacgctcaagtcagaggtg
gcgaaacccgacaggactataaagataccaggcgtttccccctggaagctccctcgtgcgctctcctgttccgaccctgccgctta
ccggatacctgtccgcctttctcccttcgggaagcgtggcgctttctcatagctcacgctgtaggtatctcagttcggtgtaggtcgttcg
ctccaagctgggctgtgtgcacgaaccccccgttcagcccgaccgctgcgccttatccggtaactatcgtcttgagtccaacccggt
aagacacgacttatcgccactggcagcagccactggtaacaggattagcagagcgaggtatgtaggcggtgctacagagttctt
gaagtggtggcctaactacggctacactagaagaacagtatttggtatctgcgctctgctgaagccagttaccttcggaaaaagag
ttggtagctcttgatccggcaaacaaaccaccgctggtagcggtggtttttttgtttgcaagcagcagattacgcgcagaaaaaaag
gatctcaagaagatcctttgatcttttctagtgtgcg

***attP*-Puro donor DNA:**
gatgccagctcattcctcccactcatgatctatagatcccccgggctgcaggaattctacccactctgtcgataccccaccgagacc
ccattggggccaatacgcccgcgtttcttccttttccccaccccaccccccaagttcgggtgaaggcccagggctcgcagccaacg
tcggggcggcaagcttacatcgagatcccggtttgtctggtcaaccaccgcggtctcagtggtgtacggtacaaacccgtgagc
aagggcgaggagctgttcaccggggtggtgcccatcctggtcgagctggacggcgacgtaaacggccacaagttcagcgtgtc
cggcgagggcgagggcgatgccacctacggcaagctgaccctgaagttcatctgcaccaccggcaagctgcccgtgccctgg
cccaccctcgtgaccaccctgacctacggcgtgcagtgcttcagccgctaccccgaccacatgaagcagcacgacttcttcaagt
ccgccatgcccgaaggctacgtccaggagcgcaccatcttcttcaaggacgacggcaactacaagacccgcgccgaggtgaa
gttcgagggcgacaccctggtgaaccgcatcgagctgaagggcatcgacttcaaggaggacggcaacatcctggggcacaag
ctggagtacaactacaacagccacaacgtctatatcatggccgacaagcagaagaacggcatcaaggtgaacttcaagatccg
ccacaacatcgaggacggcagcgtgcagctcgccgaccactaccagcagaacacccccatcggcgacggccccgtgctgct
gcccgacaaccactacctgagcacccagtccgccctgagcaaagaccccaacgagaagcgcgatcacatggtcctgctggag
ttcgtgaccgccgccgggatcactctcggcatggacgagctgtacaagagcggcctgaggagcagagcccaggcgagcaac
agcgccgtggacgccaccatgggcgatcgcccgggaattgactagtgcggccgcctaggatccatgccgatagcgttggttgag
tggataaccgtattaccgccaagcttatgcatgtgcccgtcagtgggcagagcgcacatcgcccacagtccccgagaagttggg
gggaggggtcggcaattgaaccggtgcctagagaaggtggcgcggggtaaactgggaaagtgatgtcgtgtactggctccgcct
ttttcccgagggtgggggagaaccgtatataagtgcagtagtcgccgtgaacgttctttttcgcaacgggtttgccgccagaacaca
ggtaagtgccgtgtgtggttcccgcgggcctggcctctttacgggttatggcccttgcgtgccttgaattacttccacctggctgcagta
cgtgattcttgatcccgagcttcgggttggaagtgggtgggagagttcgaggccttgcgcttaaggagccccttcgcctcgtgcttga
gttgaggcctggcctgggcgctggggccgccgcgtgcgaatctggtggcaccttcgcgcctgtctcgctgctttcgataagtctctag
ccatttaaaatttttgatgacctgctgcgacgctttttttctggcaagatagtcttgtaaatgcgggccaagatctgcacactggtatttcg
gtttttggggccgcgggcggcgacggggcccgtgcgtcccagcgcacatgttcggcgaggcggggcctgcgagcgcggccac
cgagaatcggacgggggtagtctcaagctggccggcctgctctggtgcctggcctcgcgccgccgtgtatcgccccgccctgggc
ggcaaggctggcccggtcggcaccagttgcgtgagcggaaagatggccgcttcccggccctgctgcagggagctcaaaatgg
aggacgcggcgctcgggagagcgggcgggtgagtcacccacacaaaggaaaagggcctttccgtcctcagccgtcgcttcat
gtgactccacggagtaccgggcgccgtccaggcacctcgattagttctcgcgcttttggagtacgtcgtctttaggttggggggagg
ggttttatgcgatggagtttccccacactgagtgggtggagactgaagttaggccagcttggcacttgatgtaattctccttggaatttg
ccctttttgagtttggatcttggttcattctcaagcctcagacagtggttcaaagtttttttcttccatttcaggtgtcgtgagctagcccacc
atgaccgagtacaagcccacggtgcgcctcgccacccgcgacgacgtccccgggccgtacgcaccctcgccgccgcgttcg
ccgactaccccgccacgcgccacaccgtcgacccggaccgccacatcgagcgggtcaccgagctgcaagaactcttcctcac
gcgcgtcgggctcgacatcggcaaggtgtgggtcgcggacgacggcgccgcggtggcggtctggaccacgccggagagcgt

cgaagcgggggcggtgttcgccgagatcggcccgcgcatggccgagttgagcggttcccggctggccgcgcagcaacagatg
gaaggcctcctggcgccgcaccggcccaaggagcccgcgtggttcctggccaccgtcggcgtctcgcccgaccaccagggca
agggtctgggcagcgccgtcgtgctccccggagtggaggcggccgagcgcgccggggtgcccgccttcctggagacctccgc
gccccgcaacctccccttctacgagcggctcggcttcaccgtcaccgccgacgtcgaggtgcccgaaggaccgcgcacctggtg
catgacccgcaagcccggtgccggatcgggagagggcagaggaagtctgctaacatgcggtgacgtcgaggagaatcctggc
ccaccggtcgccaccagcgagctgattaaggagaacatgcacatgaagctgtacatggagggcaccgtggacaaccatcactt
caagtgcacatccgagggcgaaggcaagccctacgagggcacccagaccatgagaatcaaggtggtcgagggcggccctct
ccccttcgccttcgacatcctggctactagcttcctctacggcagcaagaccttcatcaaccacacccagggcatccccgacttcttc
aagcagtccttccctgagggcttcacatgggagagagtcaccacatacgaagacgggggcgtgctgaccgctacccaggaca
ccagcctccaggacggctgcctcatctacaacgtcaagatcagaggggtgaacttcacatccaacggccctgtgatgcagaaga
aaacactcggctgggaggccttcaccgagacgctgtaccccgctgacggcggcctggaaggcagaaacgacatggccctgaa
gctcgtgggcgggagccatctgatcgcaaacatcaagaccacatatagatccaagaaacccgctaagaacctcaagatgcctg
gcgtctactatgtggactacagactggaaagaatcaaggaggccaacaacgagacctacgtcgagcagcacgaggtggcagt
ggccagatactgcgacctccctagcaaactggggcacaagcttaattaagaattctctagaggatccagacatgataagatacat
tgatgagtttggacaaaccacaactagaatgcagtgaaaaaaatgctttatttgtgaaatttgtgatgctattgctttatttgtaaccatta
taagctgcaataaacaagttaacaacaacaattgcattcattttatgtttcaggttcagggggaggtgtgggaggttttttaaagcaag
taaaacctctacaaatgtggtatggctgattatgatcctgcaagcctcgtcgccgcggtttattctgttgacaattaatcatcggcatag
tatatcggcatagtataatacgacaaggtgaggaactaaaccatgggatcggccattgaacaagatggattgcacgcaggttctc
cggccgcttgggtggagaggctattcggctatgactgggcacaacagacgatcggctgctctgatgccgccgtgttccggctgtca
gcgcaggggcgcccggttctttttgtcaagaccgacctgtccggtgccctgaatgaactgcaggacgaggcagcgcggctatcgt
ggctggccacgacgggcgttccttgcgcagctgtgctcgacgttgtcactgaagcgggaagggactggctgctattgggcgaagt
gccggggcaggatctcctgtcatctcaccttgctcctgccgagaaagtatccatcatggctgatgcaatgcggcggctgcatacgct
tgatccggctacctgcccattcgaccaccaagcgaaacatcgcatcgagcgagcacgtactcggatggaagccggtcttgtcgat
caggatgatctggacgaagagcatcaggggctcgcgccagccgaactgttcgccaggctcaaggcgcgcatgcccgacggcg
aggatctcgtcgtgacccatggcgatgcctgcttgccgaatatcatggtggaaaatggccgcttttctggattcatcgactgtggccg
gctgggtgtggcggaccgctatcaggacatagcgttggctacccgtgatattgctgaagagcttggcggcgaatgggctgaccgc
ttcctcgtgctttacggtatcgccgcccccgattcgcagcgcatcgccttctatcgccttcttgacgagttcttctgagcgggactctggg
gttcgaataaagaccgaccaagcgacgtctgagagctccctggcgaattcggtaccaataaaagagctttattttcatgatctgtgt
gttggtttttggccgcgttgctggcgtttttccataggctccgcccccctgacgagcatcacaaaaatcgacgctcaagtcagaggtg
gcgaaacccgacaggactataaagataccaggcgtttccccctggaagctccctcgtgcgctctcctgttccgaccctgccgctta
ccggatacctgtccgcctttctcccttcgggaagcgtggcgctttctcatagctcacgctgtaggtatctcagttcggtgtaggtcgttcg
ctccaagctgggctgtgtgcacgaaccccccgttcagcccgaccgctgcgccttatccggtaactatcgtcttgagtccaacccggt
aagacacgacttatcgccactggcagcagccactggtaacaggattagcagagcgaggtatgtaggcggtgctacagagttctt
gaagtggtggcctaactacggctacactagaagaacagtatttggtatctgcgctctgctgaagccagttaccttcggaaaaagag
ttggtagctcttgatccggcaaacaaaccaccgctggtagcggtggtttttttgtttgcaagcagcagattacgcgcagaaaaaaag
gatctcaagaagatcctttgatcttttctagtgtgcg

**attP Factor IX donor** (the vector backbone is pMC.BESPX-MCS1 from System Biosciences)

attP: highlighted in yellow
Factor IX intron 1: highlighted in grey
Factor IX exons 2-8 CDS: highlighted in green
Factor IX exon 8 3’UTR: underlined
bGH poly(A) signal: highlighted in cyan

AGGTTTGTCTGGTCAACCACCGCGGTCTCAGTGGTGTACGGTACAAACCTTTTCTTAAG
AGATGTAAAATTTTCATGATGTTTTCTTTTTTGCTAAAACTAAAGAATTATTCTTTTACATTT
CAGTTTTTCTTGATCATGAAAACGCCAACAAAATTCTGAATCGGCCAAAGAGGTATAATT
CAGGTAAATTGGAAGAGTTTGTTCAAGGGAACCTTGAGAGAGAATGTATGGAAGAAAAG
TGTAGTTTTGAAGAAGCACGAGAAGTTTTTGAAAACACTGAAAGAACAACTGAATTTTGG
AAGCAGTATGTTGATGGAGATCAGTGTGAGTCCAATCCATGTTTAAATGGCGGCAGTTG
CAAGGATGACATTAATTCCTATGAATGTTGGTGTCCCTTTGGATTTGAAGGAAAGAACTG
TGAATTAGATGTAACATGTAACATTAAGAATGGCAGATGCGAGCAGTTTTGTAAAAATAG
TGCTGATAACAAGGTGGTTTGCTCCTGTACTGAGGGATATCGACTTGCAGAAAACCAGA
AGTCCTGTGAACCAGCAGTGCCATTTCCATGTGGAAGAGTTTCTGTTTCACAAACTTCTA
AGCTCACCCGTGCTGAGACTGTTTTTCCTGATGTGGACTATGTAAATTCTACTGAAGCTG
AAACCATTTTGGATAACATCACTCAAAGCACCCAATCATTTAATGACTTCACTCGGGTTGT
TGGTGGAGAAGATGCCAAACCAGGTCAATTCCCTTGGCAGGTTGTTTTGAATGGTAAAG
TTGATGCATTCTGTGGAGGCTCTATCGTTAATGAAAAATGGATTGTAACTGCTGCCCACT
GTGTTGAAACTGGTGTTAAAATTACAGTTGTCGCAGGTGAACATAATATTGAGGAGACAG
AACATACAGAGCAAAAGCGAAATGTGATTCGAATTATTCCTCACCACAACTACAATGCAG
CTATTAATAAGTACAACCATGACATTGCCCTTCTGGAACTGGACGAACCCTTAGTGCTAA
ACAGCTACGTTACACCTATTTGCATTGCTGACAAGGAATACACGAACATCTTCCTCAAAT
TTGGATCTGGCTATGTAAGTGGCTGGGGAAGAGTCTTCCACAAAGGGAGATCAGCTTTA
GTTCTTCAGTACCTTAGAGTTCCACTTGTTGACCGAGCCACATGTCTTCGATCTACAAAG
TTCACCATCTATAACAACATGTTCTGTGCTGGCTTCCATGAAGGAGGTAGAGATTCATGT
CAAGGAGATAGTGGGGGACCCCATGTTACTGAAGTGGAAGGGACCAGTTTCTTAACTGG
AATTATTAGCTGGGGTGAAGAGTGTGCAATGAAAGGCAAATATGGAATATATACCAAGGT
ATCCCGGTATGTCAACTGGATTAAGGAAAAAACAAAGCTCACTTAATGAAAGATGGATTT
CCAAGGTTAATTCATTGGAATTGAAAATTAACAGGGCCTCTCACTAACTAATCACTTTCCC
ATCTTTTGTTAGATTTGAATATATACATTCTATGATCATTGCTTTTTCTCTTTACAGGGGAG
AATTTCATATTTTACCTGAGCAAATTGATTAGAAAATGGAACCACTAGAGGAATATAATGT
GTTAGGAAATTACAGTCATTTCTAAGGGCCCAGCCCTTGACAAAATTGTGAAGTTAAATT
CTCCACTCTGTCCATCAGATACTATGGTTCTCCACTATGGCAACTAACTCACTCAATTTTC
CCTCCTTAGCAGCATTCCATCTTCCCGATCTTCTTTGCTTCTCCAACCAAAACATCAATGT
TTATTAGTTCTGTATACAGTACAGGATCTTTGGTCTACTCTATCACAAGGCCAGTACCAC
ACTCATGAAGAAAGAACACAGGAGTAGCTGAGAGGCTAAAACTCATCAAAAACACTACT
CCTTTTCCTCTACCCTATTCCTCAATCTTTTACCTTTTCCAAATCCCAATCCCCAAATCAG
TTTTTCTCTTTCTTACTCCCTCTCTCCCTTTTACCCTCCATGGTCGTTAAAGGAGAGATGG
GGAGCATCATTCTGTTATACTTCTGTACACAGTTATACATGTCTATCAAACCCAGACTTGC
TTCCGTAGTGGAGACTTGCTTTTCAGAACATAGGGATGAAGTAAGGTGCCTGAAAAGTTT
GGGGGAAAAGTTTCTTTCAGAGAGTTAAGTTATTTTATATATATAATATATATATAAAATAT
ATAATATACAATATAAATATATAGTGTGTGTGTATGCGTGTGTGTAGACACACACGCATAC
ACACATATAATGGAAGCAATAAGCCATTCTAAGAGCTTGTATGGTTATGGAGGTCTGACT
AGGCATGATTTCACGAAGGCAAGATTGGCATATCATTGTAACTAAAAAAGCTGACATTGA
CCCAGACATATTGTACTCTTTCTAAAAATAATAATAATAATGCTAACAGAAAGAAGAGAAC

CGTTCGTTTGCAATCTACAGCTAGTAGAGACTTTGAGGAAGAATTCAACAGTGTGTCTTC
AGCAGTGTTCAGAGCCAAGCAAGAAGTTGAAGTTGCCTAGACCAGAGGACATAAGTATC
ATGTCTCCTTTAACTAGCATACCCCGAAGTGGAGAAGGGTGCAGCAGGCTCAAAGGCAT
AAGTCATTCCAATCAGCCAACTAAGTTGTCCTTTTCTGGTTTCGTGTTCACCATGGAACA
TTTTGATTATAGTTAATCCTTCTATCTTGAATCTTCTAGAGAGTTGCTGACCAACTGACGT
ATGTTTCCCTTTGTGAATTAATAAACTGGTGTTCTGGTTCATCTGTGCCTTCTAGTTGCCA
GCCATCTGTTGTTTGCCCCTCCCCCGTGCCTTCCTTGACCCTGGAAGGTGCCACTCCCA
CTGTCCTTTCCTAATAAAATGAGAAAATTGCATCGCATTGTCTGAGTAGGTGTCATTCTAT
TCTGGGGGGTGGGGTGGGGCAGGACAGCAAGGGGGAGGATTGGGAAGACAATAGCAG
GCATGCTGGGGATGCGGTGGGCTCTATGG

**Supplementary Sequence 2.** Sequence of codon-optimized Bxb1 plasmid

CMV enhancer/promoter: highlighted in grey
SP6 promoter: underlined
Codon-optimized Bxb1: highlighted in green
SV40 poly(A): highlighted in cyan

CGTTACATAACTTACGGTAAATGGCCCGCCTGGCTGACCGCCCAACGACCCCCGCCCA
TTGACGTCAATAATGACGTATGTTCCCATAGTAACGCCAATAGGGACTTTCCATTGACGT
CAATGGGTGGAGTATTTACGGTAAACTGCCCACTTGGCAGTACATCAAGTGTATCATATG
CCAAGTACGCCCCCTATTGACGTCAATGACGGTAAATGGCCCGCCTGGCATTATGCCCA
GTACATGACCTTATGGGACTTTCCTACTTGGCAGTACATCTACGTATTAGTCATCGCTAT
TACCATGGTGATGCGGTTTTGGCAGTACATCAATGGGCGTGGATAGCGGTTTGACTCAC
GGGGATTTCCAAGTCTCCACCCCATTGACGTCAATGGGAGTTTGTTTTGGCACCAAAAT
CAACGGGACTTTCCAAAATGTCGTAACAACTCCGCCCCATTGACGCAAATGGGCGGTAG
GCGTGTACGGTGGGAGGTCTATATAAGCAGAGCTCGTTTAGTGAACCGTCAGATCGCCT
GGAGGCGCCATCCACGCTGTTTTGACCTCCATAGAAGACACCGGGACCGATCCAGCCT
CCGGACTCTAGCCTAGGCTTTTGCAAAAAGCTATTTAGGTGACACTATAGAAGGTACGC
CTGCAGGTACCGGTCCGGAATTCGCCCTTATGATGCGGGCCCTGGTTGTGATTAGACTG
TCCAGAGTGACCGACGCCACCACCTCACCTGAGAGACAGTTGGAGAGCTGCCAGCAGC
TGTGCGCCCAGCGCGGCTGGGACGTGGTGGGCGTGGCCGAGGACCTGGACGTGTCCG
GCGCCGTGGACCCTTTTGACCGGAAGCGGAGACCTAATCTGGCCAGATGGCTGGCCTT
CGAGGAACAGCCTTTCGATGTGATCGTGGCCTACAGAGTGGACCGGCTGACCAGAAGC
ATCCGGCACCTGCAACAGCTGGTGCACTGGGCCGAGGATCACAAAAAGCTGGTGGTCT
CTGCTACAGAGGCCCACTTCGACACAACCACACCTTTCGCCGCTGTGGTGATCGCCCTG
ATGGGGACCGTGGCCCAGATGGAACTCGAGGCCATCAAGGAACGGAACAGATCTGCCG
CCCACTTTAACATCAGAGCCGGCAAGTACAGAGGAAGCCTGCCACCTTGGGGCTACCT
GCCTACCAGAGTTGACGGCGAGTGGAGACTGGTCCCCGACCCCGTGCAGAGAGAGCG
GATCCTGGAGGTGTACCACAGAGTGGTCGACAACCACGAGCCCCTGCATCTGGTGGCT
CATGATCTGAATAGAAGGGGCGTGCTGAGCCCTAAAGACTACTTCGCTCAGCTGCAGG
GCAGAGAACCCCAGGGCAGAGAGTGGAGCGCCACCGCCCTGAAGAGAAGCATGATCA
GCGAGGCCATGCTGGGATACGCCACCCTGAACGGCAAAACCGTGCGGGACGATGATG
GAGCCCCTCTGGTGCGGGCAGAGCCTATCCTAACCCGCGAGCAGCTGGAAGCTCTGAG
AGCCGAGCTGGTCAAGACATCTAGAGCCAAACCTGCTGTGTCCACCCCTAGCCTGCTG
CTGCGGGTGCTGTTCTGTGCCGTGTGCGGTGAACCTGCTTATAAGTTCGCCGGCGGCG
GCAGAAAGCACCCCAGATACCGGTGCAGATCTATGGGCTTCCCCAAGCACTGTGGCAA
CGGCACCGTGGCCATGGCCGAGTGGGATGCTTTTTGCGAGGAACAAGTGCTGGACCTG
CTGGGCGATGCCGAAAGACTAGAGAAGGTGTGGGTGGCTGGCAGCGACAGCGCCGTG
GAACTGGCTGAAGTGAACGCCGAGCTGGTGGACCTCACCAGCCTCATCGGAAGCCCGG
CCTACCGGGCCGGCTCTCCACAGAGAGAAGCCCTGGATGCCCGTATCGCCGCCCTGG
CCGCTAGGCAGGAGGAACTGGAAGGCCTGGAAGCCAGACCAAGCGGCTGGGAGTGGA
GAGAGACAGGACAGCGGTTCGGAGATTGGTGGCGGGAACAGGATACAGCCGCAAAGA
ACACCTGGCTGAGAAGCATGAACGTGCGGCTGACATTCGACGTGAGAGGCGGACTGAC
CAGAACAATCGACTTCGGCGACCTGCAGGAGTACGAGCAACACCTGAGACTGGGCAGC
GTGGTCGAGCGGCTGCACACAGGCATGAGCTAGAAGGGCGAATTCCCGGGTCGACGA
GCTCACTAGCTTGGGATCTTTGTGAAGGAACCTTACTTCTGTGGTGTGACATAATTGGAC
AAACTACCTACAGAGATTTAAAGCTCTAAGGTAAATATAAAATTTTTAAGTGTATAATGTG
TTAAACTAGCTGCATATGCTTGCTGCTTGAGAGTTTTGCTTACTGAGTATGATTTATGAAA
ATATTATACACAGGAGCTAGTGATTCTAATTGTTTGTGTATTTTAGATTCACAGTCCCAAG

GCTCATTTCAGGCCCCTCAGTCCTCACAGTCTGTTCATGATCATAATCAGCCATACCACA
TTTGTAGAGGTTTTACTTGCTTTAAAAAACCTCCCACACCTCCCCCTGAACCTGAAACAT
AAAATGAATGCAATTGTTGTTGTTAACTTGTTTATTGCAGCTTATAATGGTTACAAATAAA
GCAATAGCATCACAAATTTCACAAATAAAGCATTTTTTTCACTGCATTCTAGTTGTGGTTT
GTCCAAACTCATCAATGTATCTTATCATGTCTGGATC

**Supplementary Sequence 3**. Sequence of the lentiviral GFP reporter vector
(the vector bone is pLVX-EF1a-IRES-Puro from Takara Bio inc.).

LTR: in bold
EF1α promoter: underlined
AAVS1 target sequence: highlighted in yellow
Inverted H2B-EGFP coding sequence: highlighted in green
IRES: highlighted in grey
Puromycin resistance gene: highlighted in red

**tggaagggctaattcactcccaaagaagacaagatatccttgatctgtggatctaccacacacaaggctacttccctgat
tagcagaactacacaccagggccaggggtcagatatccactgacctttggatggtgctacaagctagtaccagttgag
ccagataaggtagaagaggccaataaaggagagaacaccagcttgttacaccctgtgagcctgcatgggatggatga
cccggagagagaagtgttagagtggaggtttgacagccgcctagcatttcatcacgtggcccgagagctgcatccgg
agtacttcaagaactgctgatatcgagcttgctacaagggactttccgctggggactttccagggaggcgtggcctggg
cgggactggggagtggcgagccctcagatcctgcatataagcagctgctttttgcctgtactgggtctctctggttagac
cagatctgagcctgggagctctctggctaactagggaacccactgcttaagcctcaataaagcttgccttgagtgcttca
agtagtgtgtgcccgtctgttgtgtgactctggtaactagagatccctcagacccttttagtcagtgtggaaaatctctagc
ag**tggcgcccgaacagggacttgaaagcgaaagggaaaccagaggagctctctcgacgcaggactcggcttgctgaagcgc
gcacggcaagaggcgaggggcggcgactggtgagtacgccaaaaattttgactagcggaggctagaaggagagagatgggt
gcgagagcgtcagtattaagcgggggagaattagatcgcgatgggaaaaaattcggttaaggccagggggaaagaaaaaat
ataaattaaaacatatagtatgggcaagcagggagctagaacgattcgcagttaatcctggcctgttagaaacatcagaaggctg
tagacaaatactgggacagctacaaccatcccttcagacaggatcagaagaacttagatcattatataatacagtagcaaccctct
attgtgtgcatcaaaggatagagataaaagacaccaaggaagctttagacaagatagaggaagagcaaaacaaaagtaaga
ccaccgcacagcaagcggccggccgctgatcttcagacctggaggaggagatatgagggacaattggagaagtgaattatata
aatataaagtagtaaaaattgaaccattaggagtagcacccaccaaggcaaagagaagagtggtgcagagagaaaaaaga
gcagtgggaataggagctttgttccttgggttcttgggagcagcaggaagcactatgggcgcagcgtcaatgacgctgacggtac
aggccagacaattattgtctggtatagtgcagcagcagaacaatttgctgagggctattgaggcgcaacagcatctgttgcaactc
acagtctggggcatcaagcagctccaggcaagaatcctggctgtggaaagatacctaaaggatcaacagctcctggggatttgg
ggttgctctggaaaactcatttgcaccactgctgtgccttggaatgctagttggagtaataaatctctggaacagatttggaatcacac
gacctggatggagtgggacagagaaattaacaattacacaagcttaatacactccttaattgaagaatcgcaaaaccagcaaga
aaagaatgaacaagaattattggaattagataaatgggcaagtttgtggaattggtttaacataacaaattggctgtggtatataaa
attattcataatgatagtaggaggcttggtaggtttaagaatagtttttgctgtactttctatagtgaatagagttaggcagggatattcac
cattatcgtttcagacccacctcccaaccccgaggggacccgacaggcccgaaggaatagaagaagaaggtggagagagag
acagagacagatccattcgattagtgaacggatctcgacggtatcgcctttaaaagaaaaggggggattggggggtacagtgca
ggggaaagaatagtagacataatagcaacagacatacaaactaaagaattacaaaaacaaattacaaaaattcaaaattttcg
ggtttattacagggacagcagagatccagtttatcgatgagtaattcatacaaaaggactcgcccctgccttggggaatcccaggg
accgtcgttaaactcccactaacgtagaacccagagatcgctgcgttcccgcccctcacccgcccgctctcgtcatcactgaggt
ggagaagagcatgcgtgaggctccggtgcccgtcagtgggcagagcgcacatcgcccacagtccccgagaagttggggga
ggggtcggcaattgaaccggtgcctagagaaggtggcgcggggtaaactgggaaagtgatgtcgtgtactggctccgcctttttcc
cgagggtgggggagaaccgtatataagtgcagtagtcgccgtgaacgttctttttcgcaacgggtttgccgccagaacacaggta
agtgccgtgtgtggttcccgcgggcctggcctctttacgggttatggcccttgcgtgccttgaattacttccacgcccctggctgcagta
cgtgattcttgatcccgagcttcgggttggaagtgggtgggagagttcgaggccttgcgcttaaggagccccttcgcctcgtgcttga
gttgaggcctggcttgggcgctggggccgccgcgtgcgaatctggtggcaccttcgcgcctgtctcgctgctttcgataagtctctag
ccatttaaaatttttgatgacctgctgcgacgctttttttctggcaagatagtcttgtaaatgcgggccaagatctgcacactggtatttcg
gtttttggggccgcgggcggcgacggggcccgtgcgtcccagcgcacatgttcggcgaggcggggcctgcgagcgcggccac
cgagaatcggacgggggtagtctcaagctggccggcctgctctggtgcctggcctcgcgccgccgtgtatcgccccgccctgggc
ggcaaggctggcccggtcggcaccagttgcgtgagcggaaagatggccgcttcccggccctgctgcagggagctcaaaatgg
aggacgcggcgctcgggagagcgggcgggtgagtcacccacacaaaggaaaagggcctttccgtcctcagccgtcgcttcat

gtgactccacggagtaccgggcgccgtccaggcacctcgattagttctcgagcttttggagtacgtcgtctttaggttgggggagg
ggttttatgcgatggagtttccccacactgagtgggtggagactgaagttaggccagcttggcacttgatgtaattctccttggaatttg
cccttttgagtttggatcttggttcattctcaagcctcagacagtggttcaaagtttttttcttccatttcaggtgtcgtgaggatctatttccg
gtgaattcAACGGGGCTCAGTCTGAAGAGCAGAGCCAGGAACCCCTGTAGGGAAGGGGCA
GGAGAGCCAGGGGCATGAGATGGTGGACGAGGAAGGGGGACAGGGAAGCCTGAGCG
CCTCTCCTGGGCTTGCCAAGGACTCAAACCCAGAAGCCCAGATTACTTGTACAGCTCGT
CCATGCCGAGAGTGATCCCGGCGGCGGTCACGAACTCCAGCAGGACCATGTGATCGCG
CTTCTCGTTGGGGTCTTTGCTCAGGGCGGACTGGGTGCTCAGGTAGTGGTTGTCGGGC
AGCAGCACGGGGCCGTCGCCGATGGGGGTGTTCTGCTGGTAGTGGTCGGCGAGCTGC
ACGCTGCCGTCCTCGATGTTGTGGCGGATCTTGAAGTTCACCTTGATGCCGTTCTTCTG
CTTGTCGGCCATGATATAGACGTTGTGGCTGTTGTAGTTGTACTCCAGCTTGTGCCCCA
GGATGTTGCCGTCCTCCTTGAAGTCGATGCCCTTCAGCTCGATGCGGTTCACCAGGGTG
TCGCCCTCGAACTTCACCTCGGCGCGGGTCTTGTAGTTGCCGTCGTCCTTGAAGAAGAT
GGTGCGCTCCTGGACGTAGCCTTCGGGCATGGCGGACTTGAAGAAGTCGTGCTGCTTC
ATGTGGTCGGGGTAGCGGCTGAAGCACTGCACGCCGTAGGTCAGGGTGGTCACGAGG
GTGGGCCAGGGCACGGGCAGCTTGCCGGTGGTGCAGATGAACTTCAGGGTCAGCTTG
CCGTAGGTGGCATCGCCCTCGCCCTCGCCGGACACGCTGAACTTGTGGCCGTTTACGT
CGCCGTCCAGCTCGACCAGGATGGGCACCACCCCGGTGAACAGCTCCTCGCCCTTGCT
CACCATGGTGGCGACCGGTGGgTCtTTAGCGCTGGTGTACTTGGTGATGGCCTTAGTAC
CCTCGGACACGGCGTGCTTGGCCAACTCCCCAGGCAGCAGCAGGCGCACGGCCGTCT
GGATCTCCCTGGAGGTGATGGTCGAGCGCTTGTTGTAATGCGCCAGGCGGGAAGCCTC
ACCTGCGATGCGCTCGAAAATGTCGTTCACAAAactATTCATGATGCCCATGGCCTTGGA
CGAAATGCCGGTGTCAGGGTGGACCTGCTTCAGAACCTTGTACACATAGATGGAATAGC
TCTCCTTGCGGCTGCGCTTGCGCTTCTTGCCGCCTTTCTTCTGCGCCTTAGTCACCGCC
TTCTTGGAGCCCTTTTTCGGGGCGGGAGCAGACTTCGCTGGCTCTGGCATggtggcGGCC
GCCGGGAACTGCCGCTGGCCCCCCACCGCCCCAAGGATCTCCCGGTCCCCGCCCGGC
GTGCTGACGTCACGGCGCTGCCCCAGGGTGTGCTGGGCAGGTCGCGGGGAGCGCTGG
GAAATGGAGTCCATTAGCAGAAGTGGCCCTTGGCCACTTCCAGGAGTCGCTGTGCCCC
GATGCACACTGGGAAGTCCGCAGCggatcccgcccctctccctcccccccccctaacgttactggccgaagccg
cttggaataaggccggtgtgcgtttgtctatatgttattttccaccatattgccgtcttttggcaatgtgagggcccggaaacctggccct
gtcttcttgacgagcattcctaggggtctttcccctctcgccaaaggaatgcaaggtctgttgaatgtcgtgaaggaagcagttcctct
ggaagcttcttgaagacaaacaacgtctgtagcgaccctttgcaggcagcggaaccccccacctggcgacaggtgcctctgcgg
ccaaaagccacgtgtataagatacacctgcaaaggcggcacaaccccagtgccacgttgtgagttggatagttgtggaaagagt
caaatggctctcctcaagcgtattcaacaaggggctgaaggatgcccagaaggtaccccattgtatgggatctgatctggggcctc
ggtgcacatgctttacatgtgtttagtcgaggttaaaaaaacgtctaggccccccgaaccacggggacgtggttttcctttgaaaaa
cacgatgataagcttgccacaacccacaaggagacgaccttccatgaccgagtacaagcccacggtgcgcctcgccacccgc
gacgacgtccccgggccgtacgcaccctcgccgccgcgttcgccgactaccccgccacgcgccacaccgtcgacccggacc
gccacatcgagcgggtcaccgagctgcaagaactcttcctcacgcgcgtcgggctcgacatcggcaaggtgtgggtcgcggac
gacggcgccgcggtggcggtctggaccacgccggagagcgtcgaagcgggggcggtgttcgccgagatcggcccgcgcatg
gccgagttgagcggttcccggctggccgcgcagcaacagatggaaggcctcctggcgccgcaccggcccaaggagcccgcg
tggttcctggccaccgtcggcgtctcgcccgaccaccagggcaagggtctgggcagcgccgtcgtgctccccggagtggaggc
ggccgagcgcgccggggtgcccgccttcctggagacctccgcgccccgcaacctccccttctacgagcggctcggcttcaccgt
caccgccgacgtcgaggtgcccgaaggaccgcgcacctggtgcatgacccgcaagcccggtgcctagacgcgtctggaaca
atcaacctctggattacaaaatttgtgaaagattgactggtattcttaactatgttgctccttttacgctatgtggatacgctgctttaatgc
ctttgtatcatgctattgcttcccgtatggctttcattttctcctccttgtataaatcctggttgctgtctctttatgaggagttgtggcccgttgtc
aggcaacgtggcgtggtgtgcactgtgtttgctgacgcaacccccactggttggggcattgccaccacctgtcagctcctttccggg
actttcgctttccccctccctattgccacggcggaactcatcgccgcctgccttgcccgctgctggacaggggctcggctgttgggca
ctgacaattccgtggtgttgtcggggaagctgacgtcctttccatggctgctcgcctgtgttgccacctggattctgcgcgggacgtcct
tctgctacgtcccttcggccctcaatccagcggaccttccttcccgcggcctgctgccggctctgcggcctcttccgcgtcttcgccttc

gccctcagacgagtcggatctccctttgggccgcctccccgcctggaattaattctgcagtcgagacctagaaaaacatggagca
atcacaagtagcaatacagcagctaccaatgctgattgtgcctggctagaagcacaagaggaggaggaggtgggttttccagtc
acacctcaggtacctttaagaccaatgacttacaaggcagctgtagatcttagccactttttaaaagaaaagaggggga**ctggaag**
**ggctaattcactcccaacgaagacaagatatccttgatctgtggatctaccacacacaaggctacttccctgattagcag**
**aactacacaccagggccaggggtcagatatccactgacctttggatggtgctacaagctagtaccagttgagccagat**
**aaggtagaagaggccaataaaggagagaacaccagcttgttacaccctgtgagcctgcatgggatggatgacccgg**
**agagagaagtgttagagtggaggtttgacagccgcctagcatttcatcacgtggcccgagagctgcatccggagtactt**
**caagaactgctgatatcgagcttgctacaagggactttccgctggggactttccagggaggcgtggcctgggcgggac**
**tggggagtggcgagccctcagatcctgcatataagcagctgctttttgcctgtactgggtctctctggttagaccagatct**
**gagcctgggagctctctggctaactagggaacccactgcttaagcctcaataaagcttgccttgagtgcttcaagtagtg**
**tgtgcccgtctgttgtgtgactctggtaactagagatccctcagacccttttagtcagtgtggaaaatctctagcagt**agtag
ttcatgtcatcttattattcagtatttataacttgcaaagaaatgaatatcagagagtgagaggccttgacattgctagcgtttaccgtcg
acctctagctagagcttggcgtaatcatggtcatagctgtttcctgtgtgaaattgttatccgctcacaattccacacaacatacgagc
cggaagcataaagtgtaaagcctggggtgcctaatgagtgagctaactcacattaattgcgttgcgctcactgcccgctttccagtc
gggaaacctgtcgtgccagctgcattaatgaatcggccaacgcgcggggagaggcggtttgcgtattgggcgctcttccgcttcct
cgctcactgactcgctgcgctcggtcgttcggctgcggcgagcggtatcagctcactcaaaggcggtaatacggttatccacagaa
tcaggggataacgcaggaaagaacatgtgagcaaaaggccagcaaaaggccaggaaccgtaaaaaggccgcgttgctggc
gtttttccataggctccgcccccctgacgagcatcacaaaaatcgacgctcaagtcagaggtggcgaaacccgacaggactata
aagataccaggcgtttccccctggaagctccctcgtgcgctctcctgttccgaccctgccgcttaccggatacctgtccgcctttctcc
cttcgggaagcgtggcgctttctcatagctcacgctgtaggtatctcagttcggtgtaggtcgttcgctccaagctgggctgtgtgcac
gaaccccccgttcagcccgaccgctgcgccttatccggtaactatcgtcttgagtccaacccggtaagacacgacttatcgccact
ggcagcagccactggtaacaggattagcagagcgaggtatgtaggcggtgctacagagttcttgaagtggtggcctaactacgg
ctacactagaagaacagtatttggtatctgcgctctgctgaagccagttaccttcggaaaaagagttggtagctcttgatccggcaa
acaaaccaccgctggtagcggtggtttttttgtttgcaagcagcagattacgcgcagaaaaaaaggatctcaagaagatcctttgat
cttttctacggggtctgacgctcagtggaacgaaaactcacgttaagggattttggtcatgagattatcaaaaaggatcttcacctag
atccttttaaattaaaaatgaagttttaaatcaatctaaagtatatatgagtaaacttggtctgacagttaccaatgcttaatcagtgagg
cacctatctcagcgatctgtctatttcgttcatccatagttgcctgactccccgtcgtgtagataactacgatacgggagggcttaccat
ctggccccagtgctgcaatgataccgcgagacccacgctcaccggctccagatttatcagcaataaaccagccagccggaagg
gccgagcgcagaagtggtcctgcaactttatccgcctccatccagtctattaattgttgccgggaagctagagtaagtagttcgcca
gttaatagtttgcgcaacgttgttgccattgctacaggcatcgtggtgtcacgctcgtcgtttggtatggcttcattcagctccggttccca
acgatcaaggcgagttacatgatcccccatgttgtgcaaaaaagcggttagctccttcggtcctccgatcgttgtcagaagtaagttg
gccgcagtgttatcactcatggttatggcagcactgcataattctcttactgtcatgccatccgtaagatgcttttctgtgactggtgagta
ctcaaccaagtcattctgagaatagtgtatgcggcgaccgagttgctcttgcccggcgtcaatacgggataataccgcgccacata
gcagaactttaaaagtgctcatcattggaaaacgttcttcggggcgaaaactctcaaggatcttaccgctgttgagatccagttcgat
gtaacccactcgtgcacccaactgatcttcagcatcttttactttcaccagcgtttctgggtgagcaaaaacaggaaggcaaaatgc
cgcaaaaaagggaataagggcgacacggaaatgttgaatactcatactcttcctttttcaatattattgaagcatttatcagggttatt
gtctcatgagcggatacatatttgaatgtatttagaaaaataaacaaataggggttccgcgcacatttccccgaaaagtgccacctg
acgtcgacggatcgggagatcaacttgtttattgcagcttataatggttacaaataaagcaatagcatcacaaatttcacaaataaa
gcatttttttcactgcattctagttgtggtttgtccaaactcatcaatgtatcttatcatgtctggatcaactggataactcaagctaaccaa
aatcatcccaaacttcccaccccataccctattaccactgccaattacctgtggtttcatttactctaaacctgtgattcctctgaattattt
tcattttaaagaaattgtatttgttaaatatgtactacaaacttagtagt
